# A Metabolite-Based Resistance Mechanism Against Malaria

**DOI:** 10.1101/2025.02.20.639306

**Authors:** Ana Figueiredo, Sonia Trikha Rastogi, Susana Ramos, Fátima Nogueira, Katherine De Villiers, António G. Gonçalves de Sousa, Lasse Votborg-Novél, Cäcilie von Wedel, Pinkus Tober-Lau, Elisa Jentho, Sara Pagnotta, Miguel Mesquita, Silvia Cardoso, Giulia Bortolussi, Andrés F. Muro, Erin M. Tranfield, Jessica Thibaud, Denise Duarte, Ana Laura Sousa, Sandra N. Pinto, Jamil Kitoko, Ghyslain Mombo-Ngoma, Johannes Mischlinger, Sini Junttila, Marta Alenquer, Maria João Amorim, Chirag Vasavda, Piter J. Bosma, Sara Violante, Bernhard Drotleff, Tiago Paixão, Silvia Portugal, Florian Kurth, Laura L. Elo, Bindu D. Paul, Rui Martins, Miguel P. Soares

## Abstract

Whether jaundice, a common presentation of *Plasmodium* (*P*.) *falciparum* malaria (1-3) arising from the accumulation of circulating bilirubin, represents an adaptive or maladaptive response to *Plasmodium* spp. infection is not understood (1-3). We found that asymptomatic *P. falciparum* infection was associated with a >10-fold higher ratio of unconjugated bilirubin over parasite burden, compared to symptomatic malaria. Genetic suppression of bilirubin synthesis by biliverdin reductase A (BVRA) (4) increased parasite virulence and malaria mortality in mice. Accumulation of unconjugated bilirubin in plasma, via genetic inhibition of hepatic conjugation by UDP glucuronosyltransferase family 1 member A1 (UGT1A1) (*5*), was protective against malaria in mice. Unconjugated bilirubin inhibited *P. falciparum* proliferation in red blood cells (RBC) via a mechanism that suppressed mitochondrial pyrimidine synthesis. Moreover, unconjugated bilirubin inhibited hemozoin (Hz) crystallization and compromised the parasite’s food vacuole. In conclusion, jaundice represents a metabolic response to *Plasmodium spp*. infection that limits malaria severity.

---

Parasites from the *Plasmodium* genus proliferate in the red blood cell (RBC) compartment of their numerous vertebrate hosts leading intravascular hemolysis and release of hemoglobin into plasma (*6, 7*). Auto-oxidation of extracellular hemoglobin precipitates the detachment of its non-covalently bound prosthetic heme groups (*6, 8-11*). This produces labile heme (*12*), an independent risk factor of *P. falciparum* malaria severity (*6*) that plays a central stage in the pathogenesis of severe malaria in mice (*8, 10, 13*).

The pathogenic effects of labile heme are countered via the induction of heme catabolism by heme oxygenase-1 (HO-1/*HMOX1*) in the infected host (*8, 9, 14*). Heme catabolism produces equimolar amounts of iron, carbon monoxide and biliverdin (*15*) and is coupled to the reduction of biliverdin into bilirubin, catalyzed by biliverdin reductase A (BVRA) (*4*) (*Fig. S1A*). This reaction produces lipophilic unconjugated bilirubin (*i*.*e*., indirect) that circulates in plasma bound to albumin (*16*) (*Fig. S1A*). Bilirubin is conjugated to glucuronic acid in hepatocytes by UDP glucuronosyltransferase family 1 member A1 (UGT1A1) (*17*) (*Fig. S1A*), producing water soluble conjugated bilirubin (*i*.*e*., direct) that is excreted via the bile or urine (*18*) (*Fig. S1A*).

Severe presentations of *P. falciparum* malaria are often associated with the accumulation of bilirubin in plasma, a condition referred to as jaundice of malaria, when associated with visible yellowing of the skin or white of the eyes (1-3). Jaundice of malaria develops once the rate of bilirubin production by BVRA exceeds that of bilirubin conjugation by UGT1A1 (*19*). Here we asked whether the accumulation of circulating bilirubin during *Plasmodium* spp. infection represents an adaptive or maladaptive metabolic response to malaria.

## Onset of *P. falciparum* malaria disease symptoms is associated with a reduction in the ratio of circulating unconjugated (indirect) bilirubin over parasite burden

A retrospective analysis of a cohort of patients with *P. falciparum* infection and asymptomatic or symptomatic malaria showed a positive correlation between the presence of clinical symptoms and parasite burden (*i*.*e*., number of parasites *per* µL blood) (*Fig. 1A*), total circulating bilirubin (*Fig. 1B*), direct bilirubin (*Fig. 1C*) and indirect bilirubin (*Fig. 1D*). While these observations are consistent with previous studies (*1-3*), the standard analytical methods used in clinical practice fail to provide an accurate quantification of unconjugated bilirubin in plasma (*16, 18, 20*). This is due to the binding of unconjugated bilirubin to albumin in plasma (*16*), which hinders the colorimetric reaction used in these assays (*16, 18, 20*). Moreover, byproducts of hemolysis, such as labile heme, can interfere with these colorimetric assays, adding to their inaccuracy in the context of *Plasmodium* infection (*6, 11*).

**Figure 1.**
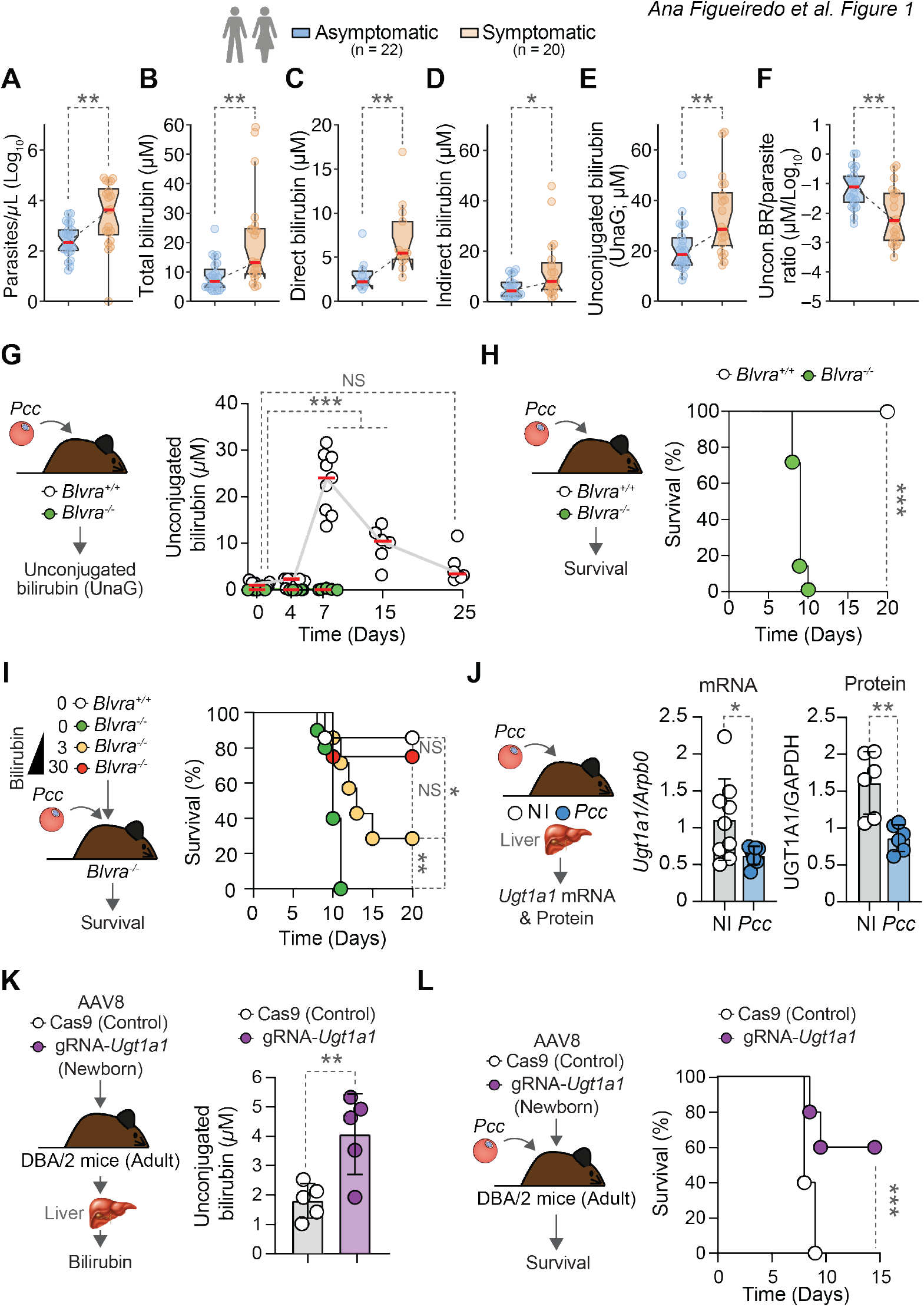
Unconjugated bilirubin confers resistance to malaria. (**A**) Parasite burden (*P. falciparum* iRBC/µL of blood) and concentrations of (**B**) total (i.e., direct plus indirect), (**C**) direct and (**D**) indirect bilirubin (measured using the Roche Diazo method) in plasma from *P. falciparum* infected individuals, stratified according to disease severity as: asymptomatic and symptomatic malaria. (**E**) Plasma concentration of unconjugated bilirubin measured by the UnaG-based assay in same patients as (A-D). **(F)** Ratio of unconjugated bilirubin measured by the UnaG-based assay (E) to parasite burden (A). Data (A-F) is represented as box plots; red lines correspond to median values and error bars correspond to the interquartile range (IQR). (**G**) Concentration of unconjugated bilirubin in plasma from *Blvra*^*+/+*^ and *Blvra*^*-/-*^ mice (n=6-9 *per* genotype), before (Day 0) and after *Pcc* infection (Days 4, 7, 15 and 25) measured by the UnaG-based assay. Data represented as mean ± SD, pooled from two independent experiments with similar trend. (**H**) Survival of *Pcc*-infected *Blvra*^*-/-*^ and control *Blvra*^+/+^ mice. Data from n=7 *per* genotype, pooled from two independent experiments with similar trend. (**I**) Survival of *Pcc*-infected *Blvra*^*-/-*^ mice receiving bilirubin (30 or 3 mg/Kg; daily; *i*.*p*.) or vehicle. Data from n=4-8 *per* treatment, pooled from two independent experiments with similar trend. (**J**) Quantification of *Ugt1a1* mRNA (qPCR; *left panel*) (n=7-9 *per* genotype) and protein (Western Blot; *right panel*) (n=6 *per* genotype) expression in the liver at day 7 post-*Pcc* infection (*Pcc*) or non-infected (NI) C57BL/6J mice. Data represented as mean ± SD, pooled from two independent experiments with similar trend. (**K**) Concentration of unconjugated bilirubin in plasma of adult DBA/2 mice transduced 2-4 days after birth with AAV8-gRNA-*Ugt1a1* repressing hepatic *Ugt1a1* or control AAV8-Cas9. Data from n=5 mice *per* group, represented as mean ± SD, from one experiment representative of three with similar trend. (**L**) Survival of the mice from (K) infected with *Pcc*. Circles in (A-F) correspond to *P. falciparum*-infected patients and in (G-L) to individual mice. *P* values determined using: (A-F) Mann-Whitney U test, (G) Two-Way ANOVA with Bonferroni’s multiple comparison test (for genotypes) and Ordinary One-Way ANOVA with Tukey’s multiple comparison (for days post-infection), (H, I, L) Log-rank (Mantel-Cox) test, (J *right panel*, K) t test, and (J *left panel*) using Mann-Whitney U test. NS: not significant; \**p*<0.05; \*\**p*<0.01; \*\*\**p*<0.001. NS: not significant, *p*>0.05.

To provide a precise concentration of unconjugated bilirubin in plasma we used a highly specific unconjugated bilirubin-inducible fluorescent protein (UnaG)-based assay (*Fig. S1B*) (*21, 22*). We found that the concentration of unconjugated bilirubin in plasma from asymptomatic *P. falciparum*-infected individuals ranged from 8 to 50 µM (*Fig. 1E*), that is, 3.75-fold higher than estimated using standard analytical methods (*Fig. S1C*). The concentration of unconjugated bilirubin in plasma from symptomatic *P. falciparum* malaria patients ranged from 14 to 67 µM (*Fig. 1E*), 2.67-fold higher than estimated using standard analytical methods (*Fig. S1C*). This confirms that standard analytical methods used in clinical practice underestimate the concentration of unconjugated bilirubin in the plasma from *P. falciparum* malaria patients (*16, 18, 20*).

Using the same UnaG-based assay, we found that asymptomatic *P. falciparum* infection was associated with a >10-fold higher ratio of circulating unconjugated bilirubin over parasite burden compared to symptomatic malaria (*Fig 1F*). As unconjugated bilirubin inhibits *P. falciparum* proliferation *in vitro* (*23*), we hypothesized that the accumulation of unconjugated bilirubin in plasma represents an adaptive response to *P. falciparum* malaria.

### Bilirubin production by BVRA is essential to survive experimental malaria

To test functionally the anti-malarial effect of unconjugated bilirubin, we infected C57BL/6J mice with *P. chabaudi chabaudi* AS (*Pcc*), a well-established non-lethal experimental model of malaria (*24*). Using the UnaG-based assay (*Fig. S1B*) (*21, 22*), we found that the concentration of circulating unconjugated bilirubin increased abruptly from day 4 to 7 post-infection to reach 13-31 µM (*Fig. 1G*), in the range of *P. falciparum* malaria patients (*Fig. 1E*). The specificity of the UnaG-based assay was confirmed using *Blvra*-deficient (*Blvra*^*-/-*^) mice (*22*), which did not express *Blvra* mRNA (*Fig. S2A*) and BVRA protein (*Fig. S2B,C*) and did not produce bilirubin (*Fig. 1G*).

As compared to the non-lethal outcome of *Pcc* infection in C57BL/6J *Blvra*^+/+^ mice, all littermate C57BL/6J *Blvra*^*-/-*^ mice succumbed to the infection within 7-10 days (*Fig. 1H*). This suggest that the production of bilirubin by BVRA confers protection against *Plasmodium* infection.

Using a biliverdin-inducible infrared fluorescent protein (iRFP)-based assay (*Fig. S2D*) (*22, 25*) to quantify the concentration of biliverdin in plasma, we found that the *Pcc*-infected *Blvra*^*-/-*^ mice (*Fig. 1H*) accumulated relatively low levels of circulating biliverdin, in the range of 2.5-6 µM (*Fig. S2E*). This suggests that the lethal outcome of *Pcc* infection in *Blvra*^*-/-*^ mice is not due to a putative accumulation of pathogenic levels of circulating biliverdin.

Pharmacologic administration of unconjugated bilirubin protected *Blvra*^*-/-*^ mice from succumbing to *Pcc* infection (*Fig. 1I*). The protective effect of bilirubin was dose-dependent, that is, higher bilirubin dosage restored the survival of *Pcc*-infected *Blvra*^*-/-*^ mice to the same extent as *Pcc*-infected *Blvra*^+/+^ mice (*Fig. 1I*). This suggests that the lethal outcome of *Pcc* infection in *Blvra*^*-/-*^ mice is attributed to lack of bilirubin production, implying that the protective effect of BVRA against malaria is mediated by bilirubin.

### Bilirubin is not essential to survive non-hemolytic infections

We asked whether the production of bilirubin by BVRA is also protective against bacterial or viral infectious diseases. Disease severity and pathogen burden were indistinguishable in *Blvra*^*-/-*^ and control *Blvra*^+/+^ mice subjected to bacterial sepsis (*Fig. S3A-C*) or to influenza A virus infection (*Fig. S3D-F*). This suggests that the protective effect of unconjugated bilirubin is specific to hemolytic conditions, such as malaria.

### Inhibition of hepatic UGT1A1 is protective against experimental malaria

The relative levels of *Hmox-1* mRNA expression were highly induced in different organs from *Pcc*-infected *vs*. non-infected C57BL/6J mice (*Fig. S4A*), consistent with previously described (*14*). In contrast, the relative levels of *Blvra* mRNA expression were not induced in *Pcc*-infected *vs*. non-infected C57BL/6J mice (*Fig. S4B*). This suggests that the induction of bilirubin production in response to *Plasmodium* infection is dependent on the induction of HO-1 to increase the amount of biliverdin that can be reduced into bilirubin by BVRA (*Fig. S1A*).

*Pcc* infection was associated with a marked decrease in the relative level of hepatic *Ugt1a1* mRNA (*Fig. 1J, S4C*) and UGT1A1 protein (*Fig. 1J, S4D*) expression, 7 days post-infection, compared to non-infected controls. This suggests that the accumulation of circulating bilirubin in response to *Plasmodium* infection is sustained via the repression of hepatic UGT1A1 (*Fig. S1A*).

To test whether inhibition of hepatic bilirubin conjugation is protective against malaria we used *Pcc* infection in DBA/2 mice, as a lethal experimental model of malaria (*26*). Hepatic UGT1A1 was repressed specifically in the liver of newborn DBA/2 mice, transduced with a recombinant adeno-associated virus serotype 8 (AAV8) encoding the *Staphylococcus aureus* (Sa) CRISPR associated protein 9 (Cas9) and a single guide RNA (gRNA) targeting *Ugt1a1* (AAV8-gRNA-*Ugt1a1*), as previously described (*27*). Adult DBA/2 mice transduced with AAV8-gRNA-*Ugt1a1* presented lower levels of hepatic UGT1A1 protein, compared to controls transduced with a AAV8 encoding SaCas9 without the targeting gRNA (AAV8-Cas9) (*Fig. S4E,F*). This was associated with an increase in the concentration of unconjugated bilirubin in plasma (*Fig. 1K*) and with a major survival advantage against *Pcc* infection, compared to control *Pcc*-infected DBA/2 mice transduced with AAV8-Cas9 (*Fig. 1L*). These observations suggest that the repression of hepatic UGT1A1 in response to *Plasmodium* infection is protective against malaria.

### Bilirubin reduces *Plasmodium* burden in experimental malaria

*P. falciparum* proliferation is inhibited by unconjugated bilirubin *in vitro* (*23*), suggesting that the accumulation of unconjugated bilirubin in plasma confers resistance to malaria (*i*.*e*., reduces the host parasite burden). In strong support of this hypothesis, *Pcc*-infected *Blvra*^*-/-*^ mice presented a clear increase in the percentage of circulating infected-RBC (iRBC; parasitemia) 7-9 days post-infection (*Fig. 2A*), accounting for 10-fold increase in the number of iRBC (*i*.*e*., parasite burden), compared to *Pcc*-infected *Blvra*^+/+^ mice (*Fig. 2B*). Consistent with this observation, the administration of unconjugated bilirubin to *Pcc*-infected *Blvra*^*-/-*^ mice reduced parasitemia (*Fig. S5A*) and parasite burden, compared to vehicle-treated *Pcc*-infected *Blvra*^*-/-*^ mice (*Fig. S5B*). Moreover, repression of bilirubin conjugation also reduced the parasitemia (*Fig. S5C*) and parasite burden (*Fig. S5D*) of *Pcc*-infected DBA/2 mice transduced with a AAV8-gRNA-*Ugt1a1 vs*. controls transduced with a AAV8-Cas9. These observations suggest that the accumulation of unconjugated bilirubin in plasma confers resistance to *Plasmodium* infection.

**Figure 2.**
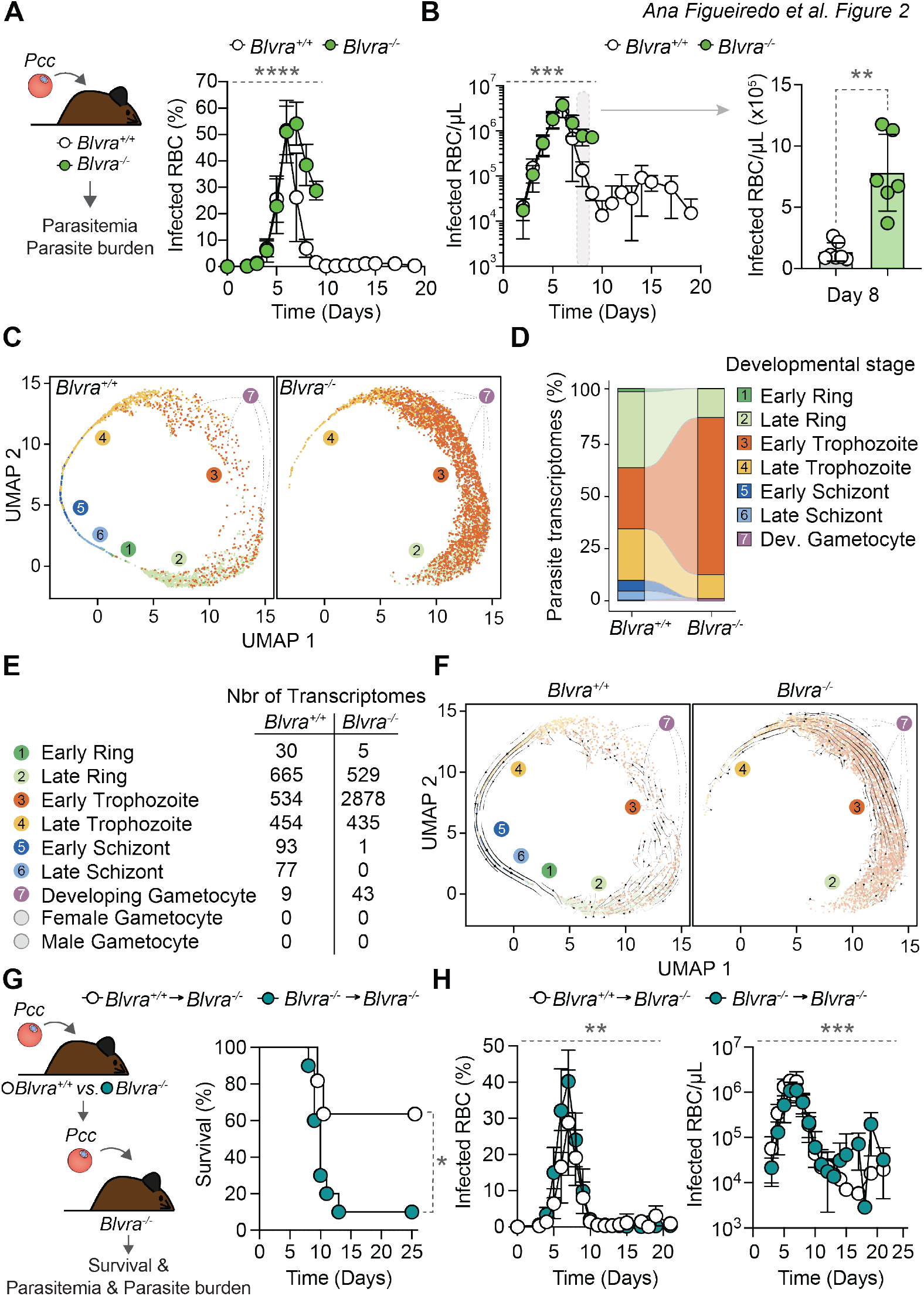
Unconjugated bilirubin regulates *Plasmodium* blood stage development and virulence. (**A**) Percentage of iRBC and (**B**) parasite burden (iRBC/µL) in *Pcc*-infected *Blvra*^*-/-*^ and control *Blvra*^+/+^ mice. Data from n=7 *per* genotype, pooled from two independent experiments with similar trend. Same mice as (Figure 1H). *Right panel* highlights Day 8 from *left panel*. (**C**) UMAP projection of single parasite transcriptomes, of FACS-sorted circulating iRBC from *Blvra*^*+/+*^ (n= 1862; *left panel*) and *Blvra*^*-/-*^ (n= 3891; *right panel*) mice, 7 days after *Pcc* infection. Colors and numbers identify different parasite developmental stages based on scmap projection to a *P. falciparum* atlas (*See Tables S1-4*). (**D**) Composition and (**E**) absolute number of parasite developmental stages. (**F**) UMAP projection as in (C) with arrows representing the relative change in transcriptional state based on RNA velocity analysis. (**G**) Survival of *Blvra*^*-/-*^ mice infected with *Pcc* iRBC, isolated from *Blvra*^*-/-*^ *vs. Blvra*^*+/+*^ mice (n=10-11 *per* genotype). (**H**) Percentage of iRBC (*left panel*), and parasite burden (iRBC/µL, *right panel*) of the same mice as (G). Data represented as mean ± SD, pooled from three independent experiments with similar trend. Circles in (B, *right panel*) represent individual mice. *P* values determined using: (A,B *left panel*, H) Two-Way ANOVA, (B *right panel*) Mann Whitney U test, and (G) Log-rank (Mantel-Cox) test. \**p*<0.05; \*\**p*<0.01; \*\*\**p*<0.001; \*\*\*\**p*<0.0001.

### Bilirubin targets *Plasmodium* inside the RBC

Unconjugated bilirubin diffuses across cellular membranes (*28, 29*), suggesting that unconjugated bilirubin can target *Plasmodium* inside the RBC. To test this hypothesis, we performed single-cell RNA sequencing (scRNAseq) of circulating *Pcc*-infected RBC (iRBC) FACS-sorted from *Blvra*^*-/-*^ *vs. Blvra*^+/+^ mice. Single parasite transcriptomes were assigned to specific developmental stages, based on a single-cell malaria atlas (*30*). Visualization, using uniform manifold approximation and projection (UMAP), revealed a circular orientation of parasites along the asexual cycle: Rings (Populations 1, 2), Trophozoites (Populations 3, 4), Schizonts (Populations 5, 6) and developing Gametocytes (population 7) (*Fig. S6A,B; Table S1*), consistent with previous descriptions (*30-32*).

Circulating iRBC from *Blvra*^*-/-*^ mice exhibited a marked increase in the proportion (*Fig. 2C,D*) and relative number (*Fig. 2E*) of parasite transcriptomes corresponding to metabolically-active early trophozoites (Population 3). This was associated with loss of early (Population 5) and late (Population 6) schizonts as well as early rings (Population 1), and with a decrease in late rings (Population 2) and late trophozoites (Population 4) (*Fig. 2C,D*), as confirmed by morphological analyzes (*Fig. S7A-C*). These changes were not attributed to modulation of parasite sequestration in different organs, as determined by shifting the “light cycle” of infection (*Fig. S7D*).

Progression of *Pcc* infection in *Blvra*^+/+^ mice was associated with the expected parasite developmental trajectory whereby late trophozoites (Population 4) developed into early (Population 5) and late (Population 6) schizonts to give rise to early (Population 1) ring stages, as assessed by RNA velocity analyses (*33*) (*Fig. 2F*). Progression through late rings (Population 2) and early trophozoites (Population 3) was diffuse in *Blvra*^+/+^ mice and contrasted sharply to the progression of the same parasite developmental stages in *Blvra*^*-/-*^ mice (*Fig. 2F*). This suggests that bilirubin reduces the fitness of early trophozoites from *Blvra*^+/+^ *vs. Blvra*^*-/-*^ mice.

Late ring stages (Population 2) from *Blvra*^*-/-*^ mice showed a marked increase in gene expression profiles associated with peptidase activity, heme binding, electron transport chain, proteosome activity and catabolic processes, compared to ring stages from control *Blvra*^+/+^ mice (*Fig. S8, S9A; Tables S2,3*). Early trophozoites (Population 3) in *Blvra*^*-/-*^ mice also showed a clear increase in gene expression associated with peptidase activity, ATP-dependent protein folding chaperone and unfolded protein binding, proteosome core complex, ubiquitin and catabolic processes, compared to early trophozoites from control *Blvra*^+/+^ mice (*Fig. S8, S9B; Tables S2,3*). In addition, there was a concomitant decrease in gene expression profiles associated with chromatin structure (*Fig. S8, S9B; Tables S2,3*). Late trophozoites (Population 4) in *Blvra*^*-/-*^ mice showed a marked increase in gene expression profiles associated with ATP-dependent protein folding, RNA binding and translational activity, glycolytic, pyruvate and carbohydrate, ADP, and ATP metabolic processes (*Fig. S8, S9C; Tables S2,3*). Moreover, there was a concomitant decrease in gene expression profiles associated with structural constituents of chromatin, DNA replication and chromosome organization/segregation (*Fig. S8, S9C; Tables S2,3*).

### Bilirubin production by BVRA reduces parasite virulence

Among the 1019 differentially expressed genes in *Pcc* late rings and early and late trophozoites from *Blvra*^*-/-*^ *vs. Blvra*^+/+^ mice (*Fig. S8*), 118 (11.6%) (*Tables S2,4*) were previously linked to *Pcc* virulence (*34*). These included 53 upregulated and 65 downregulated genes in *Blvra*^*-/-*^ *vs. Blvra*^+/+^ mice (*Tables S2,4*). The direction of gene regulation showed a significant association with previous studies (Fisher’s exact test, *p*<0.0001, *Tables S2,4*) for 92% of the upregulated genes (49 out of 53) and 49% of downregulated genes (32 out of 65) (*34*). Among the upregulated genes were four *Plasmodium* Interspersed Repeat (*pir*) genes (*PCHAS_030190, PCHAS_041960, PCHAS_104230*, and *PCHAS_110030*), encoding variant surface CIR proteins linked to parasite “immune evasion” (*35, 36*) and genes involved in the parasite metabolic adaptation (*34*). Other virulence genes, included 17 *Pc-fam* (or rodent malaria parasites; RMP-fam) genes (*37*); and eight genes encoding exported proteins (*34*), including the *cir* gene *PCHAS_110030*, a *P. falciparum* RIF orthologue linked to *Pcc* virulence (*36, 38*). These observations suggest that *Pcc* parasite virulence (*i*.*e*., ability to cause disease or damage to the host) increases in *Blvra*^*-/-*^ *vs. Blvra*^+/+^ mice. We then asked whether the genetic fingerprint of higher virulence in parasites from *Blvra*^*-/-*^ *vs*.

*Blvra*^+/+^ mice contributes to increased malaria mortality. In strong support of this hypothesis, the incidence of mortality (*Fig. 2G*), parasitemia and parasite burden (*Fig. 2H*) were increased in *Blvra*^*-/-*^ mice infected with parasites isolated from *Blvra*^*-/-*^ *vs. Blvra*^+/+^ mice (*Fig. 2G,H*). This suggests that the protective effect of bilirubin against malaria is mediated via a resistance mechanism that reduces *Plasmodium* burden and virulence.

### Unconjugated bilirubin accumulates specifically in *P. falciparum*-infected RBC

To explore the mechanism via which bilirubin reduces *Plasmodium* virulence, we asked whether bilirubin targets *P. falciparum* directly inside RBC *in vitro*. We used scanning electron microscopy (SEM) to confirm that unconjugated bilirubin precipitates in culture medium (*39*) (*Fig. S10A*), accounting for a 67% reduction of its expected concentration (*Fig. S10B-D*), as determined using the UnaG-based assay (*Fig. S1B,C*) (*21, 22*). There was a marked accumulation of bilirubin in *P. falciparum* 3D7 iRBC exposed *in vitro* to unconjugated bilirubin, at a concentration in the range of *P. falciparum* malaria patients (*Fig. 3A, Table S5*). In sharp contrast, there was no detectable accumulation of bilirubin in non-infected RBC, similar to vehicle-treated controls (*Fig. 3A, Table S5*). This suggests that unconjugated bilirubin accumulates specifically in *Plasmodium*-iRBC when present at a concentration in the range of that seen in *P. falciparum* malaria patients.

**Figure 3.**
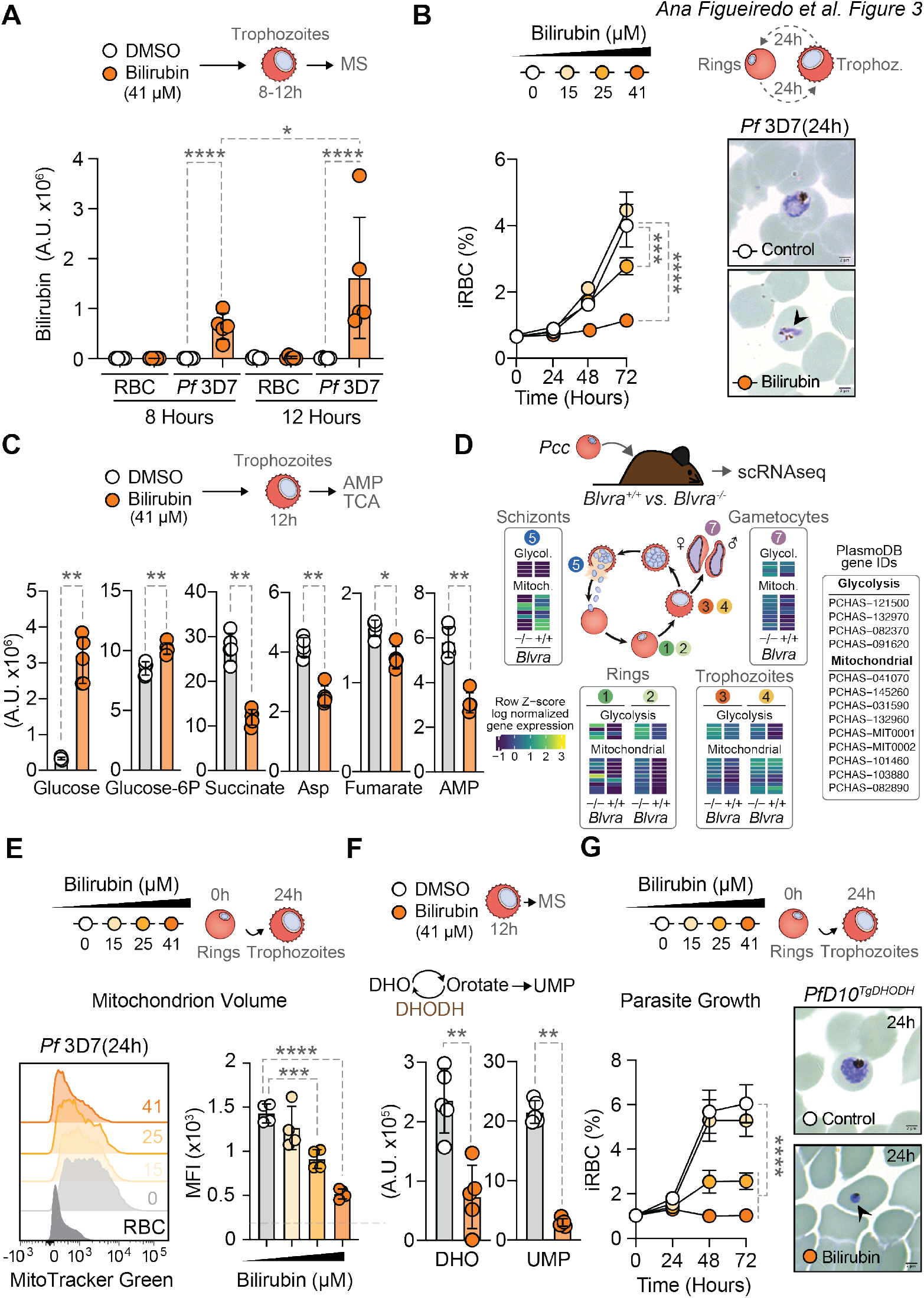
Bilirubin inhibits *P. falciparum* proliferation and disrupts its mitochondrion integrity and function. (**A**) Bilirubin accumulation in *P. falciparum* 3D7 (*Pf*3D7) iRBC (trophozoites), 8 and 12h after exposure to unconjugated bilirubin (41 µM) or vehicle. Data represented as mean ± SD, from one experiment, with 5 technical replicates. (**B**) Percentage (*left panel*) of *Pf*3D7 iRBC, 24-72h after exposure of ring stages to increasing concentrations of unconjugated bilirubin. Data represented as mean ± SEM, pooled from six independent experiments with similar trend, with four technical replicates *per* experiment. *Right panel* is a representative Giemsa-stained thin smear, 24h after exposure to unconjugated bilirubin (41 µM) or vehicle. Black arrowhead highlights parasite nuclear DNA fragmentation. Scale bar: 2 µm. (**C**) Relative levels of Glucose, Glucose 6-phosphate, Succinate, Aspartate (Asp), Fumarate and Adenosine Monophosphate (AMP) in *Pf*3D7 iRBC (trophozoites), detected by Mass Spectrometry 12h after exposure to unconjugated bilirubin (41 µM) or vehicle. Data represented as mean ± SD, from one experiment, with 5 technical replicates (*See Table S5*). (**D**) Schematic representation of differentially expressed genes involved in metabolic processes in *P. chabaudi chabaudi* iRBC isolated from *Blvra*^*-/-*^ vs. *Blvra*^*+/+*^ mice, as determined by scRNA sequencing, in the different Populations identified in (Fig. 2C-F). (**E**) Representative histograms (*left panel*) and quantification of median fluorescence intensity (MFI; *right panel*) of mitochondrion volume (MitoTracker Green), 24h after exposure of *Pf*3D7 iRBC (rings) to increasing concentrations of unconjugated bilirubin. Grey histogram represents background staining in non-infected RBC and dotted grey line (*right panel*) represents the average background signal from all replicates in non-infected RBC. Data represented as mean ± SD from four replicates in one out of two independent experiments with similar trend. (**F**) Relative level of dihydroorotate (DHO, *left panel*) and uridine monophosphate (UMP, *right panel*), in *Pf*3D7-iRBC (trophozoites), detected by Mass Spectrometry 12h after exposure to unconjugated bilirubin (41 µM) or vehicle. (*See Table S5*). (**G**) Percentage (*left panel*) of *PfD10*^*TgDHODH*^ iRBC (*P. falciparum* D10 transgenic strain expressing a cytoplasmic *Saccharomyces cerevisiae* DHODH) (rings), after exposure to increasing concentrations of unconjugated bilirubin. Data represented as mean ± SEM, pooled from 3 independent experiments with similar trend, with four biological replicates *per* experiment. *Right panel* is representative Giemsa-stained thin smears, 24h after exposure to unconjugated bilirubin (41 µM) or vehicle. Black arrowhead highlights pyknotic parasites. Scale bar: 2µm. Dots in (A,E,F) represent technical replicates and in (C) individual mice. *P* values determined using: (A, E) Ordinary One-Way ANOVA with Tukey’s multiple comparison test, (B, G) Two Way ANOVA with Tukey’s multiple comparison test, and (C, F) Mann Whitney U test. *p<0.05; **p<0.01; ***p<0.001; ****p<0.0001. Concentration of unconjugated bilirubin in (A,B,C,E,F,G) were calculated according to the UnaG-based assay (*21, 22*)(*see Fig. S10*).

### Unconjugated bilirubin arrests the proliferation and kills *P. falciparum*

Unconjugated bilirubin inhibited the proliferation of *P. falciparum* 3D7 (*Fig. 3B*) as well as that of the *P. falciparum* multidrug-resistant Dd2 (*Fig. S11A*) and IPC 5202 (*Fig. S11B*) strains *in vitro*. This anti-proliferative effect was dose-dependent, that is, the higher the concentration of bilirubin the lower *P. falciparum* proliferation (*Fig. 3B; Fig. S11A,B*). Importantly, the anti-proliferative effect of bilirubin occurred *in vitro*, at concentrations in the range of those of *P. falciparum* malaria patients, as assessed using the UnaG-based assay (*Fig. 1E*).

The anti-proliferative effect of bilirubin was not associated with the induction of hemolysis, as assessed by microscopy (*Fig. 3B; Fig. S11A,B*) and confirmed by lactate dehydrogenase (LDH) release from lysing RBC (*Fig. S11C*). This suggests that unconjugated bilirubin targets *Plasmodium* specifically in RBC without causing hemolysis.

A water-soluble bilirubin ditaurate derivative failed to inhibit the proliferation of *P. falciparum* 3D7 (*Fig. S11D*), instead promoting ring stage proliferation at concentrations in the 40-80 µM range (*Fig. S11D*). This suggests that the anti-proliferative effect of bilirubin is restricted to its unconjugated form, which diffuses across cellular membranes (*28, 29*) and accumulates specifically in *Plasmodium*-iRBC (*Fig. 3A, Table S5*).

Biliverdin had no anti-proliferative effect on *P. falciparum* 3D7 (*Fig. S12A*), Dd2 (*Fig. S12B*) or IPC 5202 (*Fig. S12C*) strains, at concentrations up to 10 times higher than those detected in *Pcc*-infected *Blvra*^*-/-*^ mice (*Fig. S2D*). Of note, at a maximal concentration of 120µM there was a marginal effect of biliverdin on the proliferation of *P. falciparum* 3D7 (*Fig. S12A*) and IPC 5202 (*Fig. S12C*) strains, consistent with previous observations (*40*). This suggests that circulating biliverdin does not exert anti-proliferative effects on blood stages of *Plasmodium* at concentrations in the range of those detected in *Pcc*-infected *Blvra*^*-/-*^ mice (*Fig. S2E*).

### Unconjugated bilirubin impairs *Plasmodium* energy metabolism

Unconjugated bilirubin caused a striking accumulation of glucose and, to a lower extent, glucose-6-phosphate in *P. falciparum* 3D7 iRBC, as compared to control iRBC exposed to vehicle (*Fig. 3C, Table S5*). Moreover, unconjugated bilirubin reduced the relative levels of succinate, aspartate, fumarate and adenosine monophosphate (AMP) in *P. falciparum* 3D7 iRBC, compared to control vehicle-treated iRBC (*Fig. 3C, Table S5*). This suggests that bilirubin compromises *P. falciparum* central carbon metabolism (*41*).

Consistent with the notion that unconjugated bilirubin compromises *Plasmodium* spp. central carbon metabolism, *Pcc* ring stages and trophozoites from *Blvra*^*-/-*^ mice showed an increase in the expression of a number of glycolytic genes, including *PCHAS-121500* (enolase, putative; ENO); *PCHAS-100970* (glucose-6-phosphate isomerase, putative), *PCHAS-1329700* (phosphoglycerate kinase, putative), *PCHAS-082370* (phosphoglycerate kinase; PGK, putative) and *PCHAS-091620* (phosphoglycerate mutase; PGM1, putative), compared to control *Blvra*^+/+^ mice (*Fig. 3D, Tables S2,3*). This suggests that unconjugated bilirubin compromises the central carbon metabolism of *Plasmodium* via a mechanism that targets the expression of the parasite’s glycolytic genes.

Moreover, *Pcc* ring stages and early trophozoites from *Blvra*^*-/-*^ mice also presented an increase in the expression of a number of genes involved in mitochondrion ATP metabolic processes, including *PCHAS-041070* (ATP synthase F0 subunit d-like protein, putative), *PCHAS-145260* (ATP synthase subunit beta, mitochondrial, putative), *PCHAS-031590* (ATP synthase subunit alpha, mitochondrial, putative) and *PCHAS-MIT0001* (cytochrome c oxidase subunit 3; COX3) (*Fig. 3D, Tables S2,3*). This suggests that unconjugated bilirubin compromises *Plasmodium* mitochondrion function *in vivo*.

### Unconjugated bilirubin disrupts *Plasmodium* mitochondrion function

Consistent with the notion that unconjugated bilirubin compromises *Plasmodium* spp. mitochondrion, *P. falciparum* 3D7 mitochondrion volume was severely reduced upon *in vitro* exposure to unconjugated bilirubin, as assessed by flow cytometry (*Fig. 3E*) and confirmed by live confocal microscopy imaging (*Fig. S13A,B*). This effect was dose-dependent, that is, the higher the concentration of bilirubin, the lower the parasite’s mitochondrion volume (*Fig. 3E*), at concentrations in the range of *P. falciparum* malaria patients, as assessed using the UnaG-based assay (*Fig. 1E*). Equimolar concentrations of biliverdin failed to reduce *P. falciparum* mitochondrion volume *in vitro* (*Fig. S13C*).

Unconjugated bilirubin decreased *P. falciparum* 3D7 mitochondrion membrane potential (*Fig. S14A*) and superoxide accumulation (*Fig. S14B*) *in vitro*, compared to vehicle-treated controls (*Fig. S14A,B*). This effect was dose-dependent, that is, the higher the concentration of bilirubin, the lower the parasite’s membrane potential (*Fig. S14A*) and superoxide accumulation (*Fig. S14B*). Equimolar amounts of biliverdin failed to reduce *P. falciparum* 3D7 mitochondrion membrane potential (*Fig. S14C*) or superoxide accumulation (*Fig. S14D*).

### Bilirubin represses *Plasmodium’s* mitochondrion pyrimidine synthesis

*Plasmodium spp*. replication relies on *de novo* pyrimidine synthesis via the conversion of dihydroorotate (DHO) into orotate, catalyzed by the mitochondrial inner membrane dihydroorotate dehydrogenase (DHODH) (*42*). Exposure of *P. falciparum* 3D7 iRBC to unconjugated bilirubin caused a marked reduction in DHO and uridine monophosphate (UMP), a downstream product of DHODH in the parasite’s *de novo* pyrimidine synthesis (*Fig. 3F, Table S5*). This suggests that the anti-proliferative effect of unconjugated bilirubin is exerted via a mechanism that impairs mitochondrion-dependent pyrimidine synthesis.

### The anti-malarial effect of bilirubin acts beyond *Plasmodium’s* mitochondrion

To determine whether the anti-proliferative effect of bilirubin relies exclusively on the inhibition of mitochondrion pyrimidine synthesis, we used a transgenic *P. falciparum* D10 strain expressing a cytoplasmic *Saccharomyces cerevisiae* DHODH (*PfD10*^*TgDHODH*^) (*42*). In contrast to the parental *PfD10* strain, *PfD10*^*TgDHODH*^ parasites can support pyrimidine synthesis irrespectively of the mitochondrion (*42*). As expected (*43*), inhibition of the mitochondrion electron transport chain cytochrome bc_1_ complex by Atovaquone (ATQ) arrested the proliferation and killed the parental *PfD10* but not the *PfD10*^*TgDHODH*^ strain (*Fig. S15A*). In contrast however, unconjugated bilirubin arrested the proliferation and killed both the parental *PfD10* and the *PfD10*^*TgDHODH*^ strains (*Fig. 3G; Fig. S15B*). This suggests that the anti-proliferative effect of bilirubin acts beyond the inhibition of *P. falciparum* pyrimidine synthesis.

### Bilirubin inhibits Hz crystallization

Exposure of *P. falciparum* 3D7 to unconjugated bilirubin *in vitro* was associated with dispersion of Hemozoin (Hz) crystals in the iRBC, as visualized and quantified by live confocal microscopy imaging (*Fig. 4A; Fig. S16A,B*). Equimolar amounts of biliverdin had no effect on Hz, similar to vehicle controls (*Fig. S16A-C*). This suggests that bilirubin interferes directly or indirectly with the detoxification of the heme extracted from hemoglobin into Hz crystals.

**Figure 4.**
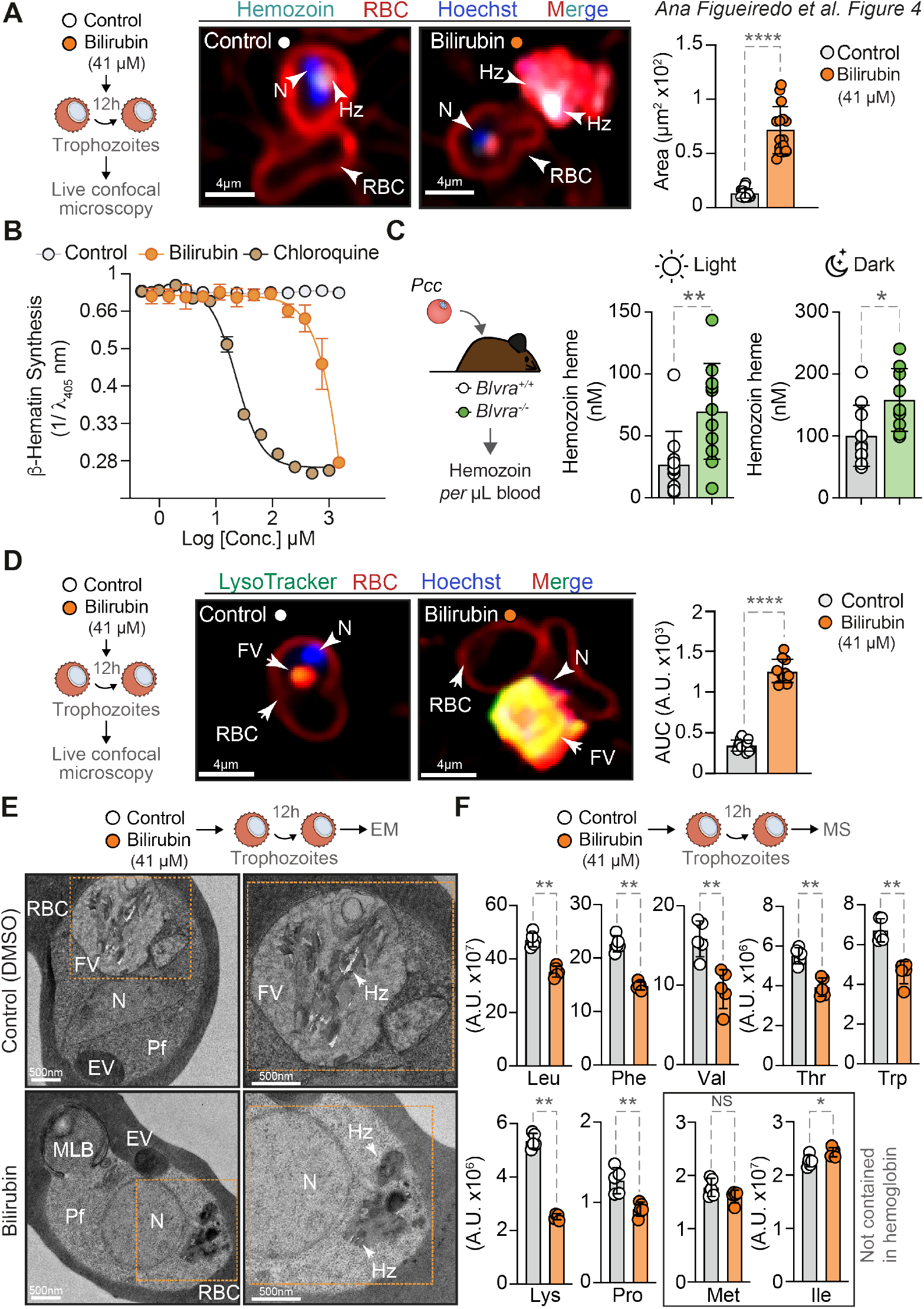
Bilirubin inhibits Hz formation and disrupts *P. falciparum* food vacuole. (**A**) Live confocal microscopy of *Pf*3D7 iRBC (trophozoites), 12h after exposure to unconjugated bilirubin (41 µM) or vehicle. Representative images (*left panels*) and quantification of Hz cellular intensity distribution (laser reflection mode, cyan; area, *right panel*). RBC membranes stained with wheat germ agglutinin (RBC; red), and nuclei (N) with Hoechst (DNA: blue). Data in *right panel* represented as mean ± SD from two independent experiments (n=10-15 parasites). (**B**) Relative inhibition of β-hematin crystallization by bilirubin *vs*. chloroquine. Equimolar amounts of Brivanib alaninate (a tyrosine kinase inhibitor) were used as control. The relative inhibition of β-hematin crystallization was inferred from heme accumulation measured at λ_405_ nm (*71*) and is represented as the mean of λ/λ _405_ nm ± SD from four replicates in one out of three independent experiments with similar trend. (**C**) Hz quantification in blood from *Blvra*^*+/+*^ and *Blvra*^*-/-*^ 7 days after *Pcc* infection, at daily light (*left panel*) and dark (*right panel*) cycle. Hz (*i*.*e*., nM heme), represented as mean ± SD, pooled from two independent experiments with similar trend (n=11; light cycle and n=9-10; dark cycle; *per* genotype). (**D**) Representative images (*left panels*) and quantification (*right panel*) of food vacuole (Area under the curve; AUC). RBC membranes stained with wheat germ agglutinin (RBC; red), parasite’s food vacuole (FV) with LysoTracker Green (green) and nuclei (N) with Hoechst (DNA: blue). Data in *right panel* represented as mean ± SD from two independent experiment (n=10-15 parasites), as in (A). (**E**) Representative transmission electron microscopy images of *Pf*3D7 iRBC (trophozoites), 12h after exposure to unconjugated bilirubin (41 µM) or vehicle. *Right panel* correspond to amplifications of the orange dotted area highlighted in *left panel*. EV: Endocytic vesicle, FV: Food vacuole, Hz: Hemozoin, MLB: Multilamellar bodies, N: Nucleus, RBC: Red blood cell. Images are representative of three independent experiments with similar results. Scale bars: 500 nm. (**F**) Relative levels of amino acids in *Pf*3D7-iRBC (trophozoites) (Leucine (Leu), Phenylalanine (Phe), Valine (Val), Threonine (Thr), Tryptophan (Trp), Lysine (Lys), Proline (Pro), Methionine (Met) and Isoleucine (Ile)) as detected by Mass Spectrometry, 12h after exposure to unconjugated bilirubin (41 µM) or vehicle (See Table S5). Circles in (A,D) represent individual parasite, in (C) individual mice and (F) technical replicates. *P* values determined using: (A, C, D, F) Mann-Whitney U test. NS: not significant; *p<0.05, **p<0.01; ****p<0.0001. Concentration of unconjugated bilirubin in (A,B,C,E,F,G) were calculated according to the UnaG-based assay (*21, 22*) (*see Fig. S10*).

We then asked whether bilirubin interferes directly with Hz crystallization. Consistent with this hypothesis, computational simulation of a putative bilirubin docking in the deep groove on the fastest growing (*i*.*e*., 001) face of the crystal suggested that bilirubin interferes directly with Hz crystallization (*Fig. S17*). The two top scoring docking poses suggest that the adsorption of bilirubin to the crystal surface is facilitated by hydrogen bonding between its carboxylic acid moiety and the pyrrole groups of heme in the nascent Hz crystals (*Fig. S17*). This suggested that bilirubin hinders heme incorporation in nascent Hz crystals, impairing the parasite’s capacity to neutralize cytotoxic labile heme (*44, 45*). This hypothesis was confirmed using a standard *in vitro* assay to quantify the crystallization of β-hematin, a synthetic heme adduct chemically and spectroscopically identical to Hz (*46*). Unconjugated bilirubin inhibited β-hematin formation (*Fig. 4B*), mediated by the lipid mimic Nonidet P-40 detergent, as monitored by the accumulation of heme as a *bis*-pyridyl complex (*47*). The IC_50_ of bilirubin was 1-1.2 mM, that is, 50-60 times less potent than that of chloroquine (IC_50_=20μM) in the same assay.

Consistent with the notion that bilirubin inhibits Hz crystallization *in vitro, Pcc* infection in *Blvra*^*-/-*^ mice was associated with higher levels of Hz accumulation in circulating iRBC, as compared to iRBC from control *Blvra*^+/+^ mice, at the light and dark cycle of *Pcc* infection (*Fig. 4C*). This suggests that bilirubin acts as a “physiologic” anti-malarial metabolite that interferes with Hz crystallization, akin to quinoline-based antimalarial drugs (*45, 48*).

### Bilirubin disrupts *P. falciparum* food vacuole

Inhibition of Hz crystallization by quinoline-based antimalarial drugs compromises the parasite’s digestive food vacuole (*45, 48*), suggesting that bilirubin also disrupts the parasite’s food vacuole. In support of this hypothesis, unconjugated bilirubin disrupted the food vacuole of *P. falciparum* 3D7 trophozoites, as revealed by leakage of its acidic content, quantified by confocal microscopy live imaging (*Fig. 4D; Fig. S18A,B*). Equimolar amounts of biliverdin failed to disrupt *P. falciparum* food vacuole, similar to vehicle-treated controls (*Fig. S18A-C*).

Disruption of *P. falciparum* food vacuole by bilirubin was further confirmed by transmission electron microscopy (TEM) (*Fig. 4E; Fig. S19*), revealing the formation of multilamellar bodies (*Fig. 4E; Fig. S19*), reminiscent to those observed when targeting endosomal vesicle delivery to the food vacuole (*49*). This was associated with the accumulation of Hz crystals in the parasite’s cytoplasm (*Fig. 4E; Fig. S19*), with a structural appearance consistent with the loss of its characteristic sharp rectangular morphology (*Fig. 4E; Fig. S19*). These “rounded-edged” cytoplasmic Hz crystals were not observed in parasites exposed to equimolar amounts of biliverdin or to vehicle (*Fig. 4E; Fig. S19*).

*Plasmodium* food vacuole is vital for the acquisition of essential amino acids (AA) contained in hemoglobin (*50*). That bilirubin impairs this vital process is supported by the reduction in the relative amount of these essential AA in *P. falciparum* 3D7 trophozoites exposed to unconjugated bilirubin, as illustrated for: Leucine (Leu), Phenylalanine (Phe), Valine (Val), Threonine (Thr), Tryptophan (Trp), Lysine (Lys) and Proline (Pro) (*Fig. 4F, Table S5*). This was also observed for non-essential AA contained in hemoglobin, including: Aspartate (Asp) (*Fig. 3C, Table S5*), Arginine (Arg), Tyrosine (Tyr) and Serine (Ser) (*Fig. S20A, Table S5*). In sharp contrast, AA not contained in hemoglobin, which are not acquired via the food vacuole (*50*), were not affected by unconjugated bilirubin, as illustrated for methionine (Met) and isoleucine (Ile) (*Fig. 4F*). Moreover, bilirubin had no effect on the AA content of non-infected RBC, compared to vehicle-treated control*s* (*Fig. S20B, Table S5*). This suggests that bilirubin compromises specifically the capacity of *P. falciparum* food vacuole to extract essential AA from hemoglobin.

A number of genes involved in hemoglobin digestion by the food vacuole were also increased in *Pcc* ring stages and trophozoites from *Blvra*^*-/-*^ *vs. Blvra*^+/+^ mice (*Fig. S20C, Tables S2,3*). These included *PCHAS-083340* (M18 aspartyl aminopeptidase, putative), *PCHAS-113650* (falcilysin; FLN, putative; highly active at acidic pH consistent with its critical role in hemoglobin degradation), *PCHAS-131350* (M17 leucyl aminopeptidase, putative) and *PCHAS-131310* (M17 leucyl aminopeptidase, putative) (*Fig. S20C, Tables S2,3*), suggesting that bilirubin also acts *in vivo* to compromise the capacity of *Plasmodium* trophozoites to extract essential AA from hemoglobin, likely contributing to the anti-malarial effects of unconjugated bilirubin.

## Discussion

Our findings support the notion that the accumulation of unconjugated bilirubin in plasma during *Plasmodium* spp. infection (*Fig. 1B-E,G*) represents a protective metabolic response to malaria (*Fig. 1H,I,L*). This is in keeping with the growing evidence suggesting that bilirubin exerts major physiological functions (*22, 51, 52*), challenging the strongly held notion that unconjugated bilirubin is a final “waste product” that accumulates in plasma due to liver dysfunction (*53*).

The protective effect of unconjugated bilirubin against *Plasmodium* infection is propelled by BVRA (*Fig. 1G,H*) and sustained by the inhibition of bilirubin conjugation via the repression of hepatic UGT1A1 (*Fig. 1J-L*). Consistent with our findings, a number of *UGT1A1* genetic hypomorphic variants are associated with mild unconjugated nonhemolytic hyperbilirubinemia in individuals of African ancestry, with the longest co-evolution with *Plasmodium* (*54, 55*). These include the hypomorphic variant *UGT1A1*\*28 (rs3064744) responsible for Gilbert’s syndrome (*19, 56*), which reduces *UGT1A1* transcription by 70% in an estimated prevalence of 15-25% of individuals of African ancestry, compared to 0-5% and 5-10% in Asian and Caucasian ancestries (*54*). However, an association between polymorphisms in or near the *UGT1A1* locus (chr2:233,760,270-233,773,300) and *P. falciparum* malaria severity was not reported in previous GWAS studies (*57-61*). One possible explanation for this is that *UGT1A1* genetic variants can increase the incidence and severity of neonatal jaundice (*54, 55*). Moreover, *UGT1A1* genetic variants might co-segregate with balanced polymorphisms conferring protection against malaria via the induction of heme catabolism, such as sickle hemoglobin (*10*).

Targeting of *Plasmodium* inside RBC by unconjugated bilirubin (*Fig. 3A*) is consistent with unconjugated bilirubin crossing cellular membranes (*28, 29*), despite its binding to albumin in plasma (*16*). Presumably this occurs via aqueous diffusion (*62*), suggesting that cellular membranes have higher affinities towards unconjugated bilirubin, compared to albumin (*62*).

The inhibition of *Plasmodium* spp. proliferation (*Fig. 2A, 3B, S5*) and virulence (*Fig. 2G, Table S2, S4*) by unconjugated bilirubin is associated with repression of the parasite’s capacity to consume glucose via glycolysis (*Fig. 3C,D*), presumably inhibiting mitochondrion tricarboxylic acid (TCA) cycle function (*Fig. 3C,D*). Moreover, unconjugated bilirubin disrupts the parasite’s mitochondrion structure (*Fig. 3E, S13A,B*) and function (*Fig. S14A*), compromising the *de novo* pyrimidine synthesis (*Fig. 3F*) and therefore inhibiting *Plasmodium spp*. proliferation (*42*). This, however, is not sufficient to fully explain the anti-malarial effects of bilirubin (*Fig. 3G*).

While *Plasmodium* spp. evolved to detoxify redox-active labile heme into redox-inert Hz crystals, the proliferation of these parasites inside RBC is associated with intravascular hemolysis and the release of labile heme into plasma (*6, 8-11*). This induces the expression of HO-1 in the infected host, which catabolizes labile heme into biliverdin (*8, 9, 14*), fueling the production of bilirubin by BVRA. As it accumulates in plasma, unconjugated bilirubin inhibits Hz crystallization (*Fig. 4A-C, S16A, B*), similar to, although far less potent than quinoline-based antimalarial drugs such as chloroquine (*Fig. 4B*) (*45, 48*). The inhibition of Hz crystallization leads to the disruption of the parasite’s food vacuole (*Fig. 4D,E, S18*), compromising the extraction of essential AA from hemoglobin (*Fig. 4F*) while presumably leading to accumulation of cytotoxic labile heme.

In conclusion, the induction of bilirubin production in response to *Plasmodium* spp. infection is a metabolite-based resistance mechanism (*63*) against malaria. We speculate that while evolutionarily conserved, this defense strategy carries as an evolutionarily trade-off (*64*), the insidious prevalence of neonatal jaundice (*65, 66*), which can lead to encephalopathy (*67*). Considering the extraordinary selective pressure exerted by malaria on human populations (*68*), it is conceivable that the anti-malarial effects of unconjugated bilirubin outcompeted the fitness costs associated with high incidence of neonatal jaundice in populations originating in endemic areas of malaria (*69, 70*).

## Acknowledgments

The authors are indebted to all members of the Inflammation group (GIMM) for insightful technical and intellectual contributions, to the staff at the flow cytometry facility, as well as to the staff at the GIMM animal facility. The authors thank DR de Waart (University of Amsterdam) for quantification of unconjugated bilirubin by HPLC (not shown), Akhil Vaidya (Drexel University, USA) for kindly providing the *PfD10* and *PfD10*^*TgDHODH*^ strains and insightful advice, the teams at Centre de Recherches Médicales de Lambaréné for support of the DEMIT study, Lara Bardtke, Paolo Kroneberg, Anna Karolina Kneller, Nathalie Schirra, Johanna Lioba Schöllgen, and Claudia Conrad for sample and data processing at Charité - Universitätsmedizin Berlin, Oleg Chertkov (Católica Biomedical Research Centre, Portugal), Protein Purification Research Facility and the Bacterial Imaging Cluster at ITQB NOVA for providing support for protein purification and quantification, and to Ana Malheiro (Advanced Electron Microscopy, Imaging and Spectroscopy – AEMIS) for the work that was carried out in part through the use of the INL User Facilities.

## Funding

This work was financed by Fundação para a Ciência e Tecnologia (2020.04797.BD and COVID/BD/153665/2024 to AF; GHTMUID/04413/2020, LA-REAL-LA/P/0117/2020 and 2022.02426.PTDC to FN; FEDER/29411/2017 to SR; 2020.04797.BD to DD; UIDB/04565/2020, LA/P/0140/2020 and 2022.03627.PTDC to SP; 2022.08590.PTDC_EXPL DOI 10.54499/2022.08590.PTDC to JK; 2021.03494.CEECIND DOI 10.54499/2021.03494.CEECIND/CP1674/CT0004 to RM; 2023.09168.CEECID to EJ; FEDER/29411/2017, PTDC/MED-FSL/4681/2020 DOI 10.54499/PTDC/MED-FSL/4681/2020, 2022.02426.PTDC DOI 10.54499/2022.02426.PTDC and Congento LISBOA-01-0145-FEDER-022170 to MPS).

European Union’s Horizon 2020 research and innovation programme under the Marie Skłodowska-Curie (grant agreement No.: 955321 to AGGS, grant agreement No.:753236 to RM).

DFG Cluster of Excellence ‘‘Balance of the Microverse’’ EXC 2051; 390713860 (EJ, MPS as associated member).

DFG IRTG 2290 “Molecular interactions in malaria” (P.T.L., F.K. and the DEMIT study).

Gulbenkian Foundation (SR, MPS and IBB 2021-51/BI-D/2021 to ST).

la Caixa Foundation HR18-00502 (EJ, JK, MPS).

Human Frontier Science Program (LT0043/2022-L to JK).

Lise Meitner Excellence Programme of the Max Planck Society (SP).

European Molecular Biology Organization (EMBO Long-term Fellowship ALTF290-2017 to RM).

European Union’s Horizon 2020 research and innovation programme (Grant 955321).

Academy of Finland (Grant 329278 to LLE).

Sigrid Juselius Foundation (LLE).

Biocenter Finland (LLE).

ELIXIR Finland (LLE).

American Heart Association/Paul Allen Frontiers Group (Project 19PABH134580006 to BDP).

NIH/NIA (BDP by 1R21AG073684-01, R01AG071512).

The Johns Hopkins Catalyst Award (BDP).

Solve ME/CFS Initiative (Grant 90089823 to BDP).

US Public Health Service (Grant DA044123 to BDP).

European Research Council (Grant 101001521 to MJA).

Oeiras-ERC Frontier Research Incentive Awards (MPS).

H2020-WIDESPREAD-2020-5-952537 SymbNET Research Grants (MPS)

## Author Contributions

Conceptualization: MPS, AF, RM, KV, SP

Data curation: AGGS, SJ, LVN, CW, PTL, FK, SV, TP, BD

Formal analysis: AGGS, SJ, LVN, CW, PTL, FK, SV, TP, BD

Resources: GMN, JN, CW, PTL, FK, BP, MJA, GB, AFM

Investigation: AF, SR, EJ, JT, DD, ALS, SP, SPa, STR, MM, SC, MA, JK, GMN, JM, CW, PTL, FK, MA, CC, RM

Visualization: AF, RM, MPS

Funding acquisition: MPS, RM,

FK Project administration: MPS, RM

Supervision: MPS, RM, FN, SJ, LLE, SNP, SP, JK, PJB, FK, ET, KV

Writing – original draft: MPS

Writing – review & editing: MPS, AF, RM, STR, SR

## Data availability

Clinical patient data will be made available by Florian Kurth upon reasonable request. Single-cell RNAseq and Metabolomics data from this publication will be made public upon publication.

## Notes

### Competing Interest Statement

The authors have declared no competing interest.

