## Supplementary material for "A Metabolite-Based Resistance Mechanism Against Malaria": Methods and Supplementary Figures

### Materials and Methods

#### *Human data*

Clinical patients' data were collected from a prospective clinical cohort study, conducted between March and July 2022 at Centre de Recherches Médicales de Lambaréné (CERMEL), Gabon. The study was approved by the ethics committee of CERMEL (DEMIT-GAB: CEI 18/2021, and all patients provided written informed consent and were included in this analysis if they fulfilled the following criteria: a) age ≥18 years; b) microscopically confirmed asexual *P. falciparum* infection by positive blood smear; c) blood drawn before or ≤24h after initiation of antimalarial treatment. Patients were excluded in case of known liver (e.g., hepatitis, hepatic cancer) or hematologic disease, pregnancy or breastfeeding, mixed *Plasmodium* infection and with missing initial parasitemia data, hematology data, liver function data or bilirubin data. Patients were grouped as asymptomatic within 7 days before and 7 days after diagnosis or symptomatic malaria according to WHO criteria (*1*). Among 42 participants meeting the criteria for this analysis, the mean age was 33.1 (SD=16.97) years (asymptomatic: 36.0 (SD=19.6); symptomatic: 29.9 years (SD=13.2). Overall, 48% (20/42) were male and 52% (22/42) female (males among asymptomatic: 45% (10/22); symptomatic: 50% (10/20)). Median initial parasitemia was 576 iRBC (IQR: 131 to 4,201) *per* µL (asymptomatic: 322,5 [110 to 851]; symptomatic: 4,289 [322 to 48,218]). Unconjugated bilirubin was measured exclusively by the UnaG-based assay (*2, 3*) in 4 asymptomatic and 1 symptomatic malaria patients out of the 42 patients. Parasitological analyses, hematology and biochemistry were performed by accredited laboratories at CERMEL and Charité - Universitätsmedizin Berlin. Study data were collected and managed using REDCap (Research Electronic Data Capture) tools hosted at CERMEL and Charité - Universitätsmedizin Berlin (*4*).

#### *Mice*

Mice were bred and maintained under specific pathogen-free (SPF) conditions at the Gulbenkian Institute of Molecular Medicine (GIMM), housed at standard *vivarium* temperature (22°C) on a regular light cycle, lights on from 8:00 am (ZT 0) to 8:00 pm (ZT 12). When indicated, mice were kept on inverted light cycle, lights on from 8:00 pm (ZT 0) to 8:00 am (ZT 12). Mice were maintained with free access to water and standard chow pellets (*ad libitum*). All experimental protocols were approved in a two-step procedure, by the Animal Welfare Body of the GIMM and by the Portuguese National Entity that regulates the use of laboratory animals in research (Direção Geral de Alimentação e Veterinária; DGAV). Experimental procedures followed the Portuguese (Decreto-Lei nº 113/2013) and European (Directive 2010/63/EU) legislation. C57BL/6J and DBA/2 mice were obtained from the GIMM animal facility. C57BL/6J *Blvra*$^{-/-}$ mice were generated at Ozgene (Australia), as described (*3*). Age-matched wild-type C57BL/6J mice were used as controls and co-housed at least two weeks with *Blvra*$^{-/-}$ *mice* before infection.

#### *Plasmodium chabaudi chabaudi AS infection and disease assessment*

Mice (females and males, 8-14 weeks old) were infected with *Plasmodium chabaudi chabaudi* AS (*Pcc* AS) or transgenic GFP-expressing *Pcc* AS (*Pcc* AS-$GFP_{ML}$) (*5*). Infections were performed by intraperitoneal (*i.p.*) administration of freshly isolated blood (Passage 34-39; $2x10^6$ or $2x10^5$ infected RBC diluted in 200 µL PBS) collected from a previously infected C57BL/6J mouse. Mice were monitored daily from day 0 (day of infection) onwards for parasitemia (% infected RBC; iRBC), parasite burden (number of iRBC *per* µL of blood) and survival, essentially as described (*6, 7*). Briefly, the number of RBC *per* µL of blood was quantified by flow cytometry (LSR

Fortessa X20 analyzer; BD Bioscience) using a standard concentration of reference latex beads (10 µm; Coulter® CC Size Standard L10, Beckman Coulter, # 6602796), gating on RBC, based on size and granularity and on bead population. For *Pcc* AS, parasitemia was determined manually by optical microscopy, counting the number of iRBC in at least 4 fields of Giemsa-stained blood smears (1000x magnification). For *Pcc* AS-$GFP_{ML}$, parasitemia was determined by flow cytometry, according to the percentage of RBC expressing GFP ($GFP^+$ RBC). Parasite burden, expressed as iRBC/µL, was determined by multiplying the parasitemia by the number of RBC. Representative Giemsa-stained thin blood smear images were acquired suing a Zeiss Imager Z2/ApoTome.2, equipped with an Axiocam 105 color camera, using the 100x 1. 4NA Oil immersion objective, in a 5×5 tile stitched with Zeiss's ZEN v3.1. The images were analyzed using the Fiji Software (ImageJ) and the background in parasite RGB images was corrected using the Color Correct plugin for Fiji, contributed by Gabriel Landini (https://github.com/landinig/IJ-Colour_Correct/blob/main/colour_correct.zip).

*Bilirubin supplementation during Plasmodium infection*

*$Blvra^{-/-}$* mice were infected with *Pcc* AS-GFPML ($2x10^5$ iRBC, 200µL, *i.p.*) and monitored as described above. Bilirubin (Frontier Scientific) stock solutions were prepared as described (*8*). Briefly, bilirubin was dissolved in 0.2N NaOH, buffered to pH 7.4 using 0.2N HCl, filtered (70µm cell strainer) to remove precipitates and stored (-80ºC). Absorbance was determined using a spectrophotometer (SmartSpec 3000) at 460nm. The concentration was calculated considering 53,846 $cm^{-1}$ $M\text{-}1^{-1}$ as molar extinction coefficient and following the Lambert-Beer law (A460nm=ε.C. l). Bilirubin was administered *i.p.* at 30mg/Kg or 3mg/Kg, , once daily, from day 4 to day 15 after infection.

*Repression of hepatic Ugt1a1 in vivo*

A recombinant adeno-associated virus serotype 8 (AAV8) encoding the *Staphylococcus aureus* (Sa) CRISPR associated protein 9 (Cas9) and a single guide RNA (gRNA) targeting *Ugt1a1* (AAV8-gRNA- *Ugt1a1*) was administered to 2-4 days old DBA/2 mice ($3x10^{14}$ viral particles/Kg body weight, in PBS; *i.v.* retro-orbitally), as described (*9*). Controls were transduced with a AAV8 encoding SaCas9 without the targeting gRNA (AAV8-Cas9). Female transduced mice were infected with *Pcc* ($2x10^6$ iRBC, 200µL, *i.p.*) at ~10 weeks after birth and monitored for survival, parasite burden and disease severity, as described above. Adult male transduced mice were sacrificed for organ collection to assess *Ugt1a1* deletion efficiency by Western blot.

*Plasmodium* virulence assay

*Pcc* parasites were adoptively transferred from *$Blvra^{+/+}$ vs. $Blvra^{-/-}$* mice into female *$Blvra^{-/-}$* mice, essentially as described (*7*). Briefly, blood was collected from both *$Blvra^{+/+}$* and *$Blvra^{-/-}$* mice 7 days after *Pcc* infection and the same number of iRBC ($2x10^6$; 200 µL PBS) were passively transferred (*i.p.*) into recipient *$Blvra^{-/-}$* mice, monitored daily for disease assessment and survival, as described above.

*Pcc-infected RBC sequestration*

*$Blvra^{+/+}$* and *$Blvra^{-/-}$* mice were infected with *Pcc* and kept on inverted light cycle, lights on from 8:00 pm (ZT 0) to 8:00 am (ZT 12). Mice were sacrificed ($CO_2$ asphyxiation) 7 days after infection, perfused *in toto* via transcardiac infusion of ice-cold PBS (1X, 15 mL), organs were harvested, snap frozen in liquid nitrogen and stored at -80ºC. The accumulation of *Pcc* AS 18S rRNA in

different organs was quantified by qRT-PCR. RNA extraction and qRT-PCR were performed as described in the section RNA extraction and qRT-PCR.

*Cecal Ligation and Puncture (Sepsis)*
Cecal Ligation and Puncture (CLP) was performed in *Blvra*$^{+/+}$ and *Blvra*$^{-/-}$ mice (females and males, 8-14 weeks old), essentially as described (*10*). Briefly, mice were anesthetized (Ketamine: 12 mg/mL; and Xylazine: 1.6 mg/mL in 1xPBS; 180-200 μL *i.p.*) and subjected to a 15-20% cecum ligation and double puncture with a 23 Gauge (G) needle. A small amount of feces was extruded, and the cecum was carefully placed back into the abdominal cavity. All animals received 0.9% saline (40 mL/Kg, *i.p.*) and Imipenem/Cilastatin (25 mg/Kg, *i.p.*), starting 2h after CLP and every 12h for 3 days. Mice were monitored daily from day 0 (day of infection) onward for body weight (Ohaus® CS200 scaler, Sigma Aldrich), rectal temperature (Rodent thermometer; BIO-TK8851, Bioset), blood glucose concentration (Accu-CHECK Performa glucometer, Roche), and survival.

*Bacterial load*
Mice were sacrificed 24 hours after CLP and peritoneal fluid was obtained by peritoneal lavage (7 mL sterile PBS). Mice were perfused with sterile ice cold 1x PBS. Whole organs were harvested and homogenized under sterile conditions in 1 mL sterile ice cold 1x PBS using a dounce tissue grinder (Sigma Ref: D8939-1SET). Serial dilutions were plated onto TrypticaseSoy Agar II with 5% Sheep Blood plates (Becton Dickinson Ref:254053) and incubated (24 h at 37 °C) in air 5% $CO_2$ (aerobes) or in an airtight container equipped with the GasPak anaerobe container system (Becton Dickinson Ref 260678). Anaerobic conditions were confirmed in all experiments using BBL Dry Anaerobic Indicator Strips (Becton Dickinson Ref: 271051).

*Influenza A virus infection*
*Blvra*$^{+/+}$ and *Blvra*$^{-/-}$ mice were maintained at the GIMM BSL-2 facility, and infected with influenza A/X-31, essentially as described (*11, 12*). Briefly, mice were anesthetized with isoflurane and 30μL of inoculum ($10^3$ PFU in sterile PBS), administered intranasally. Mice were monitored daily, as described above. Viral loads were determined at day 3 post-infection from the right lower lobes of influenza A virus-infected mice. Samples were homogenized in serum-free DMEM (Gibco) using tungsten carbide beads (Qiagen) in a TissueLyser II (Qiagen) at 20 $s^{-1}$ for 3 min and supernatants were collected after centrifugation. Titration by plaque assay was performed as previously described using Madin-Darby Canine Kidney cells (*11*).

*UnaG protein synthesis and purification*
UnaG was expressed in pMAL-6P2-6xHIS in BL21(DE3) cells (*3*). Starter cultures were grown to saturation in Luria Broth (LB) (overnight at 37°C), diluted 10-fold in LB and grown to an OD600 of 0.3 at 37°C, after which cultures were moved to 18°C. UnaG expression was induced at OD600 of 0.6, by the addition of 400 μM isopropyl β-D-1-thiogalactopyranoside (16h, 18°C). Cells were harvested by centrifugation (3,800 *g*, 4°C, 25 min), resuspended in 15 mL of resuspension buffer (50 mM HEPES, 300 mM NaCl, 0.5 mM TCEP, 10% glycerol, 1 mM PMSF, 2.34 μM leupeptin, 1.45 μM pepstatin at pH 7.4) *per* liter of culture. The sample was further lysed by sonication (QSonica Sonicator) with an amplitude of 5% in a pulse mode of 0.8sec ON and 0.5sec OFF for total 30 sec. Lysate was clarified by centrifugation (26,000 *g*, 4°C, 30 min) and loaded onto an amylose column (MBPTrap™ HP Column, GE Healthcare). Protein was eluted with 20 mM maltose in protease buffer (50 mM Tris, 150 mM NaCl, 0.5 mM TCEP, 10% glycerol,

0.01% TritonX-100 at pH 7.4). MBP was removed by the addition (16h, 4°C) of PreScission Protease (GE Healthcare). UnaG was purified via a nickel column (HisTrap™ HP Column, GE Healthcare) to remove the cleaved tag and eluted with (30 mM Tris, 1M NaCl, 0.5 mM TCEP, 10% glycerol, 500mM imidazole, pH 7.4). UnaG was further purified on a gel-filtration column (S-200, GE Healthcare) in gel filtration buffer (50 mM HEPES, 300 mM NaCl, 0.5 mM TCEP, 10% glycerol at pH 7.4), concentrated using 10kDa amicon ultracentrifugal filter (Merck), and flash frozen in gel filtration buffer at 30% glycerol for storage at −80°C.

*UnaG-based assay for quantification of unconjugated bilirubin*

Blood from *Blvra*$^{+/+}$ and *Blvra*$^{-/-}$ mice was obtained from the submandibular (facial) vein, before (Day 0) and after (Days 4, 7, 15 and 25) *Pcc* infection. Samples were collected in the dark, under a red light, immediately centrifuged (1,000 *g*, 15 min.) and plasma was collected, frozen in liquid nitrogen and stored at -80ºC until used for bilirubin quantification. Total plasma unconjugated bilirubin was determined using UnaG (*13*). Plasma sample from human patients or mice non-infected or infected with *Pcc* was diluted 1:100 in PBS. 100 pM of UnaG was added to the diluted plasma sample and incubated for 10 min at RT. After incubation, fluorescence intensity was measured at excitation $\lambda_{480}$ nm and emission $\lambda_{530}$ nm using microplate reader (Promega GloMax®). Unconjugated bilirubin (Frontier Chemicals) was used as standard.

*Clinical quantification of unconjugated bilirubin*

Clinical routine bilirubin measurements were performed at accredited laboratories at Charité – Universitätsmedizin Berlin and Centre de Recherches Médicales de Lambaréné, Gabon, using the modified colorimetric Diazo method (*14*) as implemented by Roche Diagnostics. In brief, bilirubin reacts with 3,5-dichlorophenyldiazonium salt to form azobilirubin. The red color intensity of azobilirubin is directly proportional to bilirubin concentrations. For measurement of total bilirubin, an accelerator is added to the sample to solubilize albumin-bound indirect (unconjugated) bilirubin before adding the salt, allowing for photometric measurement of total bilirubin (i.e., direct (conjugated) and indirect bilirubin). Indirect bilirubin concentrations are calculated by subtracting direct from total bilirubin.

*iRFP synthesis and purification*

The iRFP gene (Addgene plasmid #31857) was subcloned into pGEX-6P-2 (GE Healthcare Life Sciences) expression vector (*3*) and subsequently transformed into BL21(DE3) cells for protein expression. Starter cultures were grown to saturation in Luria Broth (LB) (overnight at 37°C), diluted 10-fold in LB and grown to an OD600 of 0.3 at 37°C, after which cultures were moved to 18°C. iRFP expression was induced at OD600 of 0.6, by the addition of 400μM isopropyl β-D-1-thiogalactopyranoside (16h, 18°C). Cells were harvested by centrifugation (3,800 *g*, 4°C, 25 min), resuspended (15 mL; 50 mM HEPES, 300 mM NaCl, 0.5 mM TCEP, 10% glycerol, 1 mM PMSF, 2.34 μM leupeptin, 1.45 μM pepstatin; pH 7.4) *per* liter of culture. The sample was further lysed by sonication (QSonica Sonicator) with an amplitude of 5% in a pulse mode of 0.8sec ON and 0.5sec OFF for total 30 sec. Lysate was clarified by centrifugation (26,000 *g*, 4°C, 30 min.) and loaded onto GST column (GSTrap™ Column, GE Healthcare). Protein was eluted with 10mM glutathione in protease buffer (50 mM Tris, 150 mM NaCl, 0.5 mM TCEP, 10% glycerol, 0.01% TritonX-100 at pH 7.4). GST was removed by the addition of PreScission Protease (GE Healthcare) for 16 h at 4°C. iRFP was further purified on a gel-filtration column (S-200, GE Healthcare) in gel filtration buffer (50 mM HEPES, 300 mM NaCl, 0.5 mM TCEP, 10% glycerol at pH 7.4), concentrated using 10kDa amicon ultracentrifugal filter (Merck), and flash frozen in gel filtration buffer at 30% glycerol for storage at −80°C.

*iRFP-based assay for quantification of biliverdin*

Blood was harvested from the submandibular (facial) vein of *Blvra*$^{+/+}$ and *Blvra*$^{-/-}$ mice, before (Day 0) and after (Days 4, 7 and 15) *Pcc* infection. Plasma was collected in the dark, under a red light, immediately after centrifugation (1,000 *g*, 15 min.), frozen in liquid nitrogen and stored at -80ºC until used. Biliverdin concentration in plasma was quantified using biliverdin-inducible infrared fluorescent protein (iRFP) (*15*). Briefly, plasma samples were diluted (1:100 in PBS) and incubated with iRFP (87 nM; 15min at RT). Fluorescence intensity was measured at excitation $\lambda_{690}$ nm and emission $\lambda_{713}$ nm using a microplate reader (BioTek Synergy H1). Biliverdin hydrochloride (Frontier Chemicals) was used as standard.

*Bilirubin and biliverdin*

For the *in vitro* studies, stock solution (34 mM) of high purity bilirubin and biliverdin (Frontier Scientific, Inc. Logan, Ut, USA) were dissolved in DMSO (Sigma-Aldrich) and protected from light. Stock solutions were further diluted (1mM) in RPMI 1640 supplemented with lipid-rich bovine serum albumin (0.5% AlbuMAXII; Invitrogen™, Thermo Fisher Scientific). Albumin concentration in the culture medium was 75 µM. Unconjugated bilirubin was added to RBC or *P. falciparum* iRBC in culture medium (40, 80 and 120 µM) corresponding to 15, 25 and 41 µM of unconjugated bilirubin, as quantified within 2 hours using a UnaG-based assay (see above) (*2, 3*).

*Plasmodium falciparum in vitro culture*

*Plasmodium falciparum* 3D7-GFP (*Pf*3D7-GFP; MRA-1029, MR4, ATCC® Manassas Virginia; chloroquine and artemisinin-sensitive), Dd2 (*Pf*-Dd2; MRA-150, MR4, ATCC® Manassas Virginia; chloroquine-resistant), IPC 5202 (Pf5202; MRA-1240, MR4, ATCC® Manassas Virginia; artemisinin-resistant) and *Plasmodium falciparum* D10 (*Pf*D10; atovaquone-sensitive) and its transgenic derivative (*PfD10*$^{TgDHODH}$) (*16*) were co-cultured (37°C; 95% humidity, 5% of $CO_2$) with human RBC from healthy donors (5% hematocrit), replacing human serum by 0.5% AlbuMAXII (Invitrogen™, Thermo Fisher Scientific), as described (*7, 17*). Cultures were synchronized using 5% D-Sorbitol (Sigma-Aldrich). After RBC reinvasion, the parasites at the schizont stage were suspended, layered onto 70% Percoll (Sigma-Aldrich), centrifuged (1,000g; 15 min.; no brake), collected from the upper layer of Percoll cell suspension, washed (PBS 1X) and incubated in standard culture conditions until RBC reinvasion (*18*). Ring stage (approximately 10-12h after RBC invasion) or trophozoite (approximately 18-20h after RBC invasion) stage parasites were diluted to approximately 1% parasitemia and 3% hematocrit in RPMI 1640 complete medium, (75µM bovine serum albumin), seeded on a 96-well plate and exposed to vehicle (DMSO; represented in figures as 0 µM bilirubin/biliverdin) or different concentrations of bilirubin (15-41 µM, diluted in DMSO), biliverdin or water soluble bilirubin ditaurate (40-120 µM; Frontier Specialty Chemicals Inc.) for 24, 48 and 72h. Parental *Pf*D10 and its transgenic derivative *PfD10*$^{TgDHODH}$ were also treated with atovaquone (10 nM; Sigma-Aldrich) for the same period of time. Final DMSO concentration was always kept below 0.5%. Slides were prepared at 24, 48 and 72h and stained with Giemsa solution. In parallel, parasites were suspended (PBS 1X) and analyzed by flow cytometry (Beckman Coulter CytoFLEX) to determine the percentage of iRBC. *Pf*Dd2 and *Pf*5202 parasites were stained with 0.5X SYBR green I (Invitrogen, Thermo Fisher Scientific) and *Pf*D10 and *PfD10*$^{TgDHODH}$ parasites with 0.5X SYTO Deep Red Nucleic Acid

Stain (Invitrogen, Thermo Fisher Scientific) in PBS (45 min. in the dark; 37°C) and washed in PBS prior to acquisition. *Pf*3D7-GFP parasites were acquired directly ($GFP^+$ signal). Representative Giemsa-stained thin blood smear images were acquired as described for *Pcc.*

*Hemolysis assay*

*Pf*3D7 parasites cultured in 96-well plates, were synchronized and ring stages (~1% parasitemia, 3% hematocrit) were treated with different concentrations of bilirubin, as described above. At 72h after treatment, 96-well plates were centrifuged (300 *g*, 5min.), supernatants were collected and stored at 4ºC and parasitemias were quantified by flow cytometry as described above. 2% Triton X-100 was used as a positive control to establish 100% RBC lysis. LDH release into the medium was quantified using the LDH-GloTM Cytotoxicity assay (Promega, #J2380). Briefly, culture medium was diluted in LDH Storage Buffer (200mM Tris-HCl, pH 7.3, 10% Glycerol, 1% BSA) to reach a 5X dilution. LDH detection reagent (50µL of LDH detection enzyme mix + 0,25µL Reductase Substrate) was added to each sample (1:1 ratio), incubated (60 min.; RT) and the luminescence was recorded using the GloMax microplate reader (Promega). The percentage of cytotoxicity was calculated using the following formula: (Experimental LDH release – medium background)/(Maximum LDH release control – medium background) x 100.

*RNA extraction and qRT-PCR*

Mice were sacrificed by $CO_2$ asphyxiation, transcardially perfused *in toto* with ice-cold PBS (1X, 15 mL) and organs were harvested, snap frozen in liquid nitrogen and stored at -80ºC. Total RNA was extracted using tripleXtractor reagent (GRISP), chloroform, isopropanol, and ethanol, according to manufacturer's instructions. cDNA was synthesized using the Xpert cDNA Synthesis Mastermix (GRiSP), followed by qRT-PCR using the iTaq Universal SYBR Green Supermix (Bio-Rad) on a QuantStudio™ 7 Flex Real-Time PCR System (Applied Biosystems). Transcript values were calculated from the threshold cycle (Ct) of each gene using the $2^{-\Delta\Delta CT}$ method using Acidic ribosomal phosphoprotein P0 (*Arbp0*) as the housekeeping control gene. Primers for qPCR include: *Arbp0*, Fwd: 5'-CTTTGGGCATCACCACGAA-3', Rev: 5'-GCTGGCTCCCACCTTGTCT-3'; *Blvra*-deletion confirmation, Fwd: 5′-AGCCGCTGGTAAGCTCC-3′, Rev: 5′-ACCAACCACTACCACACCAAA-3′; *Blvra*, Fwd: 5'-ATTCTGCCACCATGGAAA-3', Rev: 5'-CTCCAAGGACCCAGATTTGA-3'; *Hmox1,* Fwd: 5'-TGACACCTGAGGTCAAGCAC-3', Rev: 5'-TCTCTGCAGGGGCAGTATCT-3'; *Ugt1a1,* Fwd: 5'-TCTGGCTGATGAGAAGTGACT-3'; Rev: 5'-GAAAACAACGATGCCATGCT-3'; *Pcc* AS 18S rRNA, Fwd: 5'- AAGCATTAAATAAAGCGAATACATCCTTAT-3', Rev: 5'-GGGAGTTTGGTTTTGACGTTTATGCG-3'.

*Flow cytometry (Plasmodium falciparum)*

*Pf*3D7-GFP cultures were synchronized as described before. Ring stage (approximately 10-12h after invasion) or trophozoite stage (approximately 18-20h after invasion) parasites (approximately, 3% parasitemia) were treated with vehicle (0.12-0.47% vol/vol DMSO) or different concentrations of unconjugated bilirubin (15-41 µM) or biliverdin (40-120 µM) for 24h, collected, washed with 1X PBS and stained as previously described (*7*). Cells were stained with MitoTracker™ Green FM (200 nM in 1X PBS; Invitrogen™, Thermo Fisher Scientific), MitoTracker™ Deep Red FM (200 nM in 1X PBS; Invitrogen™, Thermo Fisher Scientific), and MitoSOX™ Red Mitochondrial Superoxide Indicator (5 µM in 1X PBS; Invitrogen™, Thermo Fisher Scientific) for 20 min., 37ºC, 5% $CO_2$. Cells were washed once with 1X PBS and stained with Hoechst 33342 (10 µM in 1X PBS; Thermo Fisher Scientific) for 20 min., 37ºC, 5% $CO_2$. Cells were again washed once with 1X PBS and analyzed in a FACSAria™ Ilu Cell Sorter (BD Biosciences). FACS data was analyzed with FlowJo V10.8.1. Gating strategy illustrated in **Data S1-2**.

*Western blot*

Mice were sacrificed by $CO_2$ asphyxiation, transcardially perfused *in toto* with ice-cold PBS (1X, 15 mL) and organs were harvested and stored at -80ºC. Tissues were lysed in 2% SDS-PAGE sample buffer (100 mM Tris, pH 6.8, 20% glycerol, 4% SDS, 0.2% bromophenol blue, 100mM DTT) supplemented with 1X protease inhibitor cocktail (cOmplete™, Mini, EDTA-free Protease Inhibitor Cocktail; Roche) and homogenized in a TissueLyser II (Qiagen) with tungsten carbide beads (Qiagen). Supernatants were collected, boiled (5 min., 95ºC) and total protein was quantified at $\lambda_{280}$ nm using the DS-11 FX Spectrophotometer (DeNovix). Proteins (50 or 150µg) were resolved on a 12% SDS-PAGE and transferred to Polyvinylidene fluoride (PVDF) membranes. Membranes were blocked (1h at RT; 5% milk in 1X TBS-T), washed (1X TBS-T) and incubated (overnight at 4°C) with primary antibodies (5% BSA in 1X TBS-T): rabbit polyclonal anti-BVRA (Invitrogen, ThermoFisher Scientific, PA5-92059; 1:1000), rabbit polyclonal anti-HO-1 (Enzo Life Sciences, ADI-SPA-896F; 1:1000), rabbit monoclonal anti-GAPDH (Cell Signaling Technology, clone 14C10, #2118; 1:1000) and rabbit polyclonal anti-β-actin (Cell Signaling Technology, #4967; 1:1000). Membranes were washed (3 times in 1X TBS-T) and incubated (2h; RT) with the peroxidase-conjugated secondary antibody (HRP conjugated goat anti-rabbit IgGH+L; Invitrogen, #31460; 1:5000; 5% milk in 1X TBS-T). Membranes were washed (3 times in 1X TBS-T) and peroxidase activity was detected using SuperSignal™ West Pico PLUS Chemiluminescent Substrate (ThermoFisher Scientific). Blots were developed using Amersham Imager 680 (GE Healthcare), equipped with a Peltier cooled Fujifilm Super CCD. Western blot analysis was performed using ImageJ (Rasband, W.S., ImageJ, U.S. NIH, Bethesda, Maryland, USA, https://imagej.nih.gov/ij/,1997-2014), from images without saturated pixels. Uncropped Western blot membranes are shown in **Data S3**. For hepatic UGT1A1 detection the procedure was modified as follows: tissues were lysed using radioimmunoprecipitation assay (RIPA) buffer (pH 7.5) (50mM TrisHCl, 1% NP40, 0,25% deoxycholic acid, 150mM NaCl, 1mM EGTA, 1mM Sodium Orthovanadate, 1mM Sodium Fluoride and 1X protease inhibitor cocktail (cOmplete™, Mini, EDTA-free Protease Inhibitor Cocktail; Roche) and homogenized in a TissueLyser II (Qiagen) with tungsten carbide beads (Qiagen). Samples were centrifuged (15000 g 15min., 4ºC), the supernatants were collected, and the total protein concentration was quantified by the Bradford assay, according to the manufacturer instructions (Bio-rad Protein Assay Dye Reagent Concentrate, #5000006). Protein (40µg) were resolved on a 10% SDS-PAGE and transferred to Polyvinylidene fluoride (PVDF) membrane. The membrane was blocked (2h at RT; 5% milk in 1X PBS-T) and incubated with rabbit polyclonal anti-UGT1A1 (Boster Biological Technology, A01865-1; 1:1000). The membrane was washed (3 times in 1X-PBS-T) and incubated (1h, RT) with the peroxidase-conjugated secondary antibody (HRP conjugated goat anti-rabbit IgGH+L; Invitrogen, #31460; 1:5000; 5% milk in 1X PBS-T). The membrane was washed (3 times in 1X-PBS-T) and the peroxidase activity was detected as described above.

*High-pressure freezing (HPF), freeze substitution and Transmission Electron Microscopy*

*Pf*3D7-GFP cultures were synchronized as described (*18*). Trophozoite stage parasites (approximately 18-20h after invasion, 4-5% parasitemia) were treated with vehicle (0.35% vol/vol

DMSO) or bilirubin (41 μM) or biliverdin (120 μM) for 8 or 12h, collected and fixed overnight at 4ºC with 1% (v/v) glutaraldehyde (Science Services) in 0.1M cacodylate buffer (Sigma-Aldrich) with 1 mM $CaCl_2$ (Alfa Aesar) and 3 mM $MgCl_2$ (Alfa Aesar). Parasites were pelleted by centrifugation (300 *g*; 5 min.), washed 3 times in 0.1M cacodylate buffer (Sigma-Aldrich) (pH 7.2), mixed in RPMI medium containing 20% (w/v) polyvinylpyrrolidone 40 (Sigma-Aldrich) as a cryoprotectant and high-pressure-frozen using a High-Pressure Freezer Compact 02 (Wohlwend Engineering Switzerland). The samples were freeze-substituted using a Leica AFS2 equipped with a Leica EM FSP Robot with 0.2% (w/v) uranyl acetate (Analar) in acetone during 1h at -140ºC, followed by an increasing slope of 4ºC/h until -90ºC and then samples were substituted for 6h at -90°C. The temperature was raised to -50ºC at a slope of 5ºC/h and the samples were washed three times in acetone for 1h each ste*P*. Samples were then infiltrated in Lowicryl HM20 resin (Polysciences) at increasing concentrations of 33, 66 and 100% and polymerized with UV light at -30ºC for 72h after a slope of 5ºC/h. Sections of 70 nm thickness were cut using a Leica EM UC7 ultramicrotome using an Ultra 45° diamond knife (Diatome) and mounted on palladium-copper 1x2mm slot grids coated with 1% (w/v) formvar (Agar Scientific) in chloroform (VWR). Sections were stained with 2% (w/v) uranyl acetate (Analar) in 70% methanol (VWR) and Reynold's lead citrate (Sigma-Aldrich; 5min. each) and analyzed using a Tecnai $G^2$ Spirit BioTWIN Transmission Electron Microscope from FEI operating at 120 kV and equipped with an Olympus-SIS Veleta CCD Camera.

*Scanning Electron Microscopy*
Vehicle (0.35% vol/vol DMSO) or bilirubin (41 μM) was added to the culture medium used to culture *Pf*D7-GFP for 12h. The samples were collected, centrifuged (500 *g*, 5 min) and washed (3X in 100 mM $NaHCO_3$, pH 9.0 and 2% SDS; 11,000 *g*, 30 min; and 5X in distilled H2O to remove the salts and detergents (4 times 11,000 *g*,10 min and the last wash 11,000 *g*, 30 min). Samples were stored in 200 μL of distilled $H_2O$ at 4ºC until processing. 20 μL of each sample were added to a silicon wafer and allowed to air dry. Before analysis, samples were coated with gold for 5 seconds. Samples were imaged in a Quanta 650 FEG Scanning Electron Microscope from FEI operating at 5 kV.

*Single-cell RNA sequencing*
Single-cell RNA sequencing was performed essentially as described (*7, 19*). Briefly, *Blvra*$^{+/+}$ and *Blvra*$^{-/-}$ mice were sacrificed by $CO_2$ asphyxiation 7 days after *Pcc* AS-GFP infection (*i.e.,* peak of infection) and the blood was collected by cardiac puncture. The blood was diluted 1:75 in 1X PBS supplemented with 1 % FBS and iRBC ($GFP^+$ cells) were sorted on a BD FACSAria Ilu. Cells were washed twice with 1X PBS supplemented with 1 % FBS and adjusted to a concentration of 1200 cells/mL.

*Single-cell gene expression library preparation and sequencing*
Library preparation and sequencing was performed essentially as previously described (*7, 19*). Briefly, samples were processed with Chromium Single Cell Controller to generate barcoded single-cell gel bead emulsions (GEMs) following the Chromium Next GEM Single Cell 3′ protocol. Single-cell cDNA libraries were obtained after the GEM-RT (reverse-transcriptase) clean-up and cDNA amplification on a Bio-Rad C1000 Touch Thermal Cycler. The cDNA profiles were checked on a Fragment Analyzer System (Agilent Technologies) according to the HS NGS Fragment kit (Agilent Technologies) manual. The scRNA-seq libraries were generated and

indexed (Dual Index Plate TT, Set A) on a Bio-Rad C1000 Touch Thermal Cycler. The final 3' Gene Expression libraries were verified and quantified on a Fragment Analyzer System (Agilent Technologies) following the HS NGS Fragment kit (Agilent Technologies) and samples were sequenced on a NextSeq 2000 (Illumina) using the NextSeq 1000/2000 P2 kit 100 cycles (Read 1: 28 Cycles; Read 2: 90 cycles).

*Single-cell RNA-seq data analysis*

Single-cell analyses were performed with Seurat (v.4.3.0) (*20-22*) and scuttle (v1.2.1) (*23*). Cells expressing less than 200, more than 1,500 genes, or more than 4,000 UMIs were filtered out. Remaining cells were annotated by projection to the malaria cell atlas, *P. falciparum* 10x set 4 (https://www.malariacellatlas.org/data-sets/) (*24*) using scmap (v1.14.0) (*25*) through 1:1 orthologs downloaded on 18.01.2024 from PlasmoDB (https://plasmodb.org/plasmo/app/search/transcript/GenesByOrthologPattern). The atlas and query samples were subset to the 1:1 ortholog gene set and log1p normalized. A cell index of the atlas was built using 500 genes and the cells from the query samples were individually projected using the scmapCell() function. The *Blvra*$^{+/+}$ and *Blvra*$^{-/-}$ samples were individually normalized with scran (v1.20.1) (*26*) using the quickCluster() method, and integrated using STACAS (v2.0.1) (*27*) with 750 genes and semi-supervised by the scmap projected stages. The integrated data was visualized by projection into UMAP (Uniform Manifold Approximation and Projection) using 8 principal components and the umap-learn method through Seurat. Analysis of differentially expressed genes of *Blvra*$^{-/-}$ *vs*. *Blvra*$^{+/+}$ was performed individually for different stage populations based on the scmap projection, using the Wilcoxon rank sum test through Seurat. Functional enrichment analysis of significantly (Bonferroni-adjusted $p<0.05$) up- or down-regulated genes was performed using gprofiler2 (v.0.2.2) (*28*). The archived version of the gprofiler2 server - Ensembl 108, Ensembl Genomes 55 (database built on 2022-12-28) was used: https://biit.cs.ut.ee/gprofiler_archive3/e108_eg55_p17/gost. All the analyses were performed with the R programming language in a containerized docker image publicly available at Docker Hub (*elolabfi/sctoolkit*) running R (v.4.2.1) (R Core Team, 2021) (*29*), RStudio server (2022.07.2 Build 576), Seurat (v.4.3.0) and its dependencies. RNA velocity was performed with velocyto (v.0.17.17) (*30*) and scvelo (v.0.2.5) (*31*) as described in (Ref. (*19*)) with python (v.3.9.5) run under Jupyter lab (v.3.5.3). All the data is available in **Tables S1-4**.

*Live cell imaging and confocal microscopy*

*Pf*3D7 cultures were synchronized, as described above and ring (10-12h after RBC invasion) or trophozoite (18-20h after invasion) stage parasites (3% parasitemia) were incubated with vehicle (0.35% vol/vol DMSO), unconjugated bilirubin (41 µM) or biliverdin (120 µM). RBC were harvested 12h or 24h for food vacuole and Hz or mitochondrion analyses, respectively. Cells were washed (1X PBS; 400 *g*, 2 min.) and stained (30 min., RT) with MitoTracker™ Green FM (200 nM in 1X PBS; Invitrogen™) or LysoTracker™ Green DND-26 (75 nM in 1X PBS; Invitrogen™), Hoechst 33342 (20 µM in 1X PBS) and Wheat Germ Agglutinin (WGA) Alexa Fluor 633 (3 µg/mL in 1X PBS; Thermo Fisher Scientific). Cells were washed twice (1X PBS; 400 *g*, 2 min.) and the cell pellet was resuspended in 1 mL of PBS. Then, 200 µL of the cell suspension were placed in chambered coverslip with 8 wells and a #1.5H glass bottom (Ibidi GmbH, Munich, Germany). After this step, the cells were analyzed using a laser scanning confocal microscope (Leica TCS-SP5) with a continuous Ar-ion and HeNe and a Ti:sapphire laser (Spectra-Physics Mai Tai BB, 710–990 nm, 100 fs, 82 MHz). Both MitoTracker™ Green FM and LysoTracker™ Green

DND-26 were imaged using the 488 nm Ar+ laser line (with emission set at 500-610 nm) and the 633 nm He−Ne laser line was used for imaging WGA-Alexa 633 dye (emission 644-736 nm). Hoechst 33342 images were recorded in the multiphoton mode under 810 nm excitation (420-550 nm), and hemozoin was visualized using the laser reflection mode. Images (512 × 512 pixels) were collected using a 63x 1.2 N.A. water immersion objective (HCX PL APO CS 63.0x 1.20WATERUV) at a scan rate of 100 Hz laser. To measure the relative area and fluorescence intensity corresponding to LysoTracker Green (*i.e.*, food vacuole) or Hz, a line was drawn across the diameter of *P. falciparum* 3D7 infected RBC, intercepting the food vacuole, using ImageJ 1.53k software (Rasband, W.S., ImageJ, U.S. NIH, Bethesda, Maryland, USA, https://imagej.nih.gov/ij/,1997-2014). Relative Intensity and the distance were measured using ImageJ. For LysoTracker Green, Area under the curve (AUC) was calculated from the above values. Values were calculated from 2 independent experiments analyzing 5-7 parasites *per* experiment. To measure the relative area corresponding to MitoTracker Green staining, images were analyzed using Imaris 10.0.0 software. The surface around the MitoTracker Green was created using the Surface Tool in Imaris and the area *per* parasite was measured. Values were calculated from 2 independent experiments analyzing 15-17 parasites *per* experiment.

*Hz quantification*

Hz was quantified essentially as described (*32, 33*), with the following adaptations. Briefly, blood was obtained by heart puncture, from *Blvra*$^{+/+}$ and *Blvra*$^{-/-}$ mice 7 days after *Pcc* infection. Mice were housed at a regular light cycle: lights on from 8:00 am (Zeitgeber time: ZT 0) to 8:00 pm (ZT 12), or at an inverted light cycle: lights on from 8:00 pm (ZT 0) to 8:00 am (ZT 12). Blood (300-400 µL) was collected at ZT 3 and 15, hypotonically lysed (4 mL $H_2O$), centrifuged (11,000 *g*, 45 min.). The supernatant fraction was removed, and pellets (Hz) were washed (3X in 100 mM $NaHCO_3$, pH 9.0 and 2% SDS; 11,000 *g*, 30 min.). Hz was dissolved (1 mL of 100 mM NaOH, 2% SDS, 3 mM EDTA), sonicated (1 min. Branson SLPe Digital Sonifier) and centrifuged (11,000 *g*, 30 min.). The heme released from Hz (200 µL) was quantitated by spectrophotometry (405nm) in a 96 well plate reader (MultiscanSky, ThermoFisher) with molar extinction coefficient of 5.7 × $10^4$. Heme concentration was calculated according to the Lambert-Beer law: A=ε.c.l (A – absorbance at $l_{405}$ nm; ε – extinction coefficient of hemin- 57000; c – concentration in molar (M); l – path length) and normalized to the volume of blood used. Data is presented as Hz (*i.e.*, nM heme).

*β-Hematin inhibition assay*

The Nonidet P-40 (NP-40) detergent-mediated assay (*34*) adapted for high-throughput screening in 96-well plate was used to determine inhibition of β-hematin formation. Following a period of 4-5h incubation at 37°C, the formation of a *bis*-pyridyl hemochromogen is detectable at a wavelength of 405 nm, and allows quantification of the free heme component that has not reacted to form β-hematin (*35*). Measurements were made using a Thermo Scientific Multiskan GO plate reader. Sigmoidal dose-response curves were plotted in GraphPad Prism to determine half maximal inhibitory concentration ($IC_{50}$) values. Measurements were performed twice, each in technical duplicate, and the values are reported together with standard deviation.

*Metabolite extraction*

*Pf*3D7-GFP cultures were synchronized as described (*18*). Trophozoite stage parasites (approximately 18-20h after invasion, 15% parasitemia, 5% HCT, 2 mL culture *per* sample) and treated with vehicle (0.35% vol/vol DMSO) or bilirubin (41 μM) for 8 or 12h. The chemicals used were LC-MS grade water, acetonitrile (ACN), methanol (MeOH), and isopropanol (IPA), which were obtained from Th. Geyer (Germany). High-purity methyl tert-butyl ether (MTBE), ammonium formate, formic acid, ammonium acetate, and acetic acid were purchased from Merck (Germany). Stable isotope labelled internal standards for metabolomics (MSK-A2-1.2; Cambridge Isotope Laboratories, MA, USA) were used at final concentrations of 1.0% (vol/vol). The culture medium was completely aspirated and the cells were washed with 1XPBS (500 *g*, 5 min). For biphasic extraction of lipids and polar metabolites, samples were initially quenched by incubation on dry ice with 400 μL of 75% (vol/vol) cold methanol for 20 min and the appropriate internal standards (0.5 μL each/sample) were added. After incubation, the samples were vortexed for 5 min at maximum speed, lysed using an ultrasonic bath to sonicate the sample for 5 min and vortexed again briefly after sonication. After addition of 1000 μL of cold MTBE, the monophasic mixture was vortexed for 60 s and incubated at -20°C for 20 min. For phase separation, 250 μL of cold water were added, followed by another vortexing and incubation step (see previous conditions). The biphasic solvent system was then centrifuged for 15 min at 14,000 *g* and 4 °C. For metabolomics analysis, 400 μL of the bottom aqueous phase were transferred, dried under a stream of nitrogen, and reconstituted in 75 μL 80% MeOH (v/v). The final samples were vortexed for 10 min, centrifuged (see previous conditions) and the supernatants were transferred to analytical glass vials for LC-MS/MS analysis.

*LC-MS/MS analysis*

LC-MS/MS analysis was performed on a Vanquish Horizon UHPLC system coupled to an Orbitrap Exploris 240 high-resolution mass spectrometer (Thermo Scientific, MA, USA) in negative and positive ESI (electrospray ionization) mode. To perform untargeted metabolomics, chromatographic separation was carried out on an Atlantis Premier BEH Z-HILIC column (Waters, MA, USA; 2.1 mm x 100 mm, 1.7 μm) at a flow rate of 0.25 mL/min. The mobile phase consisted of water:acetonitrile (9:1, v/v; mobile phase phase A) and acetonitrile:water (9:1, v/v; mobile phase B), which were modified with a total buffer concentration of 10 mM ammonium acetate (negative mode) and 10 mM ammonium formate (positive mode), respectively. The aqueous portion of each mobile phase was pH-adjusted (negative mode: pH 9.0 via addition of ammonium hydroxide; positive mode: pH 3.0 via addition of formic acid). The following gradient (20 min total run time including re-equilibration) was applied (time [min]/%B): 0/95, 2/95, 14.5/60, 16/60, 16.5/95, 20/95. Column temperature was maintained at 40°C, the autosampler was set to 4°C and sample injection volume was 5 μL. Analytes were recorded via a full scan with a mass resolving power of 120,000 over a mass range from 60 – 900 m/z (scan time: 100 ms, RF lens: 70%). To obtain MS/MS fragment spectra, data-dependent acquisition was carried out (resolving power: 15,000; scan time: 22 ms; stepped collision energies [%]: 30/50/70; cycle time: 900 ms). Ion source parameters were set to the following values: spray voltage: 4100 V (positive mode) / -3500 V (negative mode), sheath gas: 30 psi, auxiliary gas: 5 psi, sweep gas: 0 psi, ion transfer tube temperature: 350°C, vaporizer temperature: 300°C. All experimental samples were measured in a randomized manner. Pooled quality control (QC) samples were prepared by mixing equal aliquots from each processed sample. Multiple QCs were injected at the beginning of the analysis in order to equilibrate the analytical system. A QC sample was analyzed after every 5th experimental sample to monitor instrument performance throughout the sequence. For determination of background signals and subsequent background subtraction, an additional processed blank sample was recorded. Data was processed using MS DIAL 4.9.221218 (*36*) and raw peak intensity data was normalized via total ion count of all detected analytes (*37*). Level 1 feature identification was based on an in-house library for metabolomics (EMBL-MCF 2) (*38*) using accurate mass, isotope pattern, MS/MS fragmentation, and retention time information and a minimum matching score of 80%. All the raw data results for annotated metabolites are available in **Table S5.**

*Statistical analysis*

For clinical data we used Mann Whitney U test. Analysis was done in JMP Pro version 16 (SAS Institute Inc, Cary, NC, USA). For all other analyses, statistically significant differences between two experimental groups were assessed using a two-tailed unpaired Mann-Whitney U test or t test (in samples tested positive for normal distribution) and comparisons between more than two groups were assessed using one-way ANOVA, two-way ANOVA (parasitemias and parasite burden) or two-way ANOVA with Bonferroni's or Tukey's multiple comparison test. Survival curves are represented by Kaplan-Meier plots and differences between the groups were assessed using the log-rank test. All statistical analyses were performed using GraphPad Prism software. Differences were considered statistically significant at a $P$ value <0.05. NS: not-significant, $p$>0.05; *$p$<0.05; **$p$<0.01; ***$p$<0.001; ****$p$<0.0001.

For *in vivo Plasmodium* infection experiments, sample size was maintained no less than 5 mice *per* experimental condition, in at least 2 independent experiments, except when indicated in the Figure Legends. For *in vitro* experiments, sample size was maintained no less than 4 replicates *per* experimental condition, in a minimum of 3 independent experiments, except when indicated in the Figure Legends.

Randomization was used in mice monitoring after infection. Samples were analyzed blindly, often being processed by a different investigator from the one preparing/collecting the samples.

Inclusion/exclusion criteria for human studies are detailed in the *Human data* section. For in vivo *Plasmodium* infection experiments, control mice presenting a >2 days delay in the initial dynamics of parasite growth and peak of parasitemia were considered as technical outliers and were excluded from further analysis.

#### Supplementary Figures

*Ana Figueiredo et al. Fig.S1*

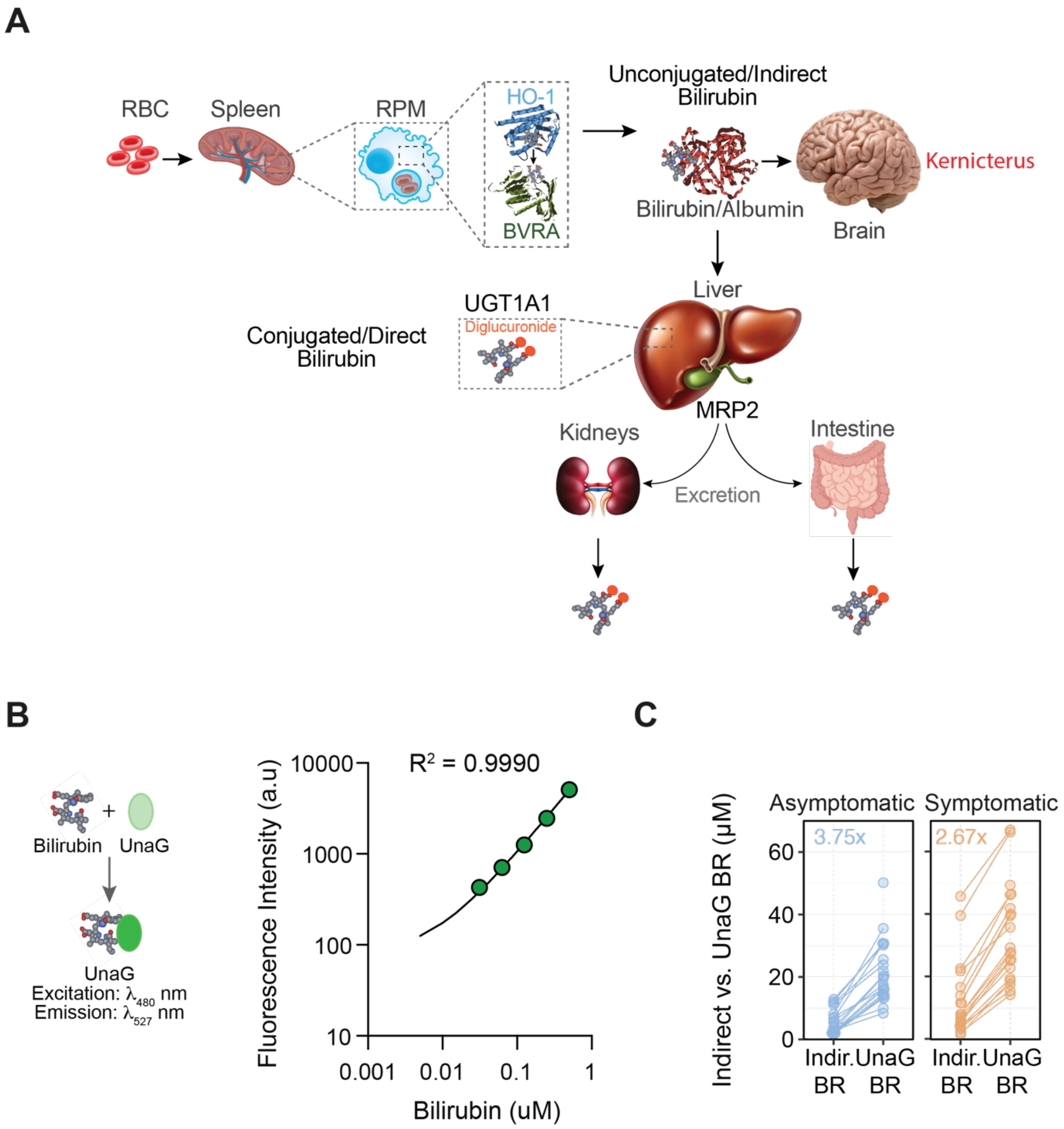

**Fig. S1. A UnaG-based assay for quantification of unconjugated bilirubin in plasma.** (**A**) Schematic illustration of systemic bilirubin metabolism. Senescent and damaged red blood cells (RBC) are phagocytosed in the spleen by red pulp macrophages (RPM), which catabolize the prosthetic heme groups of hemoglobin via heme oxygenase-1 (HO-1). Heme catabolism generates biliverdin, which is converted into bilirubin by biliverdin reductase (BVRA) in macrophages. Unconjugated bilirubin is released from macrophages and circulates in plasma bound to albumin (*i.e.,* indirect bilirubin) to reach hepatocytes where it is conjugated to UDP-glucuronic or glucuronic acid, via a reaction catalyzed by UDP-glucuronosyl transferase 1A1 (UGT1A1). Conjugated (*i.e.,* direct) bilirubin is transported to the bile via the multidrug resistance protein 2 (MRP2) and is then excreted via the bile or by the kidneys upon liver damage. Partial deficiency of UGT1A1 results in increased concentration of unconjugated bilirubin in plasma, that can reach neurotoxic levels in newborns and in patients suffering from inherited UGT1A1 deficiency causing permanent brain damage known as kernicterus which may lead to death. (**B**) Linear range of the standard curve of purified unconjugated bilirubin in the UnaG-based assay. Fluorescence intensity was measured at excitation at $\lambda_{480nm}$ and emission at $\lambda_{527nm}$. (**C**) Comparison of unconjugated bilirubin concentration measured by the Roche Diazo method (indirect bilirubin; BR) and by the UnaG-based assay (UnaG BR) (*2, 3*) in the plasma of the same *P. falciparum*-infected patients, as in Figure 1A. Circles represent individual patients. Values inside the plots indicate the fold difference of UnaG BR over Indir. BR.

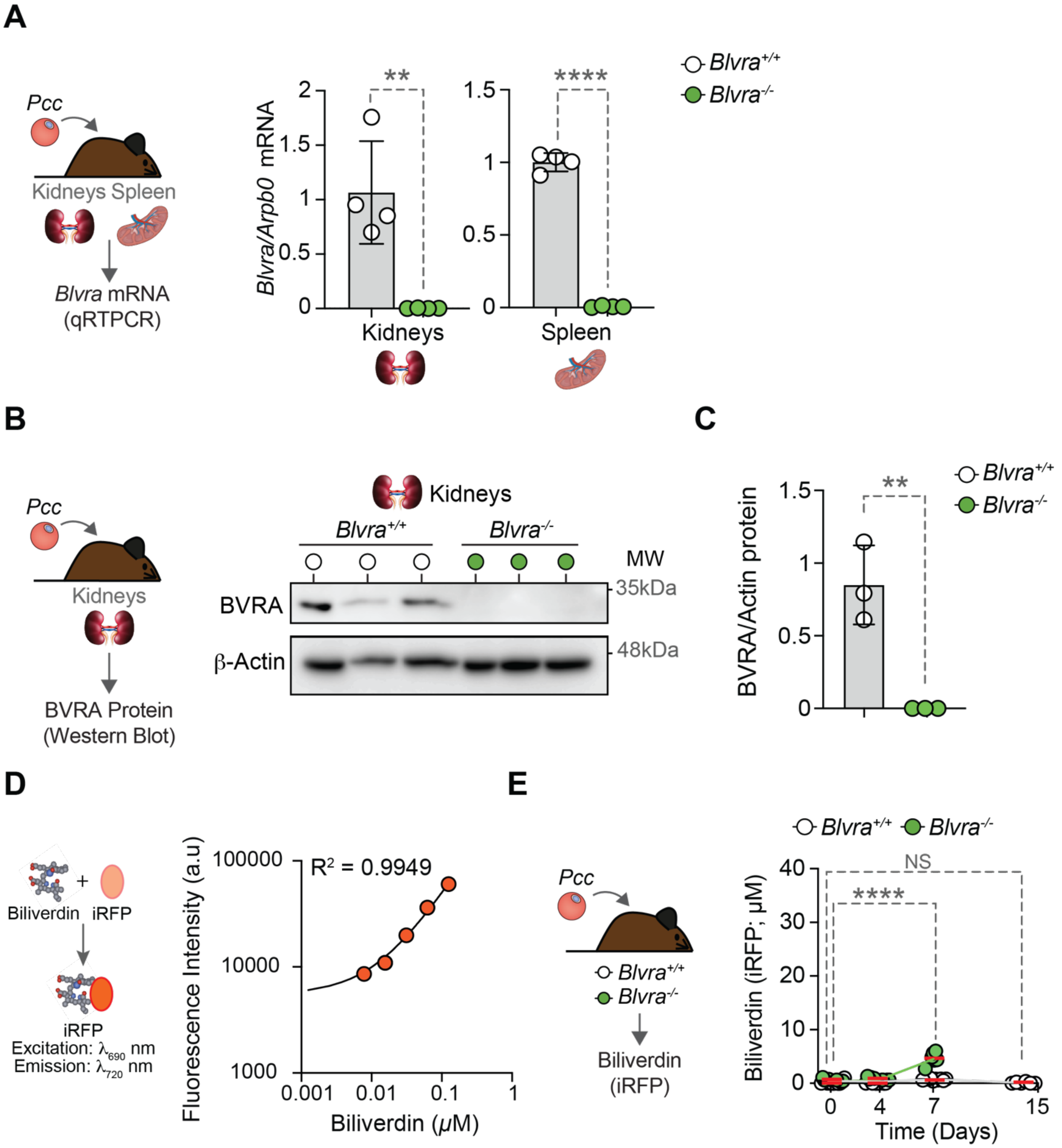
A
Pcc
Kidneys Spleen
Blvra mRNA
(qRTPCR)
Blvra/Arpb0 mRNA
**
****
Kidneys
Spleen
Blvra+/+
Blvra-/-
B
Pcc
Kidneys
BVRA Protein
(Western Blot)
Kidneys
Blvra+/+
Blvra-/-
MW
BVRA
β-Actin
35kDa
48kDa
C
BVRA/Actin protein
**
Blvra+/+
Blvra-/-
D
Biliverdin
iRFP
iRFP
Excitation: λ690 nm
Emission: λ720 nm
Fluorescence Intensity (a.u)
R2 = 0.9949
Biliverdin (μM)
E
Pcc
Blvra+/+
Blvra-/-
Biliverdin
(iRFP)
Blvra+/+ Blvra-/-
NS
****
Biliverdin (iRFP; μM)
Time (Days)

**Fig. S2. Residual accumulation of circulating biliverdin in *Blvra*-deleted mice.** (**A**) Relative levels of *Blvra* mRNA, normalized to *Arbpo* mRNA, quantified by qRT-PCR in the kidneys and spleen of *Blvra*-deleted (*Blvra*$^{-/-}$) and control *Blvra*$^{+/+}$ mice, at steady state conditions. Data represented as mean ± SD (n=4 *per* genotype), pooled from two independent experiments with similar trend. (**B**) Western blot and (**C**) relative quantification of BVRA protein, normalized to ß-Actin, protein expression, in the kidneys of *Blvra*$^{-/-}$ and control *Blvra*$^{+/+}$ mice, at steady state conditions. Data represented as mean ± SD (n=4 *per* genotype), from one experiment. (**D**) Linear range of the standard curve of purified biliverdin in the iRFP-based assay. Fluorescence intensity was measured at excitation at $\lambda_{690nm}$ and emission at $\lambda_{720nm}$. (**E**) Biliverdin concentration in plasma from *Blvra*$^{+/+}$ and *Blvra*$^{-/-}$ mice (n=6-9 *per* genotype), before (Day 0) and after (Days 4, 7 and 15) *Pcc* infection. Data represented as mean ± SD, pooled from two independent experiments with similar trend. Circles in (A, C, E) correspond to individual mice. *P* values determined using: (A, C) t test, (E) Two-Way ANOVA with Bonferroni's multiple comparison test (for genotypes) and Ordinary One-Way ANOVA with Tukey's multiple comparison (for days post-infection). NS: not significant; ***p*<0.01; *****p*<0.0001.

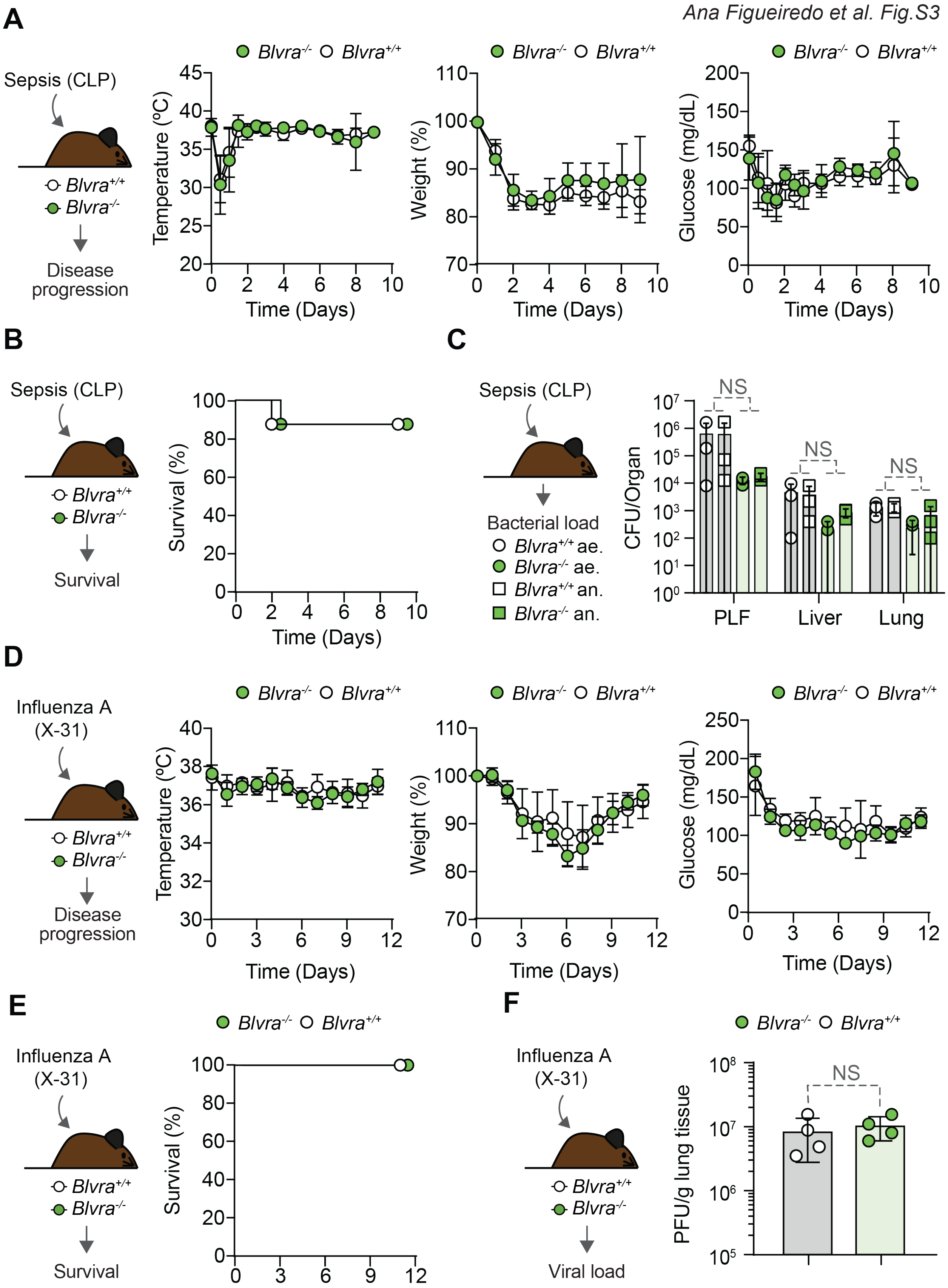
A
Sepsis (CLP)
Blvra+/+
Blvra-/-
Disease progression
Blvra-/- Blvra+/+
Temperature (ºC)
Time (Days)
Weight (%)
Glucose (mg/dL)
B
Sepsis (CLP)
Survival
Survival (%)
C
Sepsis (CLP)
Bacterial load
Blvra+/+ ae.
Blvra-/- ae.
Blvra+/+ an.
Blvra-/- an.
CFU/Organ
NS
PLF
Liver
Lung
D
Influenza A (X-31)
Disease progression
E
Influenza A (X-31)
Survival
F
Influenza A (X-31)
Viral load
PFU/g lung tissue
NS

**Fig. S3. BVRA is not essential to prevent the lethal outcome of non-hemolytic infections.** (**A**) Body temperature (*left panel*), percentage initial weight (*middle panel*) and blood glucose concentration (*right panel*) in *Blvra*$^{-/-}$ *vs. Blvra*$^{+/+}$ mice subjected to polymicrobial sepsis (cecal ligation and puncture; CLP). Data represented as mean ± SD from n=7-8 mice *per* genotype, pooled from two independent experiments with similar trend. (**B**) Survival of the same mice as in (A). (**C**) Bacterial (ae: aerobic; an: anaerobic) load, in different organs from *Blvra*$^{-/-}$ *vs. Blvra*$^{+/+}$ mice 24h after CLP. PLF: peritoneal lavage fluid. (**D**) Body temperature (*left panel*), percentage of initial weight (*middle panel*) and blood glucose concentration (*right panel*) in *Blvra*$^{-/-}$ *vs. Blvra*$^{+/+}$ mice during influenza A virus (X-31) infection. Data represented as mean ± SD from n=4 *per* genotype, from one experiment. (**E**) Survival of the same mice an in (D). (**F**) Viral load in lung tissue of *Blvra*$^{-/-}$ *vs. Blvra*$^{+/+}$ mice, 3 days after influenza A virus (X-31) infection. Circles in (C, F) represent individual mice. *P* values were determined using: (A, D) Two-Way ANOVA with Bonferroni's multiple comparison test, (B, E) Log-rank (Mantel-Cox) test, (C) Two-Way ANOVA with Tukey's multiple comparison test, (D) Mann-Whitney U test. NS: not significant.

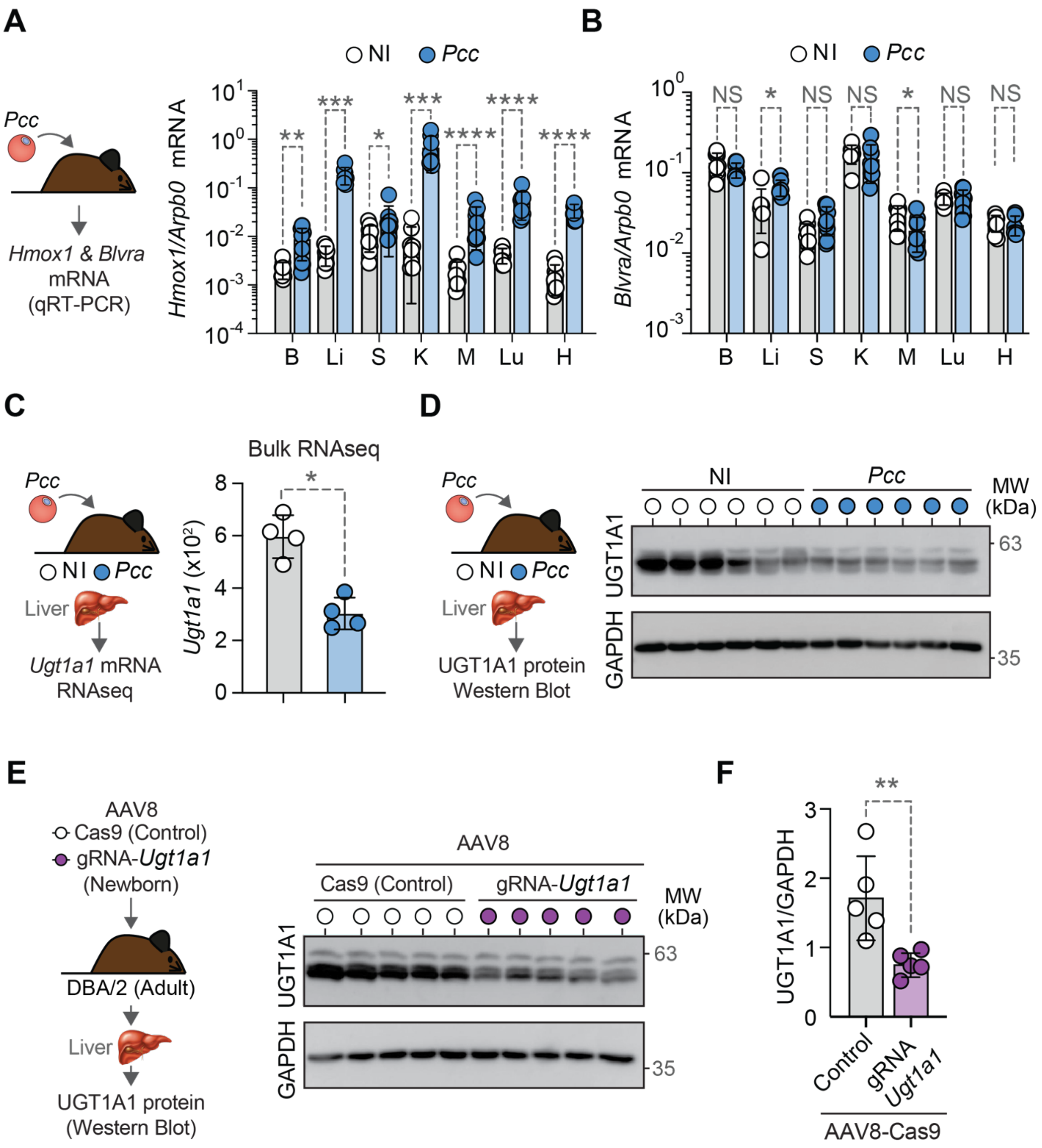
A
NI
Pcc
Pcc
Hmox1 & Blvra
mRNA
(qRT-PCR)
Hmox1/Arpb0 mRNA
B Li S K M Lu H
B
NI
Pcc
Blvra/Arpb0 mRNA
NS * NS NS * NS NS
B Li S K M Lu H
C
Pcc
NI Pcc
Liver
Ugt1a1 mRNA
RNAseq
Bulk RNAseq
Ugt1a1 (x10²)
D
Pcc
NI Pcc
Liver
UGT1A1 protein
Western Blot
NI
Pcc
MW
(kDa)
UGT1A1
GAPDH
63
35
E
AAV8
Cas9 (Control)
gRNA-Ugt1a1
(Newborn)
DBA/2 (Adult)
Liver
UGT1A1 protein
(Western Blot)
AAV8
Cas9 (Control)
gRNA-Ugt1a1
MW
(kDa)
UGT1A1
GAPDH
63
35
F
UGT1A1/GAPDH
Control
gRNA
Ugt1a1
AAV8-Cas9

**Fig. S4.** ***Hmox1*****,** ***Blvra*** **and** ***Ugt1a1*** **expression during** ***Pcc*** **infection.** (**A, B**) Relative level of (**A**) *Hmox1* and (**B**) *Blvra* mRNA, normalized to *Arbpo* mRNA expression in different organs of *Blvra*$^{+/+}$mice, not infected (NI) or 7 days after *Pcc* infection, quantified by qRT-PCR. B: Brain; Li: Liver; S: Spleen; K: Kidney; M: Muscle; Lu: Lung; H: Heart. Data represented as mean ± SD (n=3-5 *per* group), from one experiment. (**C**) Relative level of hepatic *Ugt1a1* mRNA expression from *Blvra*$^{+/+}$ mice, not infected (NI) or 7 days after *Pcc* infection, assessed by bulk RNA sequencing (*dataset from* (*7*)). (**D)** Detection of UGT1A1 and GAPDH protein expression by Western blot in the liver of *Blvra*$^{+/+}$ mice, not infected (NI) or 7 days after *Pcc* infection. (**E**) Detection and (**F**) relative quantification of UGT1A1 and GAPDH protein expression detected by Western blot, in the liver of adult DBA/2 mice, transduced 2-4 days after birth with recombinant AAV8 gRNA-*Ugt1a1* or control AAV8-Cas9. Data represented as mean ± SD (n=5 *per* genotype), from one experiment. Circles in (A-C,F) correspond to individual mice. *P* values determined using: (A, B) Mann-Whitney U test, (C,F) using t test. NS: not significant; *p<0.05; **p<0.01; ***p<0.001; ****p<0.0001.

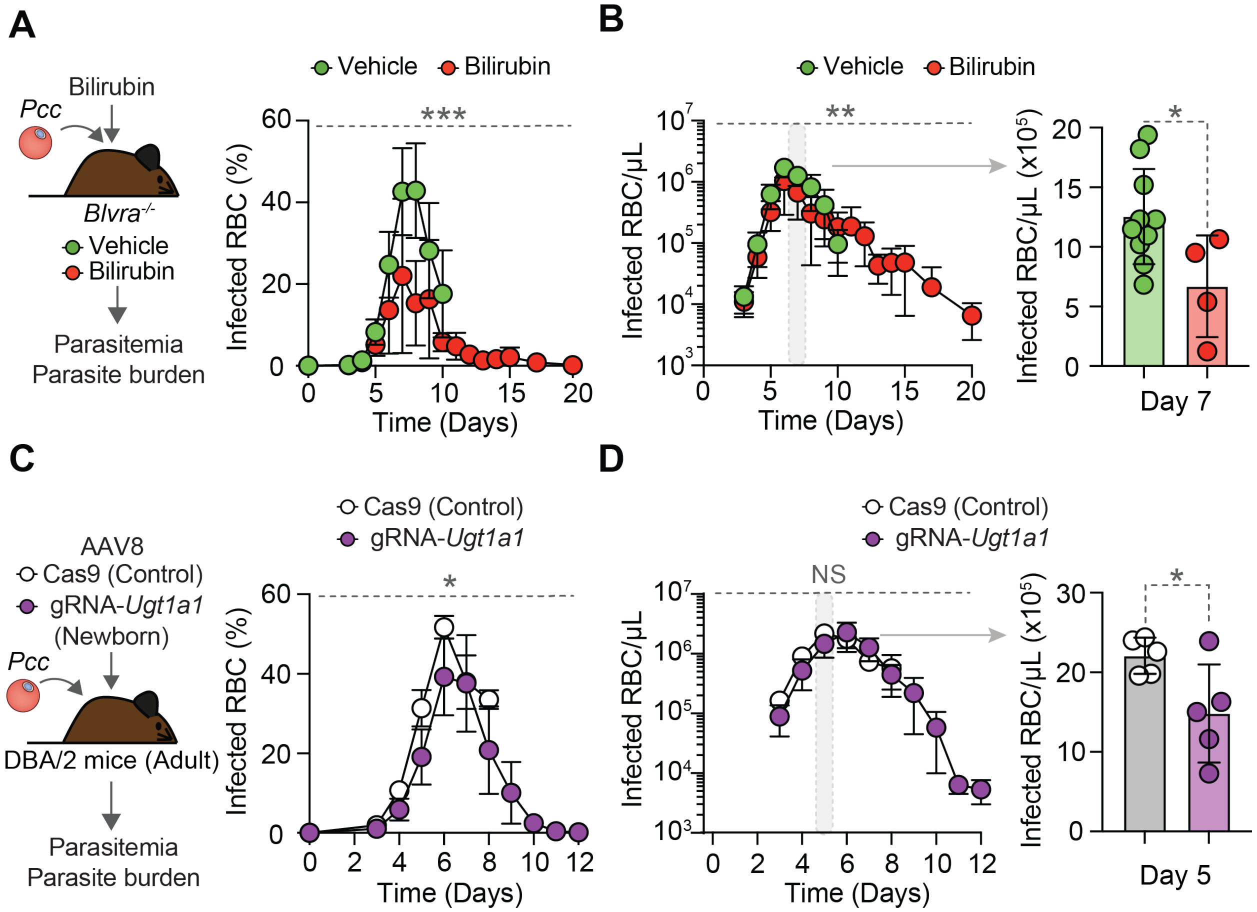


**Fig. S5. Unconjugated bilirubin reduces parasite burden *in vivo*. (A)** Percentage of iRBC (*i.e.,* parasitemia) and **(B)** number of iRBC (*i.e.,* parasite burden) in *Pcc*-infected *Blvra*$^{-/-}$ mice treated with unconjugated bilirubin (30 mg/kg; daily, i.p.) *vs.* controls receiving vehicle. *Right panel* highlights Day 7 from *left panel*. Data represented as mean ± SD from n=4-8 *per* group, from two independent experiments with similar trend. **(C)** Percentage of iRBC (*i.e.,* parasitemia) and **(D)** number of iRBC (*i.e.,* parasite burden) in *Pcc*-infected DBA/2 mice transduced with a AAV8-Cas9-gRNA-*Ugt1a1 vs.* control AAV8-Cas9, 2-4 days after birth. *Right panel* highlights Day 5 from *left panel*. Data represented as mean ± SD from n=5 *per* group, from one independent experiment, from three independent experiments with similar trend. Circles in (B *right panel* and D *right panel*) correspond to individual mice. *P* values were determined using: (A, C, B *left panel* and D *left panel*) Two-Way ANOVA, (B *right panel* and D *right panel*) t test. NS: not significant; *$p$<0.05; **$p$<0.01; ***$p$<0.001.

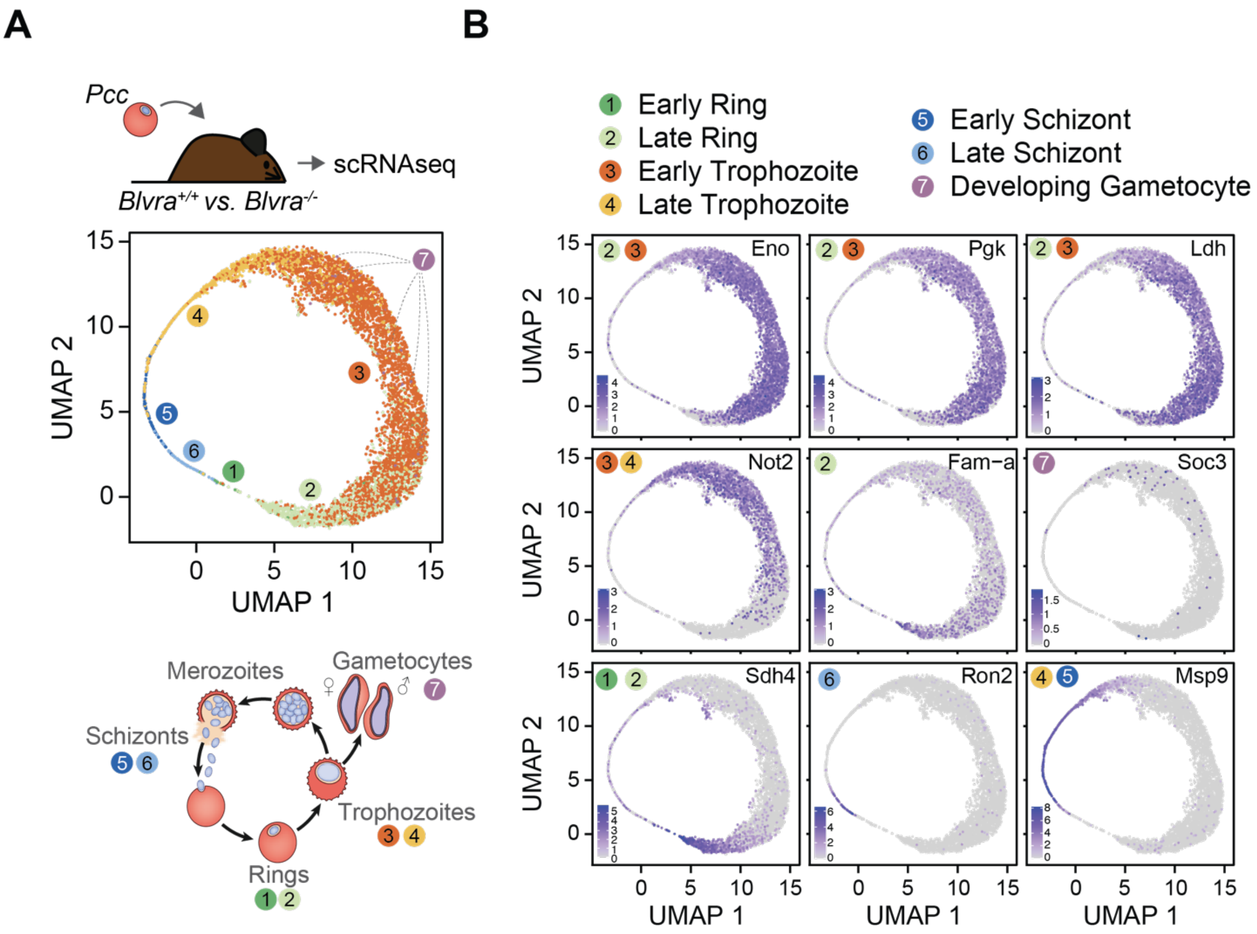


**Fig. S6. Bilirubin production by BVRA regulates the development of intraerythrocytic *Pcc in vivo*.** (**A**) Combined UMAP projection of single parasite transcriptomes of FACS-sorted circulating iRBC from *Blvra*$^{+/+}$ (n= 1862) and *Blvra*$^{-/-}$ (n= 3891) mice 7 days after *Pcc* infection. Colors and numbers identify different parasite developmental stages based on scmap projection to a *P. falciparum* atlas (**B**) Expression pattern of parasite stage-specific genes. In ring stages: enolase (*eno*, *PCHAS_1215000*) and succinate dehydrogenase subunit 4 (*sdh4*, *PCHAS_1209400*); in trophozoites: phosphoglycerate kinase (*pgk*, *PCHAS_0823700*) and *fam a* (*PCHAS_120120*); in schizonts: rhoptry neck protein 2 (*ron2*, *PCHAS_1319000*) and merozoite surface protein 9 (*msp9*, *PCHAS_1445500*), in developing gametocytes: carbon catabolite repression-negative on TATA-less, (CCR4-Not) subunit 2 (*not2*, *PCHAS_0924800*) and in gametocytes: SOC protein 3 (*soc3*, *PCHAS_1307700*).

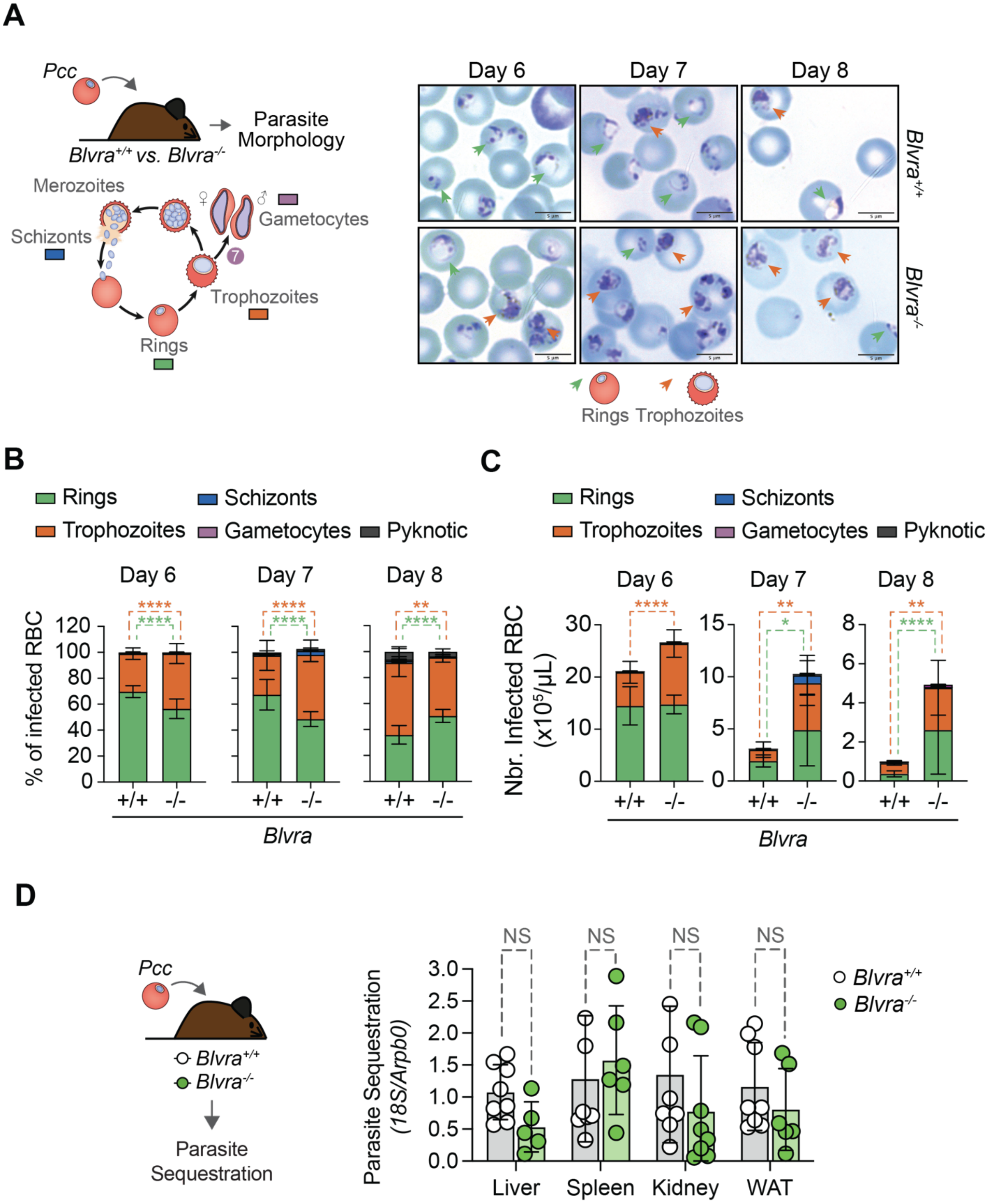

A
Pcc
Blvra+/+ vs. Blvra-/-
Parasite Morphology
Merozoites
Schizonts
Gametocytes
Trophozoites
Rings
Day 6
Day 7
Day 8
Blvra+/+
Blvra-/-
Rings
Trophozoites
B
Rings
Schizonts
Trophozoites
Gametocytes
Pyknotic
% of infected RBC
+/+
-/-
Blvra
C
Nbr. Infected RBC (x10^5/μL)
D
Parasite Sequestration
Parasite Sequestration (18S/Arpb0)
NS
Liver
Spleen
Kidney
WAT

**Fig. S7. Bilirubin production by BVRA regulates the development of intraerythrocytic *Pcc in vivo*.** (**A**) Representative Giemsa-stained thin blood smears from *Blvra*$^{+/+}$ and *Blvra*$^{-/-}$ mice at 6, 7 and 8 days after *Pcc* infection (n=5-10 per genotype). Green arrows highlight *Pcc* ring stage and orange arrows trophozoite stage parasites, identified based on morphology analysis. Scale bar is 5 µm. Data representative of three independent experiments with similar trend. (**B)** Percentage and (**C**) number of *Pcc* iRBC containing parasites at different developmental stages in the blood of the same mice as in (A). Data represented as mean ± SD, pooled from two or three independent experiments with similar trend. (**D**) Parasite sequestration (*Plasmodium* 18S rRNA/*Arbp0* mRNA) in liver, spleen, kidney, and white adipose tissue (WAT) of *Pcc*-infected *Blvra*$^{+/+}$ and *Blvra*$^{-/-}$ mice. Data from n=5-8 *per* genotype, pooled from two independent experiments with similar trend. In (D) circles represent individual mice. *P* values determined using: (B, C) Two-Way ANOVA with Bonferroni's multiple comparison test, (D) using Mann-Whitney U test. NS: not significant; *p<0.05; **p<0.01; ****p<0.0001.

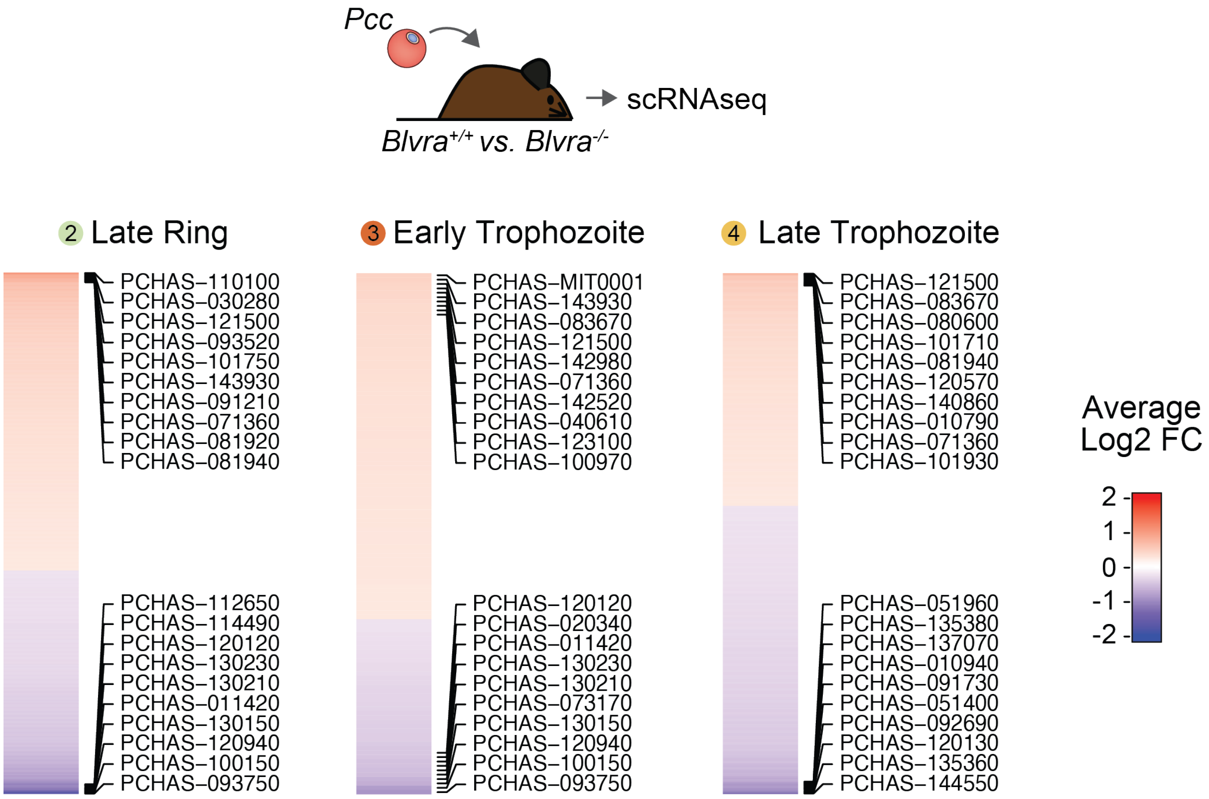


**Fig. S8. Bilirubin production by BVRA modulates *Pcc* blood stage gene expression.** Population-specific heatmap of differentially regulated genes (average log2 fold change) in late rings (Population 2), early (Population 3) and late (Population 4) trophozoites in FACS-sorted iRBC from *Blvra*$^{-/-}$ *vs*. *Blvra*$^{+/+}$ mice. A gene was considered differentially expressed (and plotted) if it has an absolute log2FC>0 and an adjusted $p$ value<0.05. R package ComplexHeatmap (v.2.2.0) was used (*39*). Same dataset as in Figure 2 (C-F) and Tables S1 and S2.

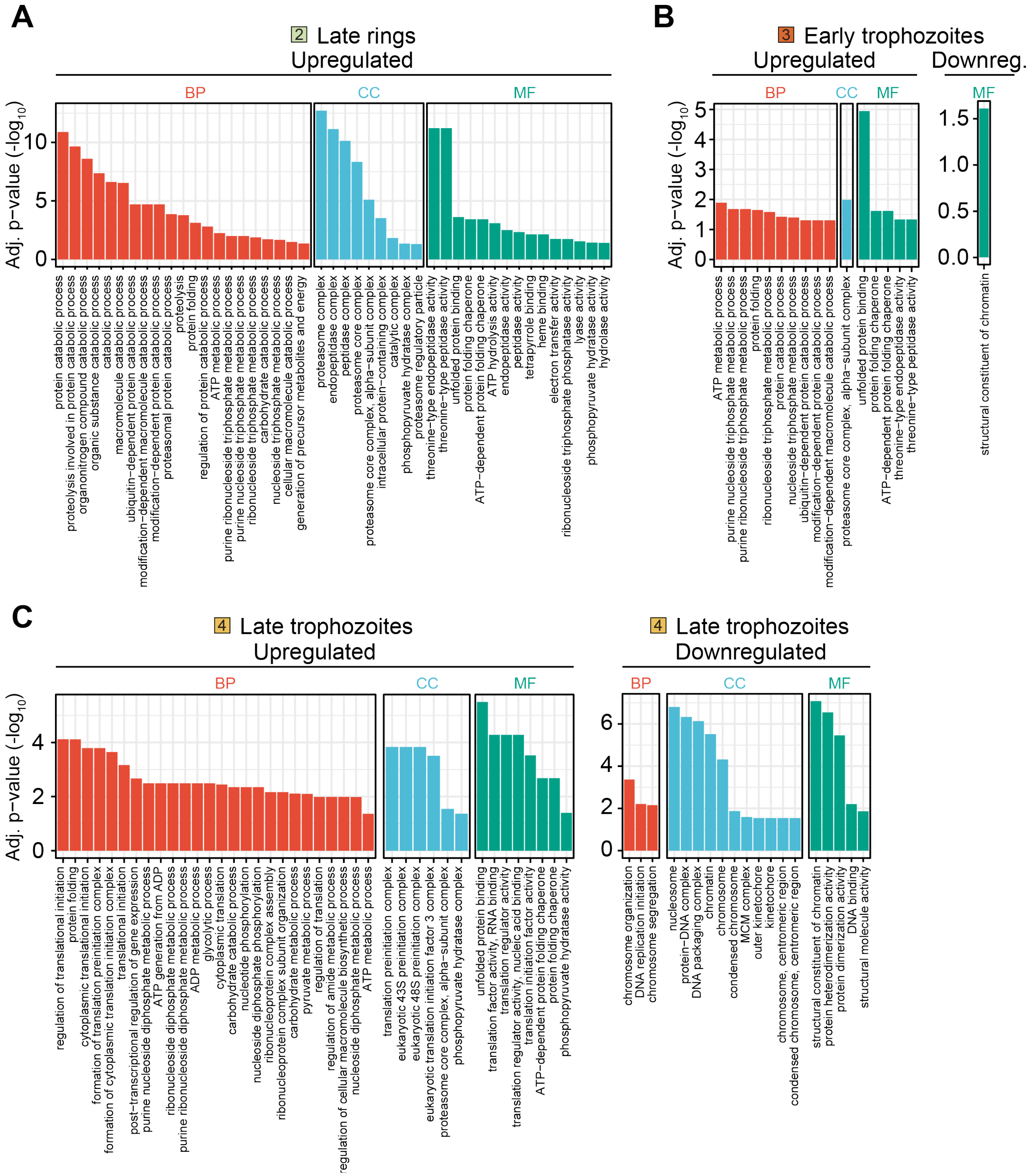


**Fig. S9. Bilirubin production by BVRA modulates *Pcc* blood stage gene expression.** (**A-C**) Functional enrichment analysis of parasite pathways (*i.e.,* GO terms database) (**A**) up-regulated in Population 2 (late rings parasites), (**B**) up- and down-regulated in Population 3 (early trophozoites parasites), and (**C**) up- and down-regulated in Population 4 (late trophozoites parasites), from the same FACS-sorted circulating iRBC in *Blvra*$^{-/-}$ *vs. Blvra*$^{+/+}$ mice as Figure 2 (C-F). See Table S3.

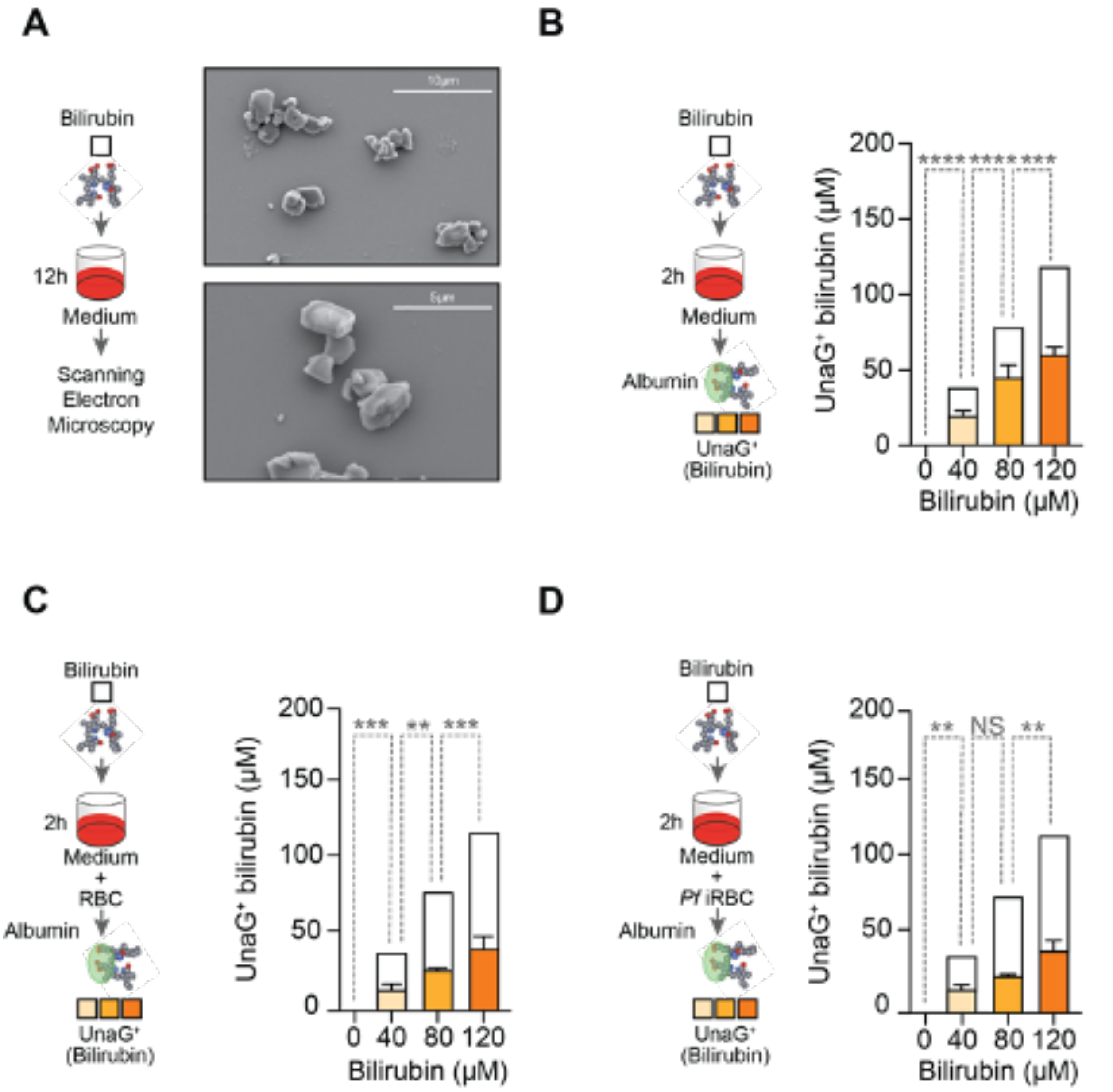


**Fig. S10. Quantification of unconjugated bilirubin concentration in culture medium.** (**A**) Representative images of scanning electron microscopy (SEM) showing bilirubin crystals 12h after the addition of 120 µM of unconjugated bilirubin to the culture medium used for *P. falciparum* proliferation assays (75µM albumin). Quantification of unconjugated bilirubin by the UnaG-based assay, in (**B**) culture medium (75µM albumin), (**C**) culture medium (75µM albumin) plus non-infected human RBC or (**D**) culture medium (75µM albumin) plus *P. falciparum* iRBC. In (B-D), white bars show total bilirubin added to the culture media, and colored bars the concentration of unconjugated bilirubin quantified in the culture medium by the UnaG-based assay (*2*), 2h after the addition of the unconjugated bilirubin. Data is represented as mean ± SD, from four replicates in one experiment. *P* values determined using: (B-D) Two-Way ANOVA with Tukey's multiple comparison test. NS: not significant; **$p<0.01$; ***$p<0.001$; ****$p<0.0001$.

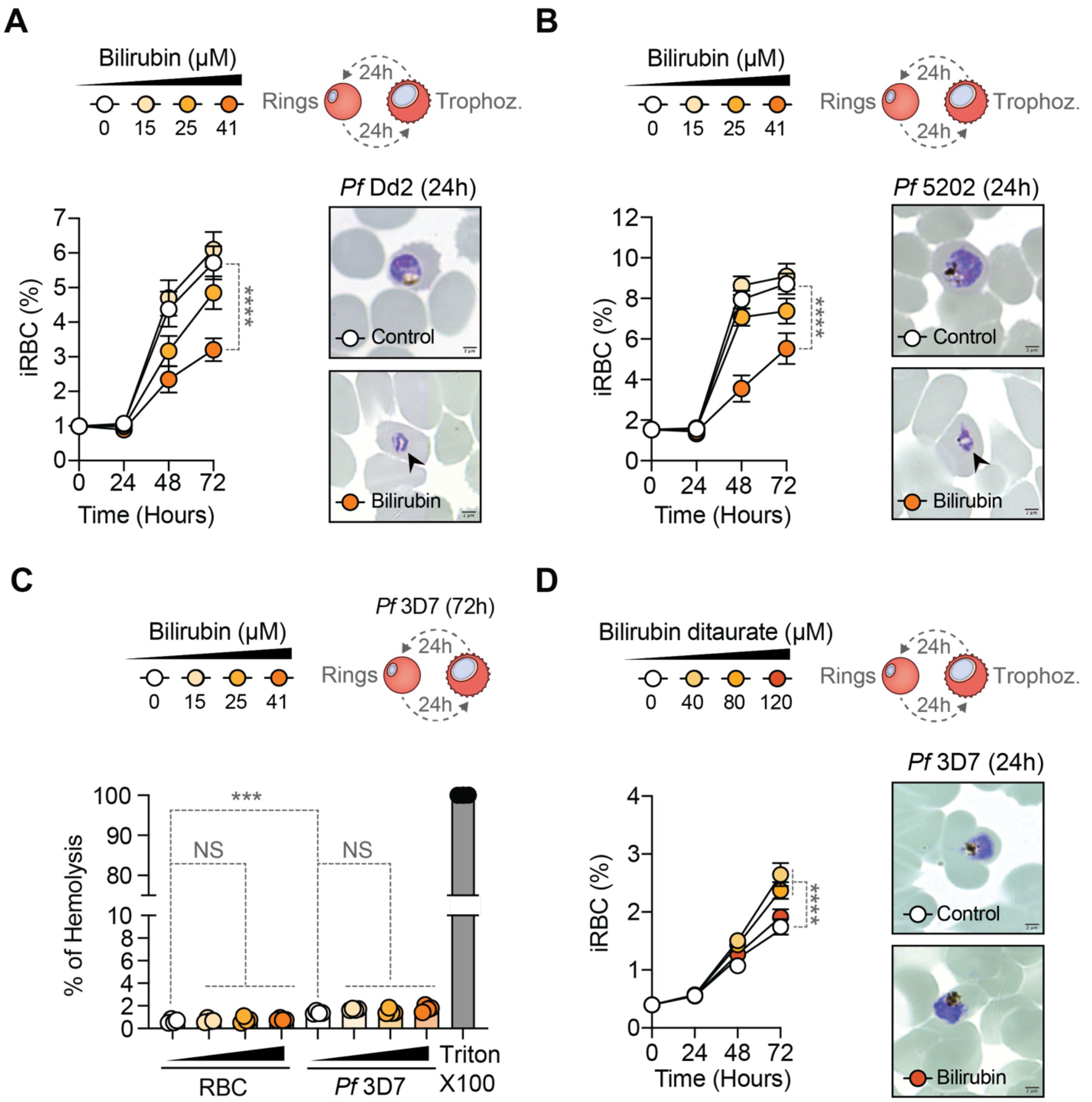
A
Bilirubin (μM)
0 15 25 41
Rings
Trophoz.
24h
24h
Pf Dd2 (24h)
iRBC (%)
Time (Hours)
****
Control
Bilirubin
B
Bilirubin (μM)
0 15 25 41
Rings
Trophoz.
24h
24h
Pf 5202 (24h)
iRBC (%)
Time (Hours)
****
Control
Bilirubin
C
Bilirubin (μM)
0 15 25 41
Pf 3D7 (72h)
Rings
24h
24h
% of Hemolysis
***
NS
NS
RBC
Pf 3D7
Triton X100
D
Bilirubin ditaurate (μM)
0 40 80 120
Rings
Trophoz.
24h
24h
Pf 3D7 (24h)
iRBC (%)
Time (Hours)
****
Control
Bilirubin

**Fig. S11. Unconjugated bilirubin arrests the proliferation of drug-resistant *P. falciparum* Dd2 and IPC 5202 ring stages.** (**A**) Percentage (*left panel*) of *P. falciparum* Dd2 (*Pf*Dd2) iRBC (rings), 24h after exposure to increasing concentrations of unconjugated bilirubin. *Right panel* is a representative Giemsa-stained thin smear 24h after exposure to unconjugated bilirubin (41 μM) or vehicle. Data represented as mean ± SEM, pooled from four independent experiments with similar trend, with four technical replicates *per* experiment. Black arrowhead highlights parasite nuclear DNA fragmentation. Scale bars: 2μm. (**B**) Percentage (*left panel*) of *P. falciparum* IPC 5202 (*Pf*5202) iRBC (rings), 24h after exposure to increasing concentrations of unconjugated bilirubin, as in (A). Data represented as mean ± SEM, pooled from three independent experiments with similar trend, with four technical replicates *per* experiment. *Right panel* is a representative Giemsa-stained thin smear 24h after exposure to unconjugated bilirubin (41 μM) or vehicle. (**C**) Hemolysis of uninfected RBC or *Pf3D7* iRBC, measured by the LDH assay, 72h after exposure to increasing concentrations of bilirubin or vehicle. Triton X-100 (2%) was used as positive control for 100% RBC lysis. Data represented as mean ± SEM, from four technical replicates in one experiment, representative of 3 assays performed at different time points. (**D**) Percentage (*left panel*) of *Pf3D7* iRBC (rings), 24h after exposure to increasing concentrations of water-soluble bilirubin ditaurate. Data represented as mean ± SEM, pooled from three independent experiments with similar trend, with four technical replicates *per* experiment. *Right panel* is a representative Giemsa-stained thin smear 24h after exposure to unconjugated bilirubin (41 μM) or vehicle. Circles in (C) represent technical replicates. *P* values determined using: (A, B, D) Two Way ANOVA with Tukey's multiple comparison test, (C) using Ordinary One-Way ANOVA with Tukey's multiple comparison test. NS: not significant; ***p<0.001, ****p<0.0001. Concentrations of unconjugated bilirubin in (A-D) were calculated according to the UnaG-based assay (*2, 3*) (*see Fig. S10*).

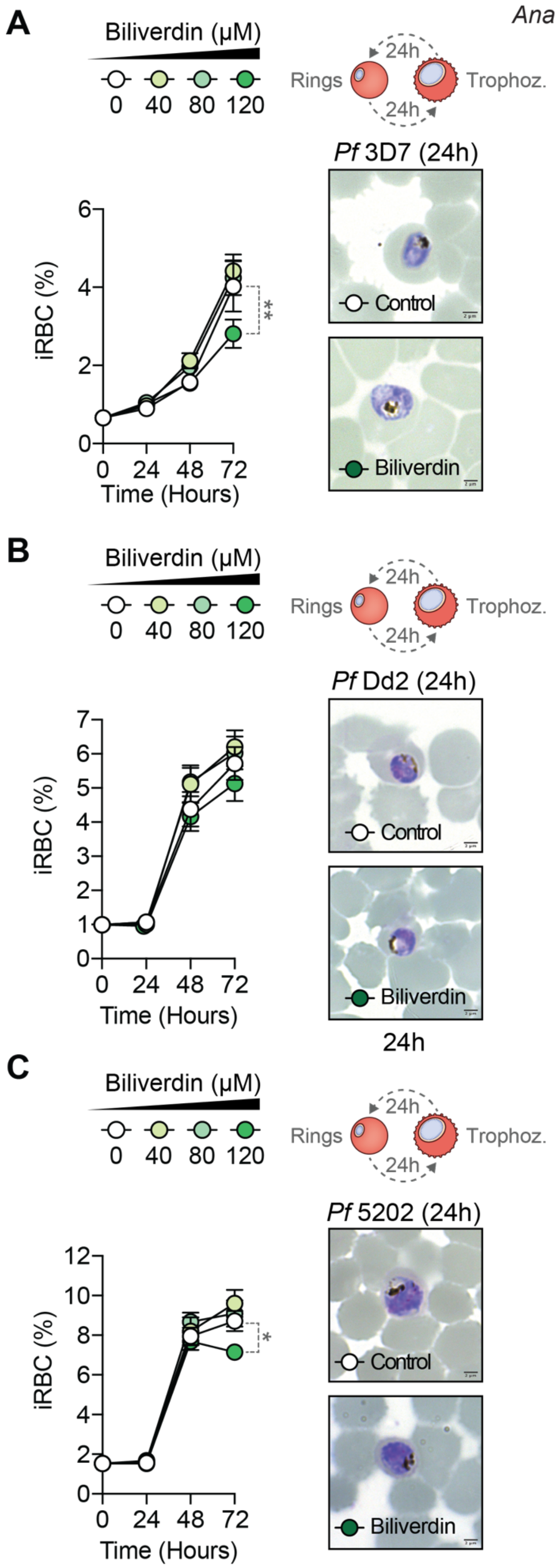
A
Biliverdin (μM)
0 40 80 120
Rings
Trophoz.
24h
24h
Pf 3D7 (24h)
iRBC (%)
Time (Hours)
Control
Biliverdin
B
Biliverdin (μM)
0 40 80 120
Rings
Trophoz.
24h
24h
Pf Dd2 (24h)
iRBC (%)
Time (Hours)
Control
Biliverdin
24h
C
Biliverdin (μM)
0 40 80 120
Rings
Trophoz.
24h
24h
Pf 5202 (24h)
iRBC (%)
Time (Hours)
Control
Biliverdin

**Fig. S12. Biliverdin does not arrest *P. falciparum* proliferation.** (**A**) Percentage (*left panel*) of *P. falciparum* 3D7 (*Pf*3D7) iRBC, 24-72h after exposure to increasing concentrations of biliverdin. *Right panel* is a representative Giemsa-stained thin smear 24h after exposure to biliverdin (120 μM) or vehicle (Control). (**B**) Percentage of *P. falciparum* Dd2 (*Pf*Dd2) iRBC (rings), 24-72h after exposure to increasing concentrations of biliverdin, as in (A). *Right panel* is a representative Giemsa-stained thin smear, 24h after exposure to biliverdin (120 μM) or vehicle (Control). (**C**) Percentage of *P. falciparum* IPC 5202 (*Pf*5202) iRBC (rings), 24-72h after exposure to increasing concentrations of biliverdin, as in (A). *Right panel* is a representative Giemsa-stained thin smear 24h after exposure to biliverdin (120 μM) or vehicle. Scale bars: 2μm. Data presented as mean ± SEM, pooled from six (A) or four (B, C) independent experiments with similar trend, with four technical replicates *per* experiment. *P* values determined using: (A-C) Two Way ANOVA with Tukey's multiple comparison test. *$p<0.05$, **$p<0.01$.

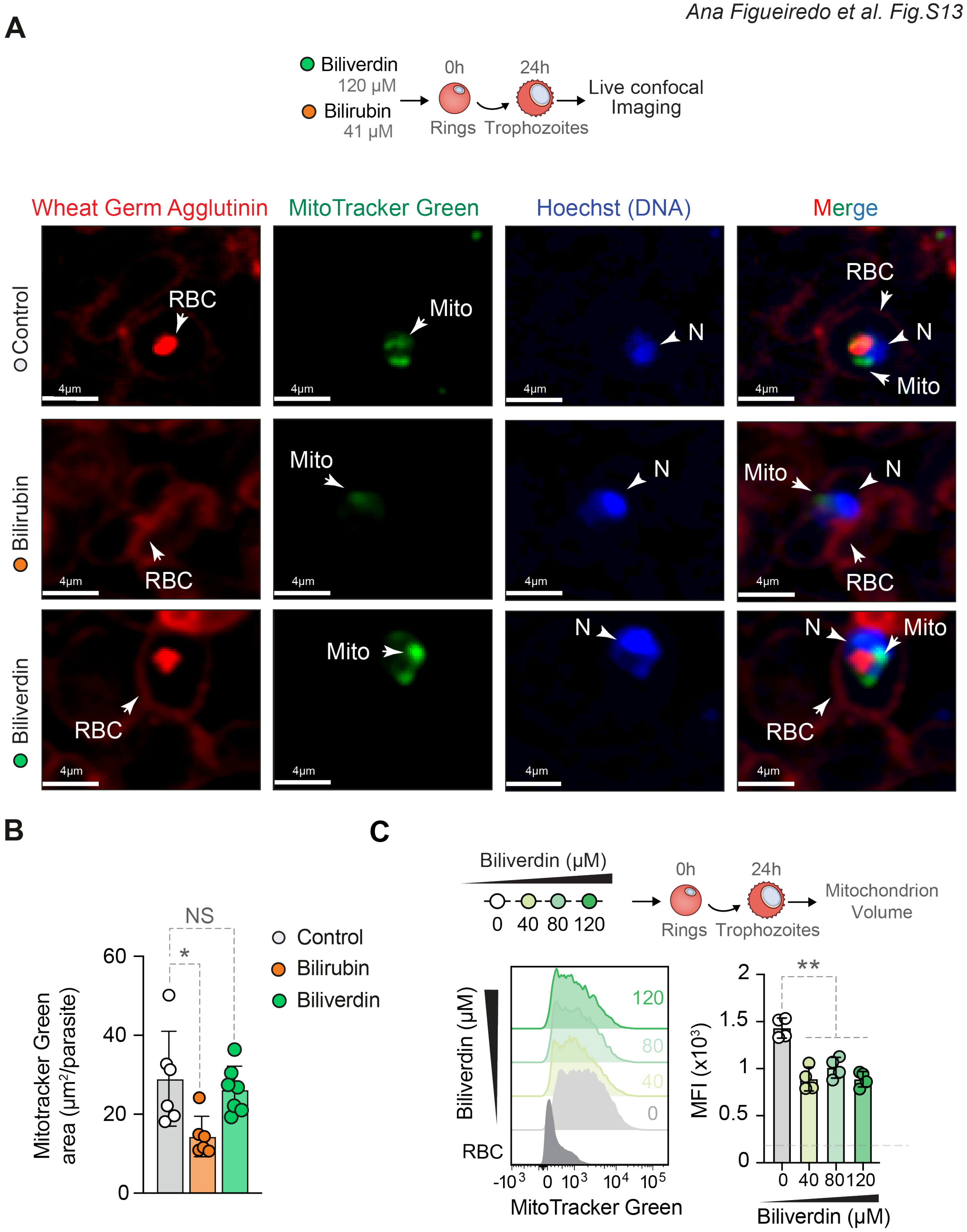
A
Biliverdin
120 µM
Bilirubin
41 µM
0h
24h
Rings
Trophozoites
Live confocal Imaging
Wheat Germ Agglutinin
MitoTracker Green
Hoechst (DNA)
Merge
Control
Bilirubin
Biliverdin
RBC
Mito
N
4µm
B
Mitotracker Green area (µm²/parasite)
NS
*
0
20
40
60
Control
Bilirubin
Biliverdin
C
Biliverdin (µM)
0 40 80 120
0h
24h
Rings
Trophozoites
Mitochondrion Volume
120
80
40
0
RBC
-10³ 0 10³ 10⁴ 10⁵
MitoTracker Green
MFI (x10³)
**
0
0.5
1
1.5
2
0 40 80 120
Biliverdin (µM)

**Fig. S13. Unconjugated bilirubin, but not biliverdin, disrupts *P. falciparum* mitochondrion integrity.** (**A**) Live confocal microscopy of *P. falciparum* (*Pf*3D7) iRBC (rings), 24h after exposure to unconjugated bilirubin (41 µM), biliverdin (120 µM) or vehicle (Control). Mitochondrion was stained using MitoTracker Green (green), RBC membranes using wheat germ agglutinin (WGA; red) and parasite´s DNA using Hoechst (blue). Scale bars: 4µm. (**B**) Quantification of the MitoTracker Green area *per* parasite after exposure to unconjugated bilirubin, biliverdin or vehicle (Control). Data represented as mean ± SD of 6-7 fields (n=30-34 parasites) from two independent experiments with similar trend. (**C**) Representative histograms (*left panel*) and quantification of median fluorescence intensity (MFI; *right panel*) of mitochondrion volume (MitoTracker Green), 24h after exposure of *Pf*3D7 iRBC (rings) to increasing concentrations of biliverdin. Grey histogram represents background staining in non-infected RBC and dashed grey (in bar plot) line represents the average background signal from all replicates in non-infected RBC. Data represented as mean ± SD from four replicates in one out of two independent experiments with similar trend. Circles in (B) represent individual parasites and in (D) technical replicates. *P* values determined using: (B, C) Ordinary One-Way ANOVA with Tukey's multiple comparison test. NS: not significant, $*p<0.05$, $**p<0.01$. Concentrations of unconjugated bilirubin in (A,B) were calculated according to the UnaG-based assay (*2, 3*) (*see Fig. S10*).

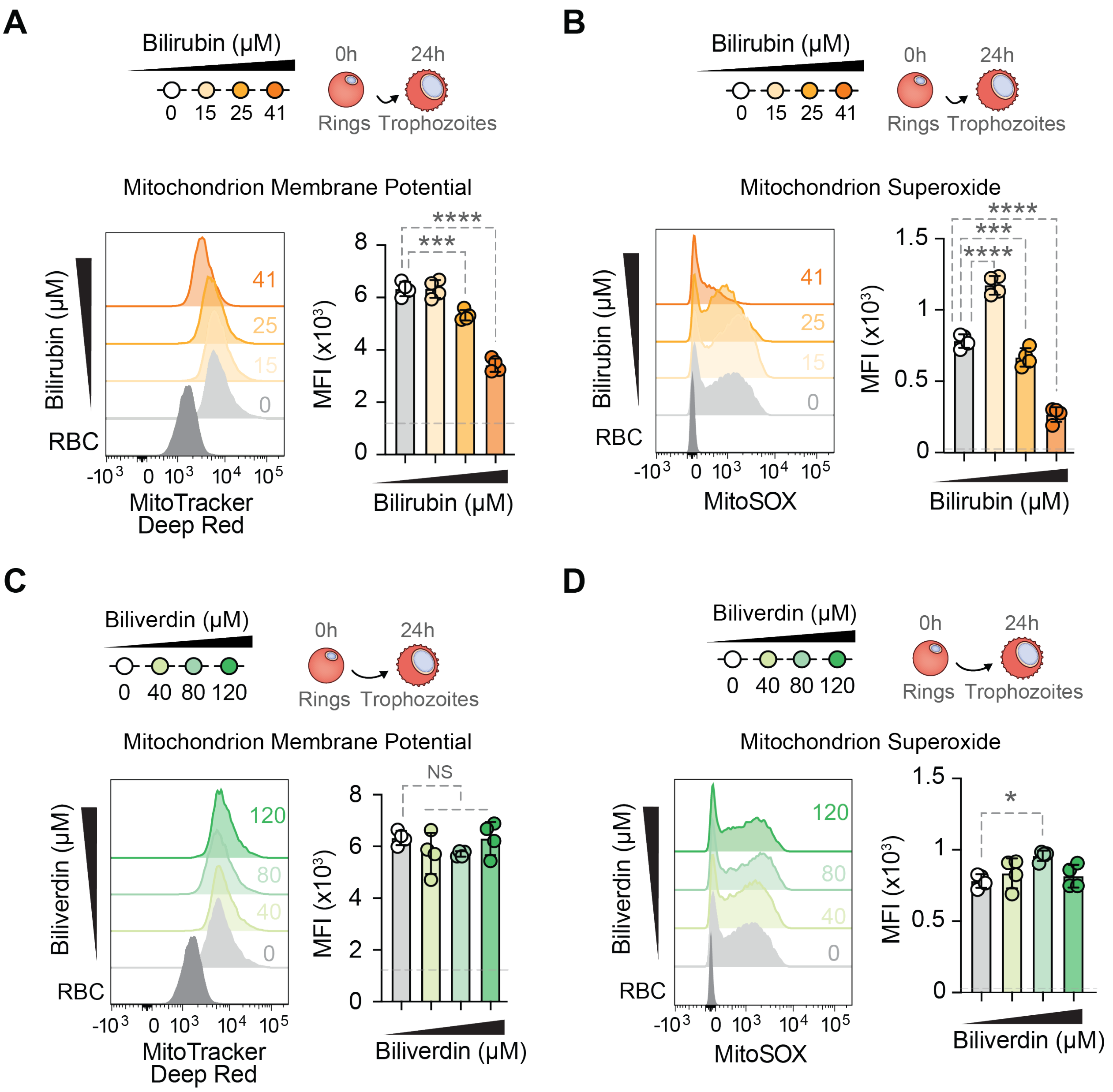
A
Bilirubin (μM)
0 15 25 41
0h 24h
Rings Trophozoites
Mitochondrion Membrane Potential
Bilirubin (μM)
RBC
MitoTracker Deep Red
MFI (x10³)
Bilirubin (μM)
B
Mitochondrion Superoxide
MitoSOX
C
Biliverdin (μM)
0 40 80 120
NS
D
*

**Fig. S14. Unconjugated bilirubin, but not biliverdin, disrupts *P. falciparum* mitochondrion function.** (**A**-**B**) Representative histograms (*left panel*) and quantification of median fluorescence intensity (MFI; *right panel*) of (**A**) mitochondrion membrane potential (MitoTracker Deep Red) and (**B**) mitochondrion superoxide production (MitoSOX), 24h after exposure of *Pf*3D7 iRBC (rings) to increasing concentrations of unconjugated bilirubin. Grey histogram represents background staining in non-iRBC and dashed grey lines (in bar plots) represent the average background signal from all replicates in non-infected RBC. Data represented as mean ± SD from four replicates in one out of two independent experiments with similar trend. (**C-D**) Representative histograms (*left panel*) and quantification of median fluorescence intensity (MFI; *right panel*) of (**C**) mitochondrion membrane potential (MitoTracker Deep Red) and (**D**) mitochondrion superoxide production (MitoSOX), 24h after exposure of *Pf*3D7 iRBC (rings) to increasing concentrations of biliverdin. Grey histogram represents background staining in non-infected RBC and dashed grey lines (bar plot) represent the average background signal from all replicates in non-infected RBC. Data represented as mean ± SD from four replicates in one out of two independent experiments with similar trend. Circles in *right panel* represent technical replicates. *P* values determined using: (A-D) Ordinary One-Way ANOVA with Tukey's multiple comparison test. NS: not significant; *p<0.05, ***p<0.001; ****p<0.0001. Concentrations of unconjugated bilirubin in (A,B) were calculated according to the UnaG-based assay (*2, 3*) (*see Fig. S10*).

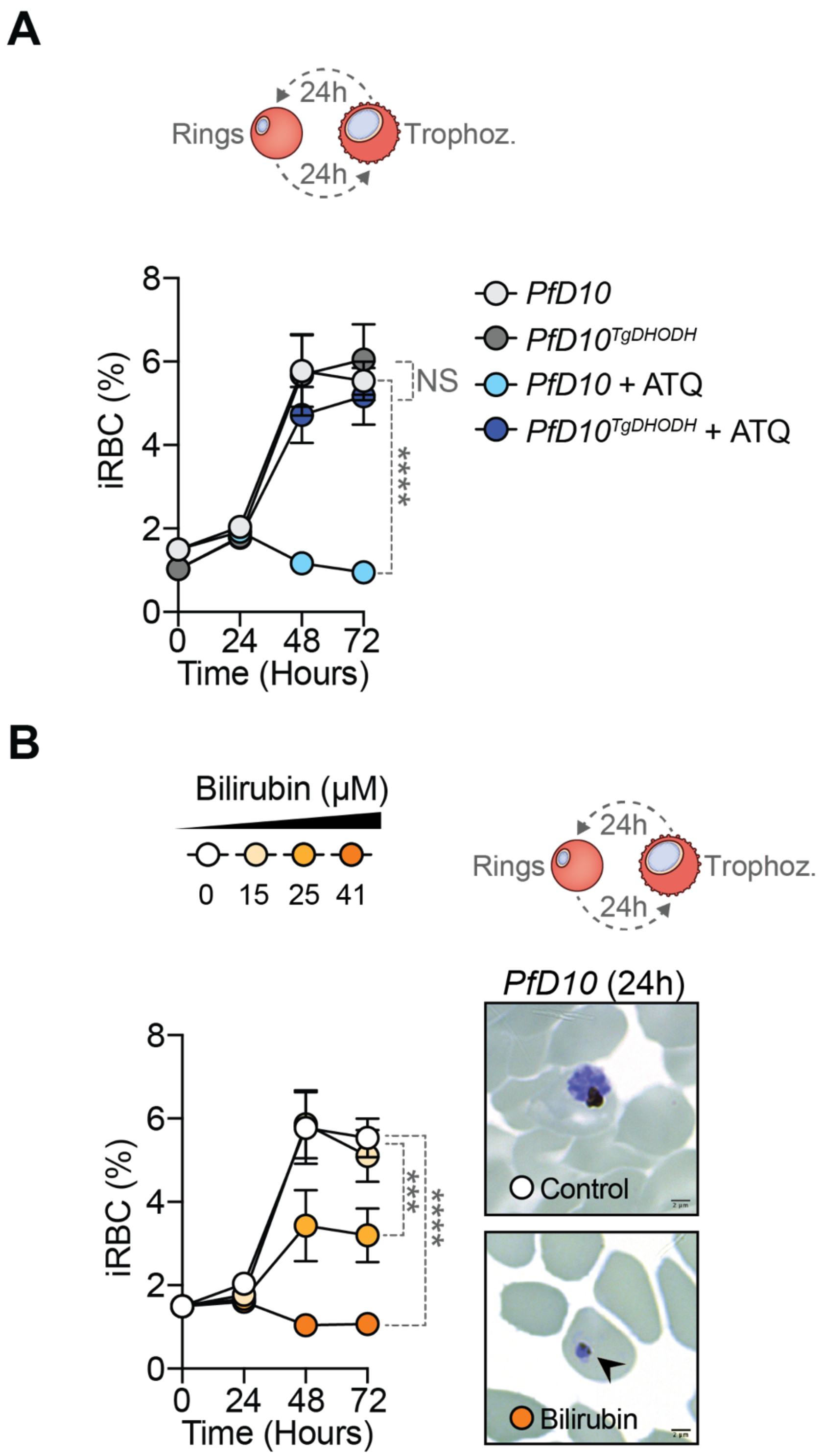
A
24h
Rings
Trophoz.
24h
iRBC (%)
8
6
4
2
0
0 24 48 72
Time (Hours)
NS
****
PfD10
PfD10TgDHODH
PfD10 + ATQ
PfD10TgDHODH + ATQ
B
Bilirubin (µM)
0 15 25 41
24h
Rings
Trophoz.
24h
PfD10 (24h)
iRBC (%)
8
6
4
2
0
0 24 48 72
Time (Hours)
***
****
Control
Bilirubin

**Fig. S15. Bilirubin inhibits the proliferation of atovaquone-resistant *P. falciparum* D10.** (**A**) Percentage of human iRBC with *P. falciparum* D10 (*Pf*D10, parental) or transgenic $PfD10^{TgDHODH}$ strain, expressing a cytoplasmic *Saccharomyces cerevisiae* DHODH (rings), 24-72h after exposure to atovaquone (10nM). (**B**) Percentage of *Pf*D10 iRBC (*left panel*), 24-72h after exposure to increasing concentrations of unconjugated bilirubin. Concentrations of unconjugated bilirubin in were calculated according to the UnaG-based assay (*2, 3*) (*see Fig. S10*). Data represented as mean ± SEM, pooled from 3 independent experiments with similar trend, with four biological replicates *per* experiment. *Right panel* in (B) is representative Giemsa-stained thin smears, 24h after exposure to unconjugated bilirubin (41 µM) or vehicle (Control). Scale bars: 2µm. Black arrowhead highlights pyknotic parasites. *P* values determined using: (A, B) Two-Way ANOVA with Tukey's multiple comparison test. NS: not significant; ***p<0.001; ****p<0.0001.

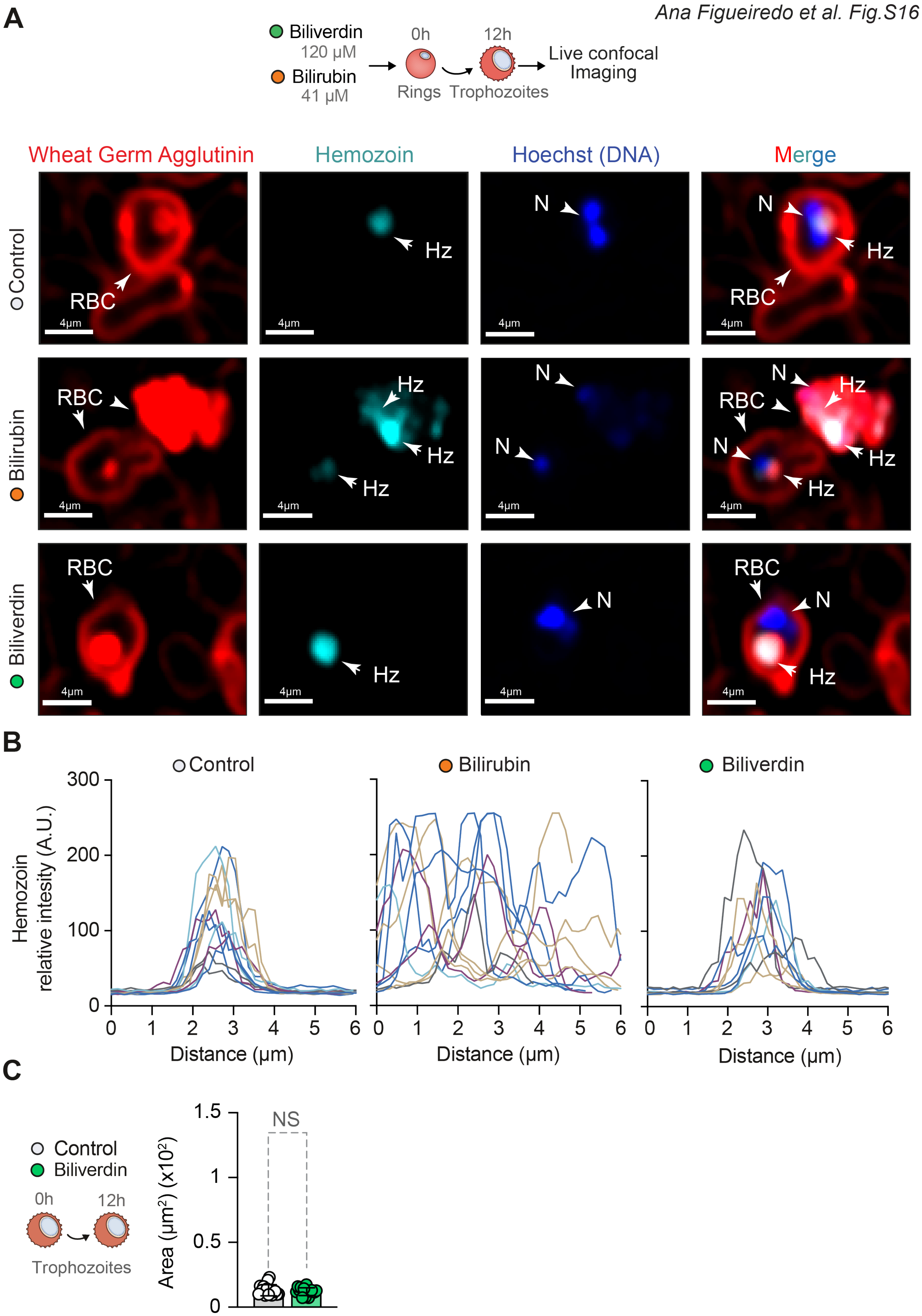
A
Biliverdin
120 µM
Bilirubin
41 µM
0h
12h
Rings
Trophozoites
Live confocal
Imaging
Wheat Germ Agglutinin
Hemozoin
Hoechst (DNA)
Merge
Control
Bilirubin
Biliverdin
RBC
Hz
N
4µm
B
Control
Bilirubin
Biliverdin
Hemozoin
relative intesity (A.U.)
300
200
100
0
0 1 2 3 4 5 6
Distance (µm)
C
Control
Biliverdin
0h
12h
Trophozoites
Area (µm²) (x10²)
1.5
1
0.5
0
NS

**Fig. S16. Bilirubin, but not biliverdin, causes Hz crystal spillover in *P. falciparum* trophozoites.** (**A**) Live confocal microscopy of *P. falciparum* (*Pf*3D7) iRBC (trophozoites), 12h after exposure to unconjugated bilirubin (41 µM), biliverdin (120 µM) or vehicle (Control). RBC membranes were stained by wheat germ agglutinin (RBC; red), using the laser reflection mode (cyan) and parasite´s DNA using Hoechst (DNA: blue). Scale bars: 4µm. (**B**) Quantification of the relative intensity of Hz fluorescence individual traces, obtained from the cross-section of 10-15 *Pf*3D7 iRBC, treated as in (A). (**C**) Total area and intensity distribution of Hz in *Pf*3D7 parasites exposed to biliverdin as in (A). Data represented as mean ± SD, pooled from two independent experiments (n=10-15 parasites). Circles in (C) represent individual parasites. *P* values determined using: (C) Mann-Whitney U test. NS: not significant. Concentrations of unconjugated bilirubin in (A,B) were calculated according to the UnaG-based assay (*2, 3*) (*see Fig. S10*).

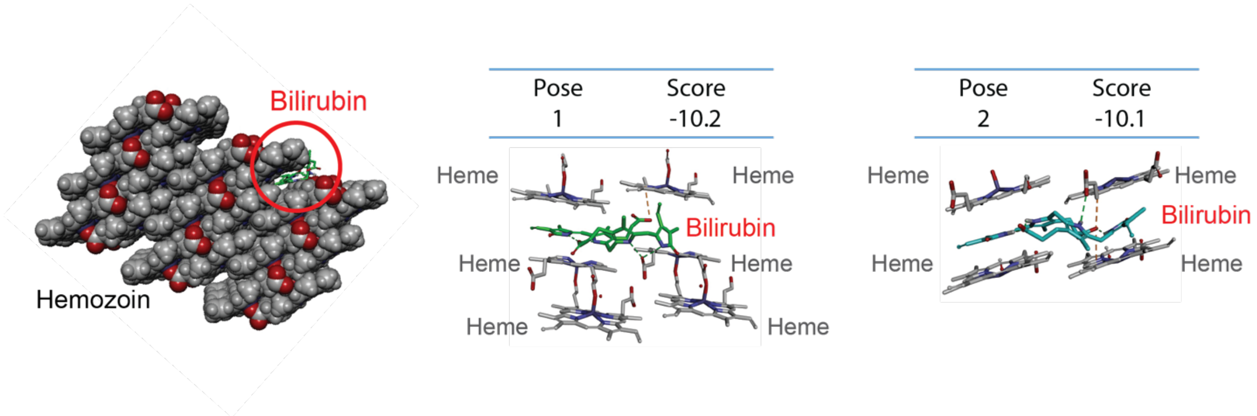


**Fig. S17. Bilirubin interaction with Hz crystals.** Predicted interaction of unconjugated bilirubin with Hz, whereby unconjugated bilirubin docks in the deep groove of the 001 fastest growing face of the β-hematin crystal (*40*). The two top scoring docking poses suggest that unconjugated bilirubin can form intramolecular hydrogen bonds between carboxylic acid moieties and pyrrole groups of heme, in β-hematin.

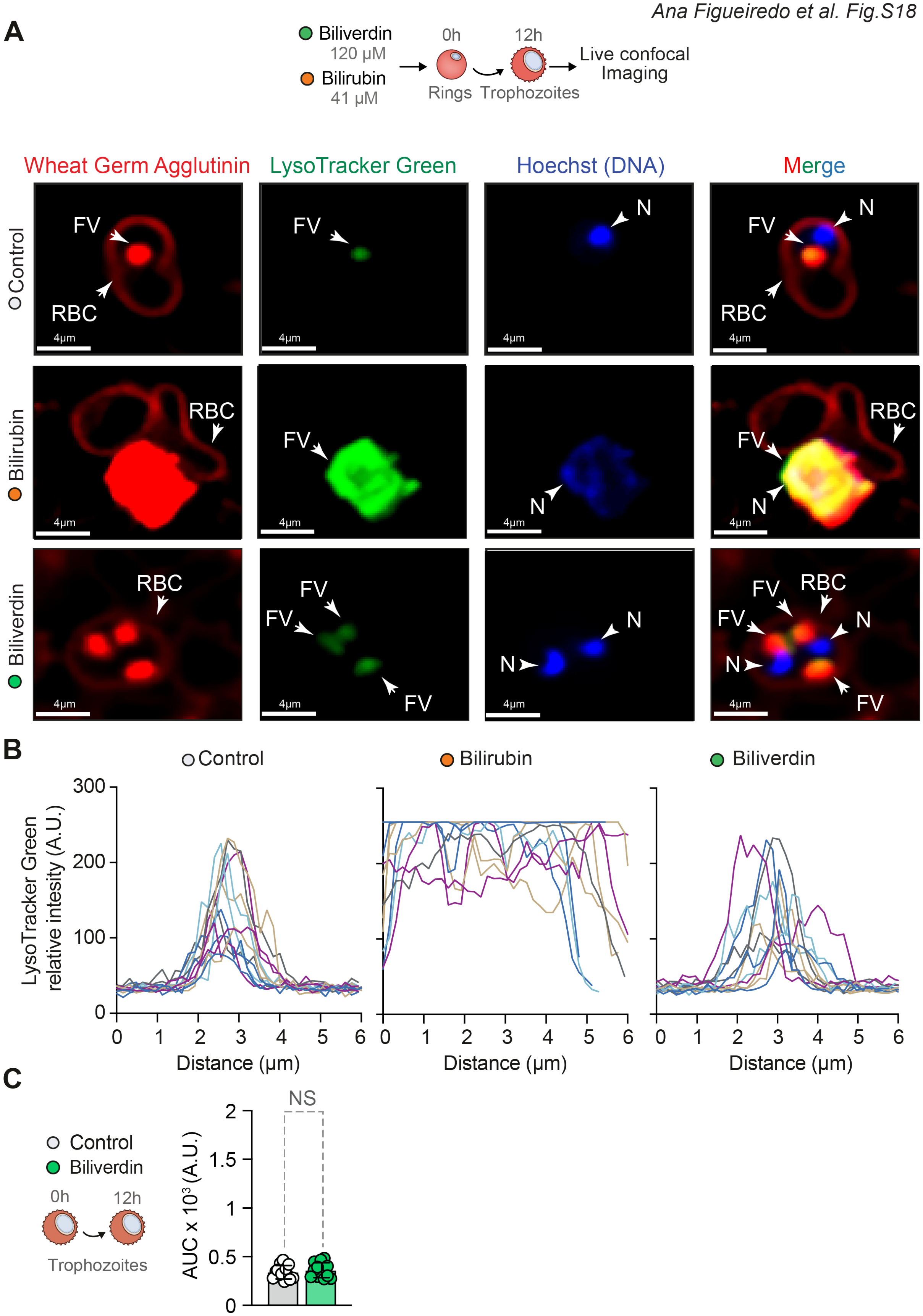
A
Biliverdin 120 µM
Bilirubin 41 µM
0h
12h
Rings
Trophozoites
Live confocal Imaging
Wheat Germ Agglutinin
LysoTracker Green
Hoechst (DNA)
Merge
Control
Bilirubin
Biliverdin
FV
RBC
N
4µm
B
Control
Bilirubin
Biliverdin
LysoTracker Green relative intesity (A.U.)
Distance (µm)
C
Control
Biliverdin
0h
12h
Trophozoites
AUC x 10³ (A.U.)
NS

**Fig. S18. Bilirubin, but not biliverdin, disrupts *P. falciparum* food vacuole.** (**A**) Live confocal microscopy of *P. falciparum* (*Pf*3D7) iRBC (trophozoites), 12h after exposure to unconjugated bilirubin (41 µM), biliverdin (120 µM) or vehicle (Control). RBC membranes were stained using wheat germ agglutinin (RBC; red), parasite´s food vacuole using LysoTracker Green (Food vacuole; FV; green) and the parasite´s DNA using Hoechst (DNA: blue). Scale bars: 4µm. (**B**) Quantification of the relative intensity of LysoTracker fluorescence individual traces, obtained from the cross-section of 10-15 *Pf*3D7 iRBC, treated as in (A). (**C**) Area under the curve (AUC) of lysotracker intensity distribution in parasites exposed to biliverdin as in (A). Data represented as mean ± SD, pooled from two independent experiments (n=10-15 parasites). Circles in (C) represent individual parasites. *P* values determined using: (C) Mann-Whitney U test. NS: not significant. Concentrations of unconjugated bilirubin in (A,B) were calculated according to the UnaG-based assay (*2, 3*) (*see Fig. S10*).

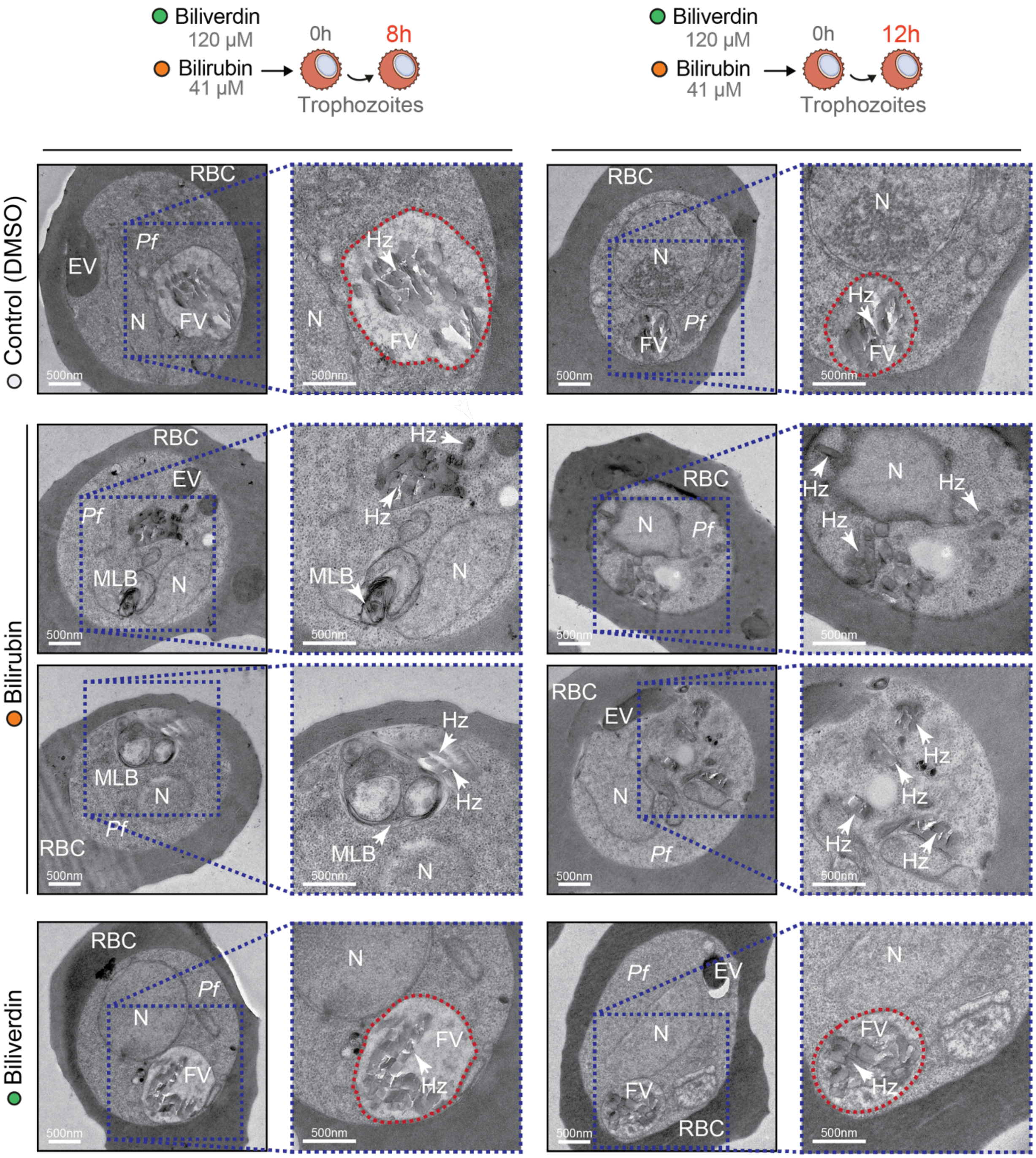
Biliverdin
120 µM
Bilirubin
41 µM
0h
8h
Trophozoites
Biliverdin
120 µM
Bilirubin
41 µM
0h
12h
Trophozoites
Control (DMSO)
Bilirubin
Biliverdin
RBC
EV
Pf
N
FV
Hz
MLB
500nm

**Fig. S19. Bilirubin, but not biliverdin, disrupts *P. falciparum* food vacuole and induces the formation of multilamellar bodies.** Representative transmission electron microscopy images of *P. falciparum* (*3D7*) iRBC. Images show trophozoites, exposed to unconjugated bilirubin (41 μM; 8h or 12h) or vehicle (Control) or biliverdin (120 μM; 8h or 12h). Concentration of unconjugated bilirubin was calculated according to the UnaG-based assay (*2, 3*) (*see Fig. S10*). Images in *right panel* correspond to amplifications of the blue dotted area highlighted in the *left panel*. Dotted red lines highlight trophozoite´s food vacuole (FV), not visible in parasites exposed to unconjugated bilirubin. Notice the "rounded-shaped" Hz crystals in the cytoplasm of trophozoites exposed to unconjugated bilirubin, compared to the sharp-edged Hz crystals in control trophozoites exposed to vehicle or biliverdin. EV: Endocytic vesicle, FV: Food vacuole, Hz: Hemozoin, MLB: Multilamellar bodies, N: Nucleus, RBC: Red blood cell. Images are representative of three independent experiments with similar results. Scale bar: 500 nm.

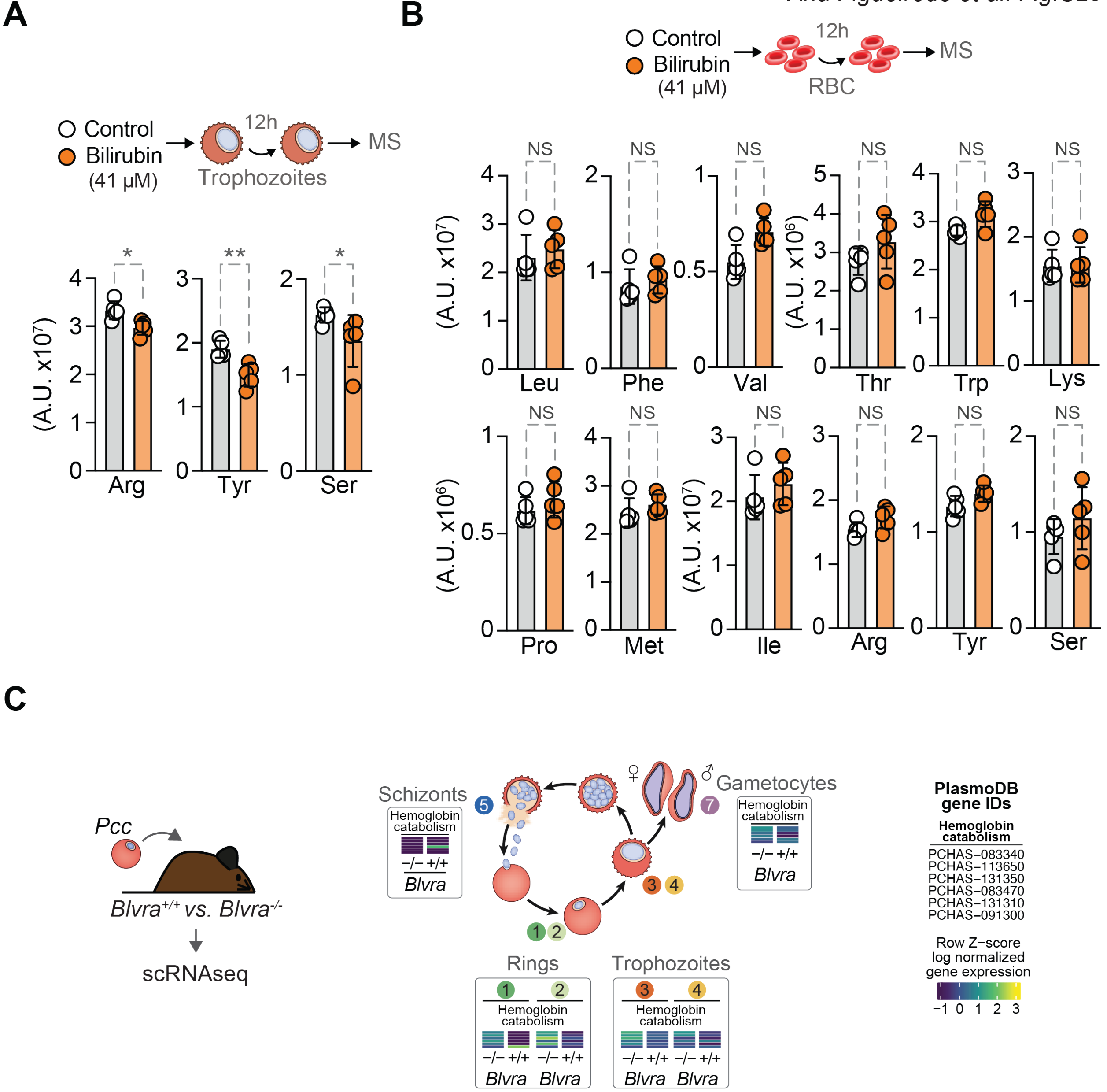


**Fig. S20. Bilirubin inhibits the capacity of *P. falciparum* to extract essential AA contained in hemoglobin.** (**A**) Relative levels of essential AA Arginine (Arg), Tyrosine (Tyr) and Serine (Ser) extracted from hemoglobin in *Pf*3D7 iRBC (trophozoites), 12h after exposure to unconjugated bilirubin (41 µM) or vehicle (Control), detected by Mass Spectrometry. (**B**) Relative levels of AA Leucine (Leu), Phenylalanine (Phe), Valine (Val), Threonine (Thr), Tryptophan (Trp), Lysine (Lys), Proline (Pro), Methionine (Met) Isoleucine (Ile), Arginine (Arg), Tyrosine (Tyr) and serine (Ser) in non-infected RBC, detected by Mass Spectrometry 12h after exposure to unconjugated bilirubin (41 µM) or vehicle (Control), as in (A). (**C**) Schematic representation of differentially expressed genes involved in hemoglobin catabolism in *P. chabaudi chabaudi* iRBC from *Blvra*$^{-/-}$ vs. *Blvra*$^{+/+}$ mice, as determined by scRNA sequencing, in the different Populations identified in (Figure 2A-E, Table S2). Circles in (A, B) represent technical replicates. *P* values determined using: (A, B) using determined using Mann-Whitney U test. NS: not significant; *p<0.05, **p<0.01. Concentrations of unconjugated bilirubin in (A,B) were calculated according to the UnaG-based assay (*2, 3*) (*see Fig. S10*).

#### Other Supplementary Material

##### Table S1 (separate Excel file).

Conserved transcripts *per* population in *Blvra*$^{+/+}$ and *Blvra*$^{-/-}$ mice.

##### Table S2 (separate Excel file).

Differentially expressed genes of parasites from *Blvra*$^{+/+}$ and *Blvra*$^{-/-}$ mice.

##### Table S3 (separate Excel file).

Pathway analysis of differentially expressed genes of parasites from *Blvra*$^{+/+}$ and *Blvra*$^{-/-}$ mice.

##### Table S4 (separate Excel file).

Shared differentially expressed genes related to parasite virulence of parasites from *Blvra*$^{+/+}$ and *Blvra*$^{-/-}$ mice.

##### Table S5 (separate Excel file).

Metabolites detected in the metabolomic analysis in negative and positive modes of *Pf*3D7 iRBC (trophozoites) or non-infected RBC exposed to unconjugated bilirubin (41 μM; 8h or 12h) or vehicle.

##### Data S1 (separate PDF file).

Gating strategy for: A) flow cytometry analysis of *Pcc*-infected red blood cells, related to Figure 2A,B,G,H and Fig.S5A-D) for flow cytometry analysis of *Pf* iRBC, related to Figure 3B, G, Fig.S11A,B,D, Fig.S12A-C, and Fig.S15A,B. Red blood cells were identified according to size (FSC-A) *vs.* granularity (SSC-A) plots. Cell doublets were excluded according to cellular area (FSC-A) *vs.* height (FSC-H), selecting for single cells. *Pcc*-infected red blood cells were identified based on GFP, SYBR Green or SYTO Deep Red positive fluorescence.

##### Data S2 (separate PDF file).

Gating strategy for flow cytometry analysis of *Pf*-iRBC, related to Figure 3E, Fig.S13C, and Fig.S14A-D. Red blood cells were identified in size (FSC-A) *vs.* granularity (SSC-A) plots. Cell doublets were excluded according to cellular area (FSC-A) *vs.* height (FSC-H) selecting for single cells. *Pcc*-infected red blood cells were identified based on Hoechst positive fluorescence. MitoTracker Green, MitoTracker Deep Red and MitoSOX mean fluorescence intensities (MFIs) were assessed in the *Pcc*-infected red blood cells.

##### Data S3 (separate PDF file).

Uncropped Western blot membranes, related to Fig.S2B and Fig.S4D, E.

*Supporting data for Figure 2A,B,G,H and Fig.S5A-D, Gating strategy*

**A**

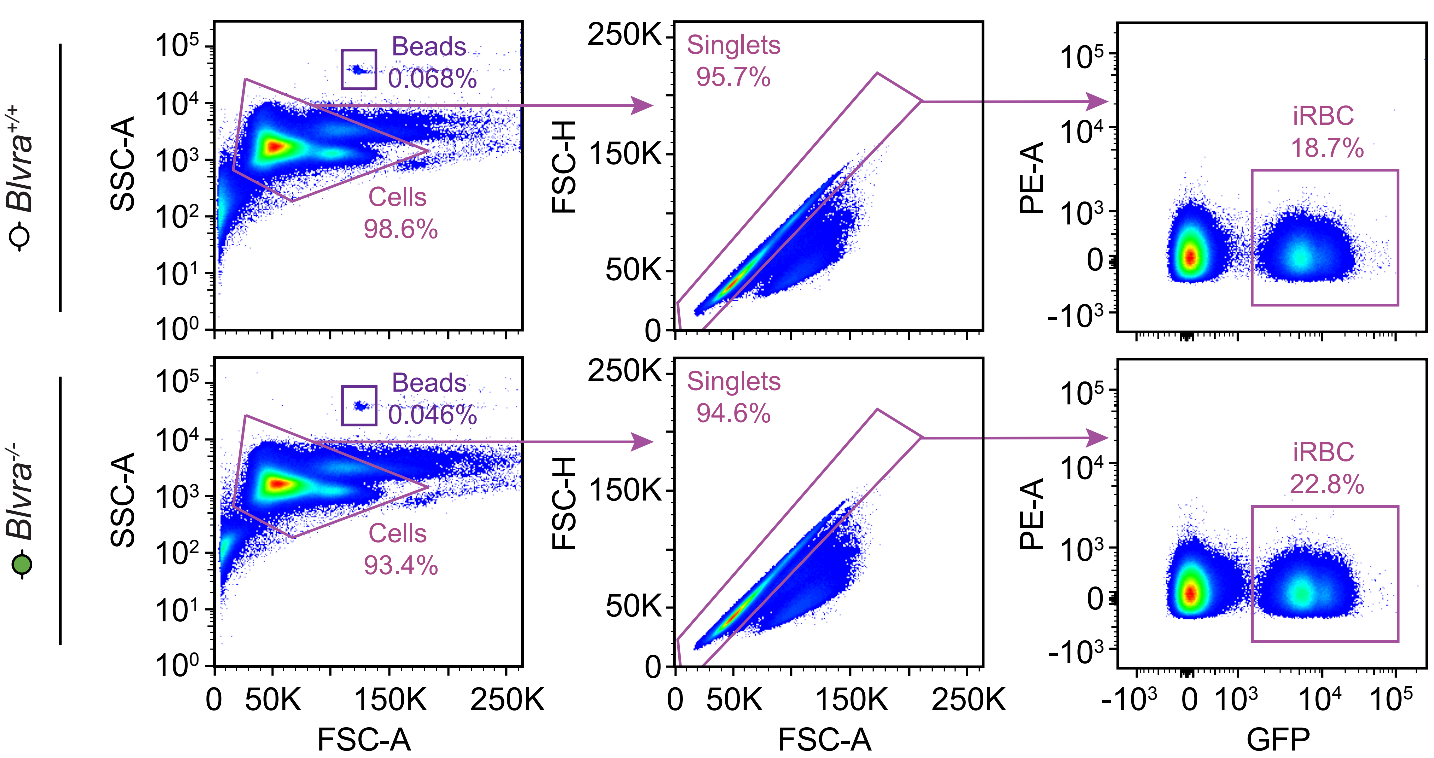


*Supporting data for Figure 3B,G, Fig.S11A,B,D, Fig.S12A-C, and Fig.S15A,B, Gating strategy*

**B**

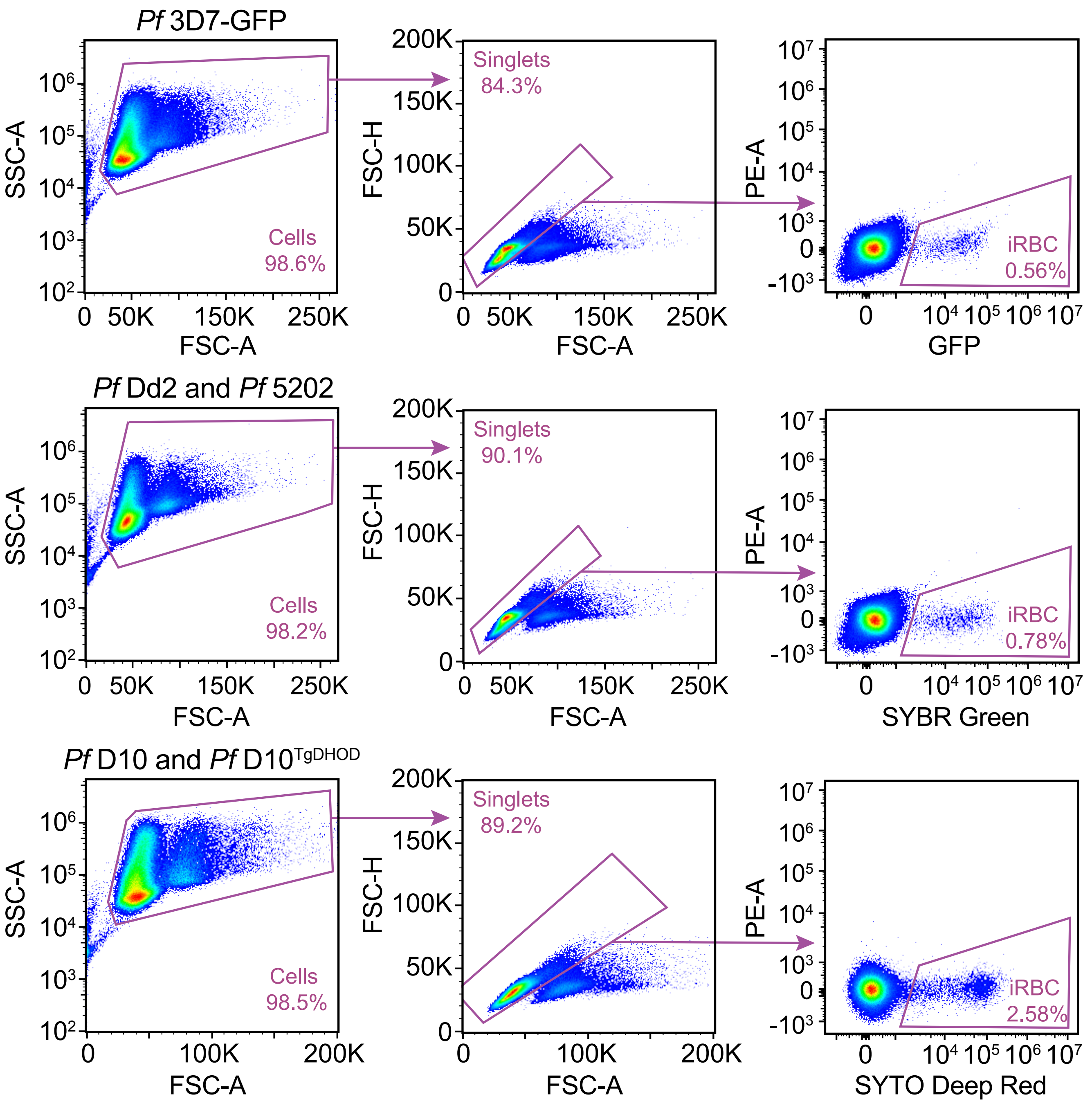

*Supporting data for Figure 3E, Fig.S13C, and Fig.S14A-D, Gating strategy*

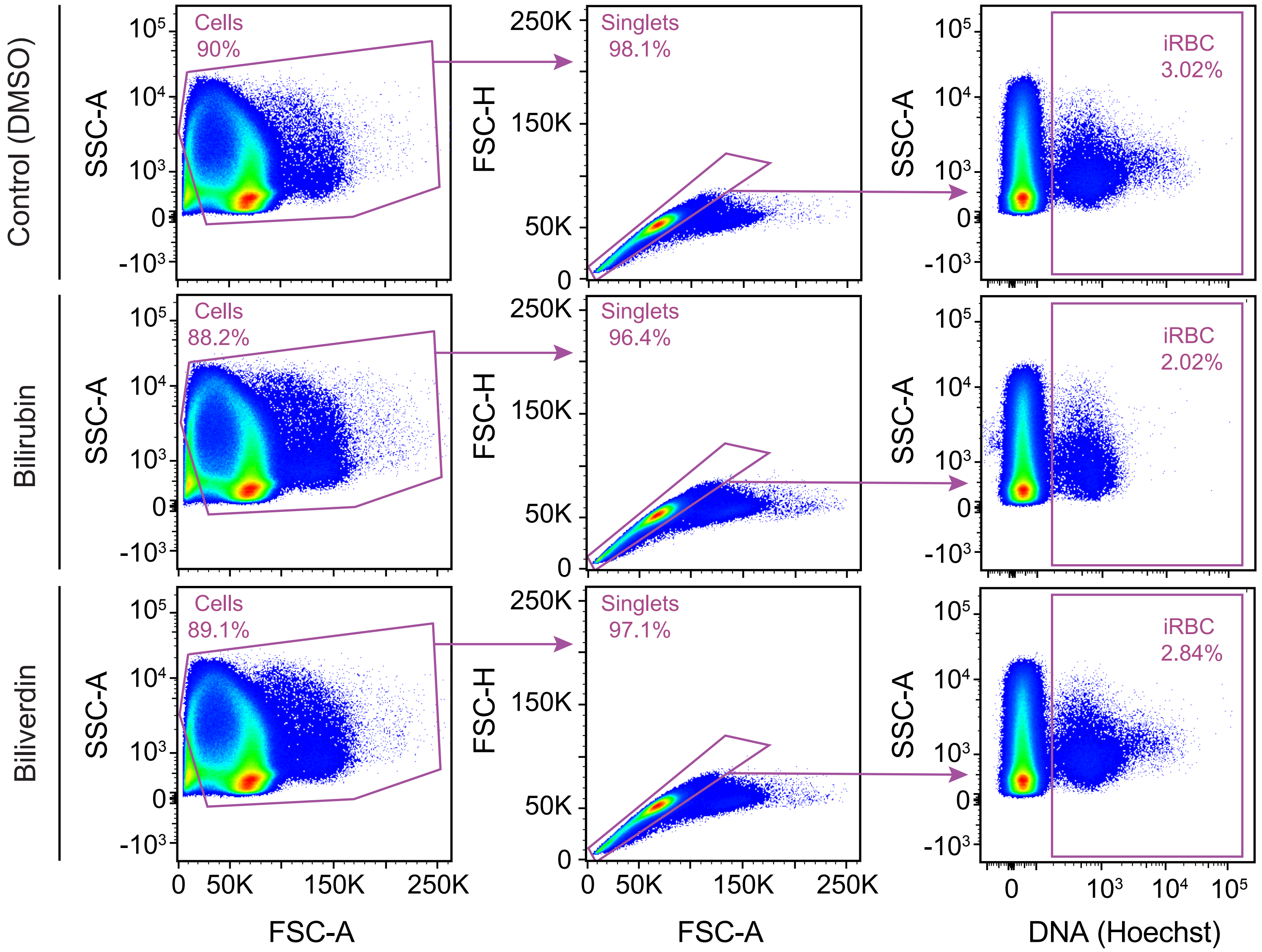

#### Supporting data for Fig.S2B, Wester Blot

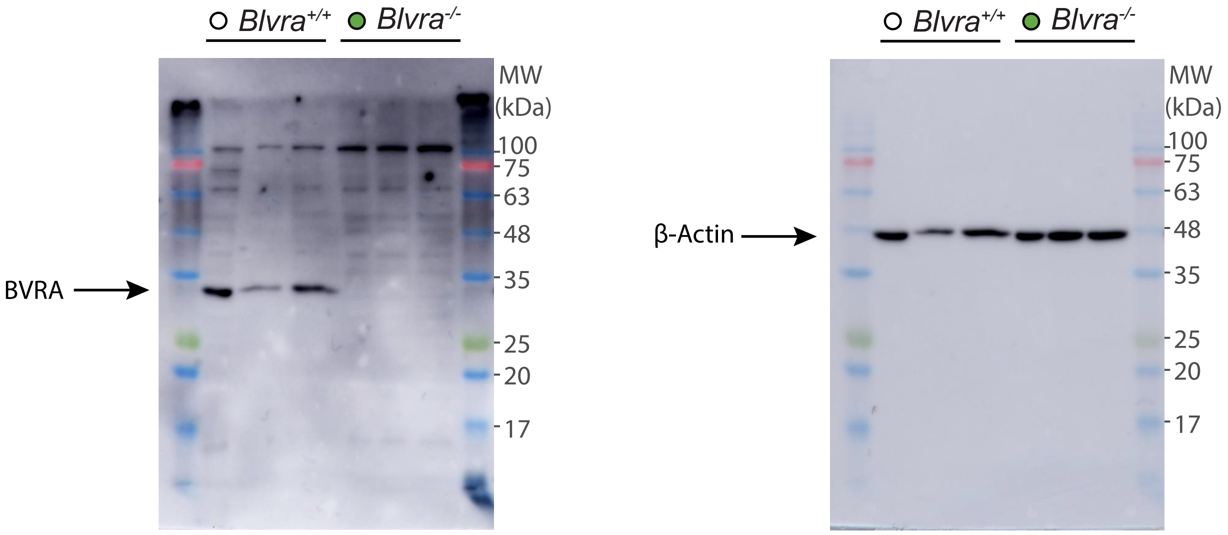


Lanes correspond to individual mice

#### Supporting data for Fig.S4D, Wester Blot

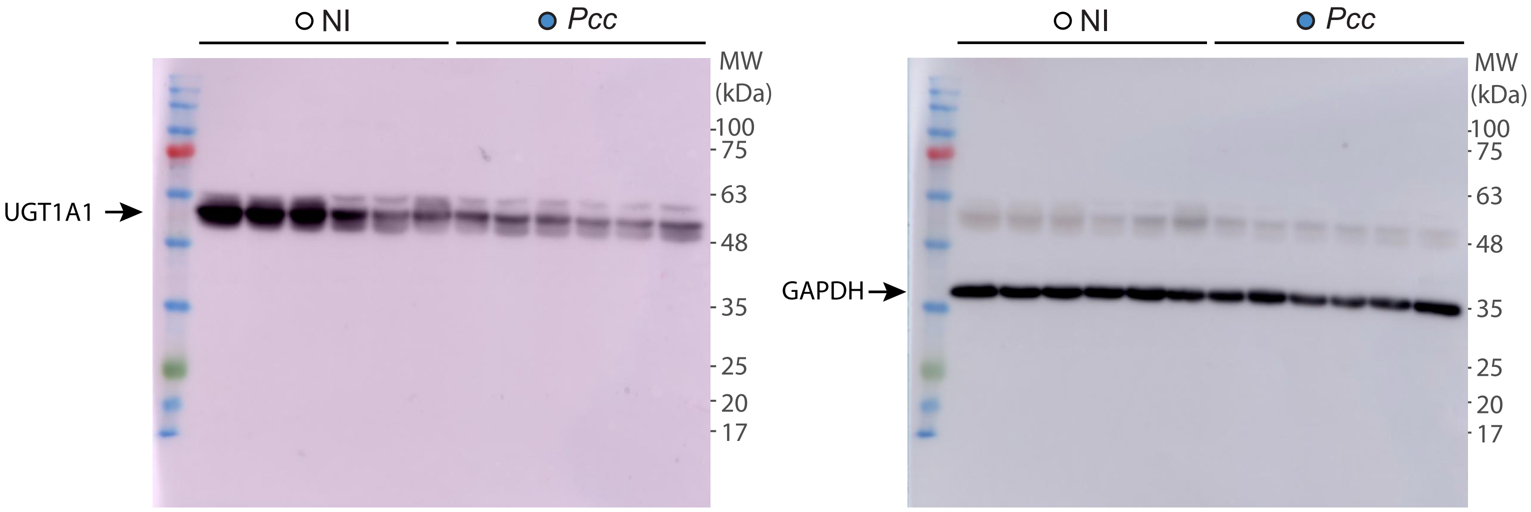


Lanes correspond to individual mice

#### Supporting data for Fig.S4E, Wester Blot

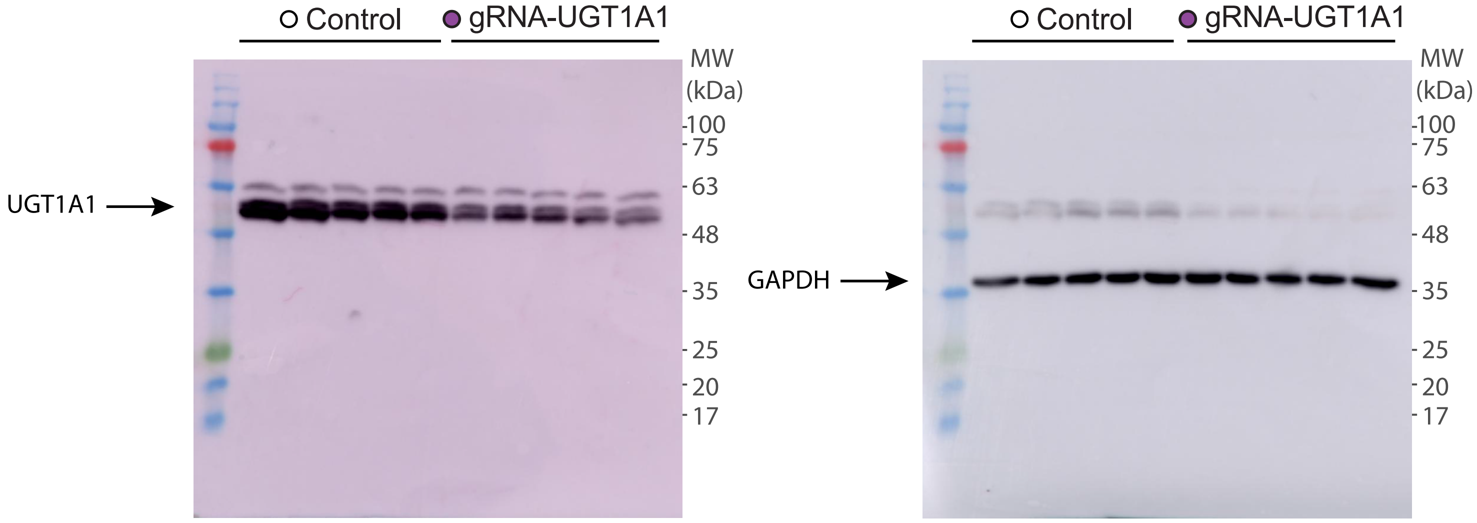


Lanes correspond to individual mice
